# Subfunctionalization of paralog transcription factors contributes to regulation of alkaloid pathway branch choice in *Catharanthus roseus*

**DOI:** 10.1101/2020.05.04.075671

**Authors:** Maite Colinas, Jacob Pollier, Dries Vaneechoutte, Deniz G. Malat, Fabian Schweizer, Liesbeth De Milde, Rebecca De Clercq, Joana G. Guedes, Teresa Martínez-Cortés, Francisco J. Molina Hidalgo, Mariana Sottomayor, Klaas Vandepoele, Alain Goossens

## Abstract

*Catharanthus roseus* produces a diverse range of specialized metabolites of the monoterpenoid indole alkaloid (MIA) class in a heavily branched pathway. Recent great progress in identification of MIA biosynthesis genes revealed that the different pathway branch genes are expressed in a highly cell type- and organ-specific and stress-dependent manner. This implies a complex control by specific transcription factors (TFs), only partly revealed today. We generated and mined a comprehensive compendium of publicly available *C. roseus* transcriptome data for MIA pathway branch-specific TFs. Functional analysis was performed through extensive comparative gene expression analysis and profiling of over 40 MIA metabolites in the *C. roseus* flower petal expression system. We identified additional members of the known BIS and ORCA regulators. Further detailed study of the ORCA TFs suggests subfunctionalization of ORCA paralogs in terms of target gene-specific regulation and synergistic activity with the central jasmonate response regulator MYC2. Moreover, we identified specific amino acid residues within the ORCA DNA-binding domains that contribute to the differential regulation of some MIA pathway branches. Our results advance our understanding of TF paralog specificity for which, despite the common occurrence of closely related paralogs in many species, comparative studies are scarce.

**SIGNIFICANCE STATEMENT:** A gene discovery program for regulators of monoterpenoid indole alkaloid biosynthesis in *Catharanthus roseus* advances our understanding of paralog specificity and subfunctionalization of the renowned class of ORCA transcription factors, particularly in terms of target gene-specificity and synergistic activity with other jasmonate-responsive transcription factors.

## INTRODUCTION

Plant specialized metabolites are chemically diverse, typically species- or taxa-specific and have often evolved in adaptation to the ecological niche of the plant. Within the plant, their production often exclusively takes place in specific organs or cell types and may be up-regulated precisely in response to defined environmental conditions [for a review, see Colinas and Goossens (2018)]. The integration of internal cues, such as cell type environment and developmental stage, and external cues, such as the presence of pathogens or herbivores, is predominantly regulated at the transcriptional level by specific transcription factors (TFs). Different TFs might act independently or cooperatively, forming regulatory modules that ensure the optimal spatiotemporal expression of specific metabolic pathway genes.

The medicinal plant *Catharanthus roseus* is well-known as being the only source of the important anti-cancer compounds vinblastine and vincristine (van der Heijden *et al.*, 2004). These complex molecules belong to the class of monoterpenoid indole alkaloids (MIAs); around 150 different MIAs are estimated to occur in *C. roseus* (van der Heijden *et al.*, 2004). The identification of many of the involved biosynthetic enzymes has revealed that genes involved in the different MIA pathway steps and branches are expressed in a cell type-specific, organ-specific and stress-dependent manner, making this species an ideal model to study how a modular system of different TFs possibly regulates specialized metabolism (Courdavault *et al.*, 2014; Dugé de Bernonville *et al.*, 2015).

Different MIA pathway steps and branches can be organized into *subdivisions* with connected gene expression patterns (Fig. 1). The first of such *subdivisions* is the *iridoid pathway*, i.e. the biosynthesis of the precursor loganic acid from geranyl diphosphate (GPP) via seven enzymatic steps occurring in the specialized internal phloem-associated parenchyma (IPAP) cells (Geu-Flores *et al.*, 2012; Asada *et al.*, 2013; Simkin *et al.*, 2013; Miettinen *et al.*, 2014; Salim *et al.*, 2014). An iridoid intermediate, presumably loganic acid, is then transported to the epidermis, possibly by the recently characterized nitrate/peptide family (NPF) transporters (NPF2.4, NPF2.5, and NPF2.6) (Yamamoto *et al.*, 2016; Larsen *et al.*, 2017; Yamamoto *et al.*, 2019). The coupling of the then converted iridoid secologanin to the tryptophan-derived indole moiety tryptamine marks the formation of the first MIA strictosidine, from which, after deglycosylation, all MIAs derive (*strictosidine pathway*). A single enzymatic step is needed to convert the strictosidine aglycone either into the heterohymbines ajmalicine or tetrahydroalstonine (and subsequently alstonine) (Stavrinides *et al.*, 2016; Dang *et al.*, 2018), or into vitrosamine (Stavrinides *et al.*, 2018). The branch leading to the precursors of the anti-cancer compounds has recently been resolved and involves seven enzymatic steps to yield the unstable intermediate dehydrosecodine (named *stemmadenine pathway* according to the name of one of the intermediates) (Tatsis *et al.*, 2017; Caputi *et al.*, 2018; Qu *et al.*, 2018; Qu *et al.*, 2019). From the latter either tabersonine or catharanthine are formed by two different enzymes (Caputi *et al.*, 2018; Qu *et al.*, 2018). Catharanthine forms one moiety of the MIA dimers vinblastine and vincristine and has been additionally suggested to have a repellent function as a monomer after secretion to the leaf surface by an epidermal ATP-binding cassette (ABC) transporter (Roepke *et al.*, 2010; Yu and De Luca, 2013). The second monomer, vindoline, is produced from tabersonine via seven enzymatic steps, from which the two last steps are thought to take place in specialized idioblasts or laticifers (St-Pierre and De Luca, 1995; St-Pierre *et al.*, 1999; Levac *et al.*, 2008; Liscombe *et al.*, 2010; Guirimand *et al.*, 2011; Liscombe and O’Connor, 2011; Besseau *et al.*, 2013; Qu *et al.*, 2015). Noteworthy, the *vindoline pathway* exclusively occurs in the green aerial parts of the plant (Liu *et al.*, 2019). By contrast, in roots (particularly hairy roots), a different set of interconvertible *root-specific MIAs* (lochnericine, hörhammericine, minovincinine among others) is formed (Laflamme *et al.*, 2001; Giddings *et al.*, 2011; Carqueijeiro *et al.*, 2018a; Carqueijeiro *et al.*, 2018b; Williams *et al.*, 2019). Finally, the assumed spatially separated monomers vindoline and catharanthine are dimerized to α-3’,4’-anhydrovinblastine, the direct precursor of vinblastine and vincristine, presumably by a peroxidase (Costa *et al.*, 2008). Noteworthy, a very recent study suggests that catharanthine, as well as ajmalicine and its oxidation product serpentine, may also accumulate in idioblasts and laticifers in addition to the epidermis (Yamamoto *et al.*, 2019). Thus, more research is needed in order to clarify the hypothesis of spatial separation of the monomers and differences in the sites of biosynthesis and accumulation of MIA monomers.

**Fig. 1.**
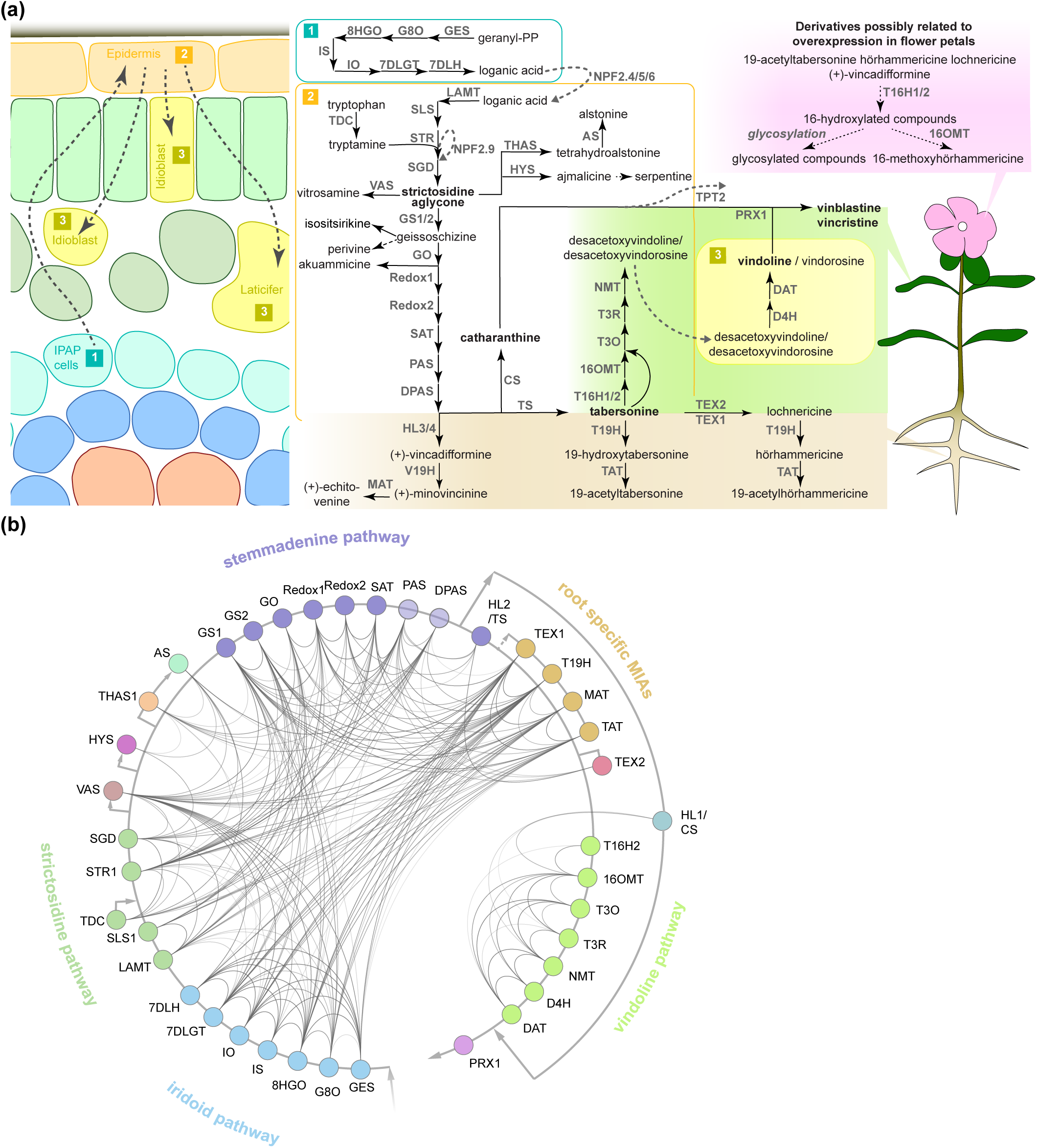
The MIA pathway consists of different branches with corresponding co-expression networks. (a) The MIA network and known or presumed cellular and organ-specific expression of the encoding MIA biosynthesis genes based on the current knowledge and available data. MIA biosynthesis is compartmentalized into the three different cell types shown on the left, although a cell type-specific expression has not been experimentally verified for all MIA biosynthesis genes such as the root MIA biosynthesis genes. Available organ-specific transcriptome data revealed that in particular two MIA branches are either root- or leaf-specific as shown on the right. In our study, transient overexpression of TFs in *C. roseus* flower petals is used to assess the impact on MIA biosynthesis at the gene expression level (by qPCR) and MIA metabolite level (by LC-MS analysis). Some MIA derivatives are detected in this flower petal expression system in this study, but it is not known where they occur in wild-type plants (upper right). Arrows indicate known characterized enzymatic reactions and corresponding enzymes; dashed black arrows indicate assumed enzymatic or chemical reactions for which no specific enzyme has been identified. Dashed grey arrows indicate transport and corresponding transporters. Note that NPF2.9 is an intracellular transporter conferring vacuolar export of strictosidine and TPT2 is an epidermis-specific transporter exporting catharanthine to the leaf surface. For clarity, only selected MIA metabolites are shown. (b) Co-expression network of MIA biosynthesis genes. From an expression atlas of 82 samples k-nearest neighbor (KNN) networks were created. A line between genes indicates co-expression. Opacity of the lines is scaled according to the mutual rank of co-expression between two genes in their KNN 500 clusters (darker = stronger co-expression). To simplify the complex MIA biosynthesis network, we have colored the different pathway branches. Abbreviations: GES, geraniol synthase; G8O, geraniol-8-oxidase; 8HGO, 8-hydroxygeraniol oxidoreductase; IS, iridoid synthase; IO, iridoid oxidase; 7DLGT, 7-deoxyloganetic acid glucosyl transferase; 7DLH, 7-deoxyloganic acid hydroxylase; LAMT, loganic acid *O*-methyltransferase; SLS, secologanin synthase; TDC, tryptophan decarboxylase; STR, strictosidine synthase; NPF, nitrate/peptide family; SGD, strictosidine β-glucosidase; VAS, vitrosamine synthase; THAS, tetrahydroalstonine synthase; AS, alstonine synthase: HYS, heteroyohimbine synthase; GS, geissoschizine synthase; GO, geissoschizine oxidase; SAT, stemmadenine-*O*-acetyl-transferase; PAS, precondylocarpine acetate synthase; DPAS, dihydroprecondylocarpine synthase; CS, catharanthine synthase; TPT, catharanthine transporter; T16H, tabersonine 16-hydroxylase; 16OMT, 16-hydroxytabersonine *O*-methyltransferase; T3O, tabersonine 3-oxygenase; T3R, tabersonine 3-reductase; NMT, 2,3-dihydro-3-hydroxytabersonine-N-methyltransferase; D4H, desacetoxyvindoline 4-hydroxylase; DAT, deacetylvindoline 4-O-acetyltransferase; PRX, peroxidase; HL, hydrolase; V19H, vincadifformine-19-hydroxylase; MAT, minovincinine-19-*O*-acetyltransferase; T19H, tabersonine 19-hydroxylase; TAT, 19-hydroxytabersonine 19-*O*-acetyltransferase; TEX, tabersonine epoxidase.

There has been a substantial increase in knowledge about which specific TFs from different families control the different MIA pathway steps and branches, although not complete yet. The main focus has been on the jasmonate (JA)-dependent transcriptional activation, because many (but not all, see below) MIA pathway steps are up-regulated upon exposure to this phytohormone that mainly mediates defense against necrotrophic pathogens and herbivores (Aerts *et al.*, 1994; Zhang *et al.*, 2011; De Geyter *et al.*, 2012; Goossens *et al.*, 2016; Zhou and Memelink, 2016; Goossens *et al.*, 2017). The primary response regulator of JA-dependent up-regulation of target genes is the basic helix-loop-helix (bHLH) TF MYC2, whose activity is post-translationally suppressed in the absence of JA (Chini *et al.*, 2007; Goossens *et al.*, 2017; Wasternack and Strnad, 2019). Recently, we have described how overexpression of the de-repressed *CrMYC2a^D126N^* variant can boost the expression of the JA-responsive MIA biosynthesis genes (Schweizer *et al.*, 2018). This up-regulation may partly occur directly and/or via the up-regulation of branch-specific TFs described below (Schweizer *et al.*, 2018). For instance the regulators clade IVa bHLH iridoid synthesis (BIS) 1 and 2 TFs exclusively up-regulate the *iridoid pathway* genes (Van Moerkercke *et al.*, 2015; Van Moerkercke *et al.*, 2016). The expression of *strictosidine pathway* genes is controlled by members of the octadecanoid derivative-responsive *catharanthus* apetala2-domain (ORCA) clade (Menke *et al.*, 1999; van der Fits and Memelink, 2000, 2001; Paul *et al.*, 2016; Paul *et al.*, 2020; Singh *et al.*, 2020). In addition, overexpression of ORCA3 has been shown to increase the expression of the *iridoid pathway* genes to some extent, part of the *stemmadenine pathway* genes and, when overexpressed in combination with *CrMYC2a^D126N^, root specific MIA* genes in a highly synergistic manner (Schweizer *et al.*, 2018). While there is some knowledge about the target gene profiles of the other ORCAs (Paul *et al.*, 2016; Paul *et al.*, 2020; Singh *et al.*, 2020), a direct comparison of them under the same experimental condition is currently lacking. Moreover, it has not been clarified, which, if any, of the ORCAs, in addition to ORCA3, act synergistically with CrMYC2a^D126N^. Noteworthy, the *vindoline pathway* does not appear to be up-regulated by JA and seems to be rather repressed upon overexpression of *CrMYC2a^D126N^* (Schweizer *et al.*, 2018). Very recently it has been shown that, the *vindoline pathway* is at least in part up-regulated by a GATA TF and down-regulated by phytochrome-interacting factor TFs in a light-dependent manner (Liu *et al.*, 2019). However, the moderate changes in expression of the vindoline pathway genes and metabolite levels induced by these TFs suggest the presence of additional regulators. Other TFs, with a more limited impact on MIA biosynthesis have been described [for a review, see Liu *et al.* (2017)].

Together, the increasing identification of MIA biosynthesis genes over the recent years revealed a network of pathway branches (*subdivisions*) with differential gene expression profiles. This suggests the existence of specific regulatory factors acting independently or cooperatively in a branch-specific manner. Here, we sought out for regulators for pathway branches for which regulation was unclear or unknown by using co-expression analysis of publicly available and hitherto unpublished in-house generated RNA-Seq datasets. We identified additional members of well-known BIS, ORCA and MYB clades. Our results further suggest that subfunctionalization of the ORCA TFs leads to differential regulation of particular subsets of MIA pathway genes. Likewise, synergistic up-regulation of MIA pathway genes in combination with de-repressed CrMYC2a appears to be paralog dependent.

## RESULTS

### TF candidate selection and screening impact on MIA biosynthesis

To enable comprehensive mining of *C. roseus* transcriptomes for novel MIA regulators, first a total of 82 RNA-Seq datasets were mapped to the latest *C. roseus* draft genome version (Kellner *et al.*, 2015) (http://medicinalplantgenomics.msu.edu/index.shtml). In addition to publicly available datasets, RNA-Seq data from hitherto unpublished in-house experiments were included, including data of cell type-enriched transcriptomes (see methods and Table S1). Across these datasets, branches of MIA pathway genes (Fig. 1b) showed a high level of co-expression. Noteworthy, in particular the *vindoline* and catharanthine branches were not co-expressed with other MIA pathway branches (Fig. 1b). A k-nearest neighbor (KNN) network was created and for each MIA biosynthesis gene the top 500 most co-expressing genes were searched for genes with the GO term ‘regulation of gene expression’ (GO:0010468) (see method section for details) yielding a list of 111 candidates (Fig. 2a). Manual sequence analysis of the candidates revealed that 15 were likely not unambiguously corresponding to genuine TFs and thus not considered for further analysis. Consistent with previous results, among the co-expressed candidates we found the previously characterized TFs BIS1 and 2 and ORCA2, 4, 5 and 6, but not ORCA3. We further selected those TFs that showed co-expression with (i) multiple pathway genes and (ii) MIA pathway genes for which till now no transcriptional regulators are known, in particular genes belonging to the *stemmadenine* or *vindoline* branches or involved in catharanthine biosynthesis, and that were not predicted to act as repressors. Out of the 40 selected candidates, 28 were successfully cloned comprising all but two candidates that were ranked as candidates of highest priority and subjected to agroinfiltration in *C. roseus* flower petals. Four additional candidates, that were not found through this co-expression analysis but through a preceding, preliminary, analysis, were also tested in this screening (gene IDs in italic font in Fig. 2b). The latter preceding co-expression analysis was performed exactly as described above but on 72 RNA-Seq samples because one dataset (Pan *et al.*, 2018) had not yet been available at the start of this project (data not shown). While the candidate lists of the two analyses greatly overlapped those four candidates were exclusively found in the first analysis. Thus altogether 32 candidates were tested by bulk overexpression in *C. roseus* flower petals as described hereafter.

**Fig. 2.**
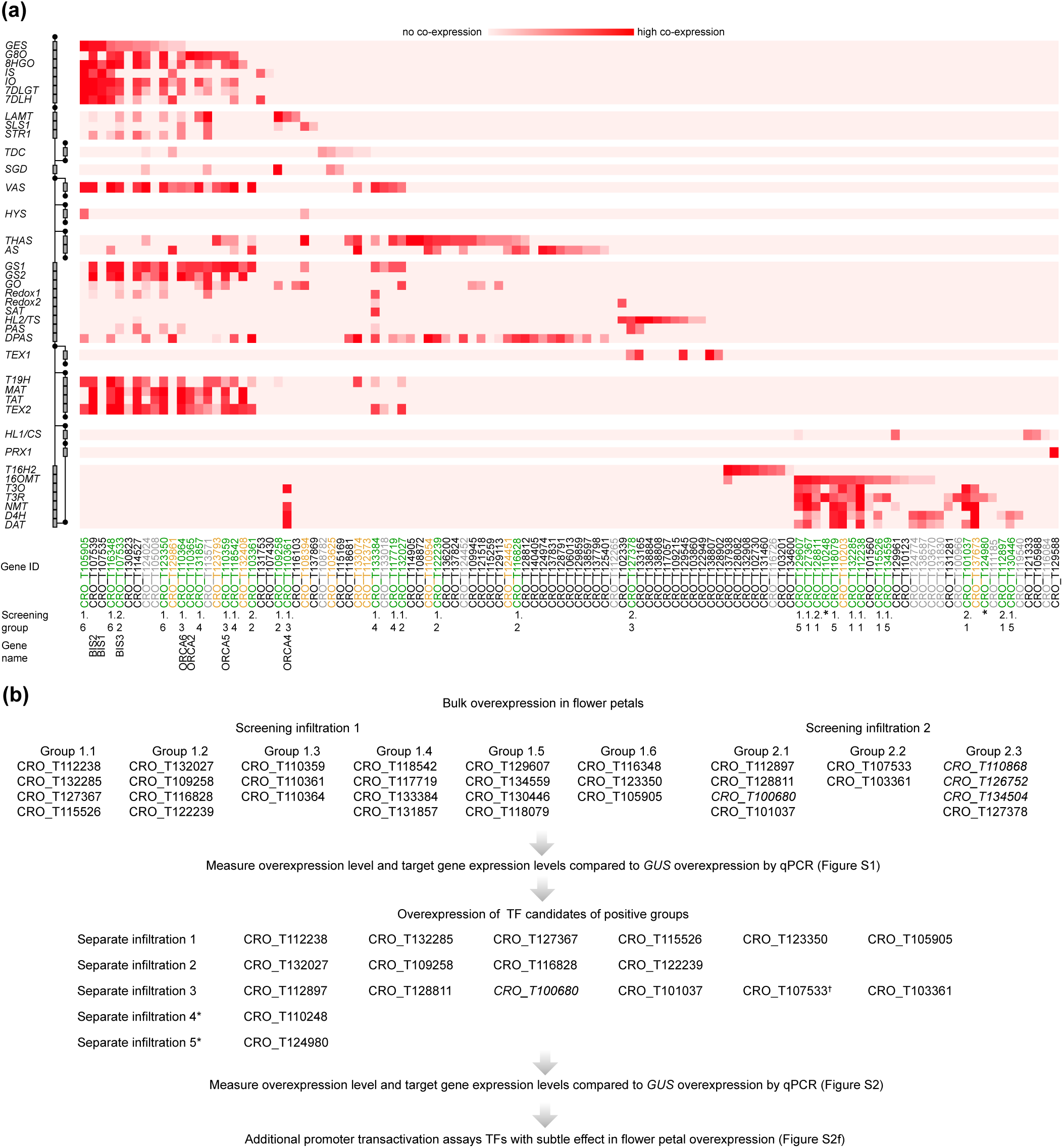
MIA regulator candidate selection and screening procedure. (a) Pathway genes (left) are plotted against regulator candidates (below); the relative rank of a regulator in the top 500 most co-expressed genes is depicted in red, with dark red indicating low ranks and strong associations between regulator and pathway gene. TF gene IDs are color-coded, indicating which genes by manual sequence analyses are likely not specific TFs and were thus not further considered (gray), which were considered for screening but cloning failed (yellow), and which were tested (green). Previously characterized TFs are labeled with their names in addition to “BIS3”, identified in this study. (b) Overview about the screening procedure. To enhance the output, TF candidates were bulk-overexpressed (“Screening groups”) by mixing *A. tumefaciens* strains containing the respective constructs. The screening was performed in two independent rounds of infiltration (“Screening infiltration 1 and 2”). In case of a detected up-regulation of target genes >2-fold and P<0.05 (tested by Student’s *t*-test) (Fig. S1), the TF candidates from the respective group were overexpressed separately (Fig. S2). Gene IDs in italics were not picked in this co-expression analysis but in a previous one.

To enhance the throughput, up to four TF candidates were co-overexpressed under the control of the *CaMV35S* promoter (p35S). We specifically combined those candidates that were co-expressed with the same (or overlapping) subsets of MIA pathway genes. This resulted in nine overexpression groups of candidates (Fig. 2b). *A. tumefaciens* harboring a plasmid for *p35S::GUS* expression was infiltrated as a control. The overexpression level of the candidate TFs and the expression level of the MIA pathway genes were then measured by qPCR (Fig. S1). In case of successful overexpression (>5-fold) and observed increase(s) (>2-fold, P-value according to Student’s *t*-test <0.05) in expression level(s) of potential target MIA pathway genes compared to the *p35S::GUS* control, the TF candidates from the respective group were overexpressed separately (Fig. 2b; Fig. S2). In the case of CRO_T132285 and CRO_T127367, for which the increase in target gene expression level was at the threshold level, additional promoter transactivation assays were carried out that did not confirm a (direct) up-regulation of MIA target genes (Fig. S2f). Altogether the results from the screening suggested that only the ORCA TFs (CRO_T110359, CRO_T110361, CRO_T110364) and clade IVa bHLH TFs (CRO_T123350, CRO_T105905, CRO_T107533) reproducibly led to an increase in MIA biosynthesis gene expression levels (Fig. S1, S2 and below).

### The *iridoid* regulatory module: Identification of an additional *BIS* paralog that clusters with *BIS1* and *2*

Overexpression of the three newly identified iridoid pathway co-expressed (Fig. 2) clade IVa bHLHs (CRO_T123350, CRO_T105905, CRO_T107533) led to enhanced expression of the iridoid genes, albeit to very different levels (Fig. 3a). Only overexpression of CRO_T107533 (referred to as *BIS3* hereafter) led to increases in *iridoid pathway* transcript levels (28-to 480-fold) similar to overexpression of *BIS1* and *2* in our previous study (Van Moerkercke *et al.*, 2016). This high up-regulation of iridoid pathway genes by BIS1, 2 and 3 in comparison to the CRO_T123350 and CRO_T105905 was further confirmed by promoter transactivation assays in transient expression assays in *Nicotiana tabacum* (tobacco) protoplasts on (Fig. 3b). We did not observe consistent differential or synergistic effects of combinations of the three BIS on iridoid pathway gene promoters (Fig. S3), suggesting that homo-or heterodimers of BIS1/2/3 have similar transactivation capacities and specificities. A ClustalW alignment of the highly conserved bHLH domains of the five clade IVa factors reveals a higher level of sequence identity between the three BIS (BIS2 and 3 bHLH sequences are identical) compared to CRO_T123350 and CRO_T105905 (Fig. 3c). Noteworthy, all three *BIS* TFs cluster within a 120-kb locus in the genome and are probably the result of recent tandem gene duplication events (Fig. 3D). This was previously not known but could now be detected because of the recent update (v2) of the *C. roseus* genome (http://medicinalplantgenomics.msu.edu).

**Fig. 3.**
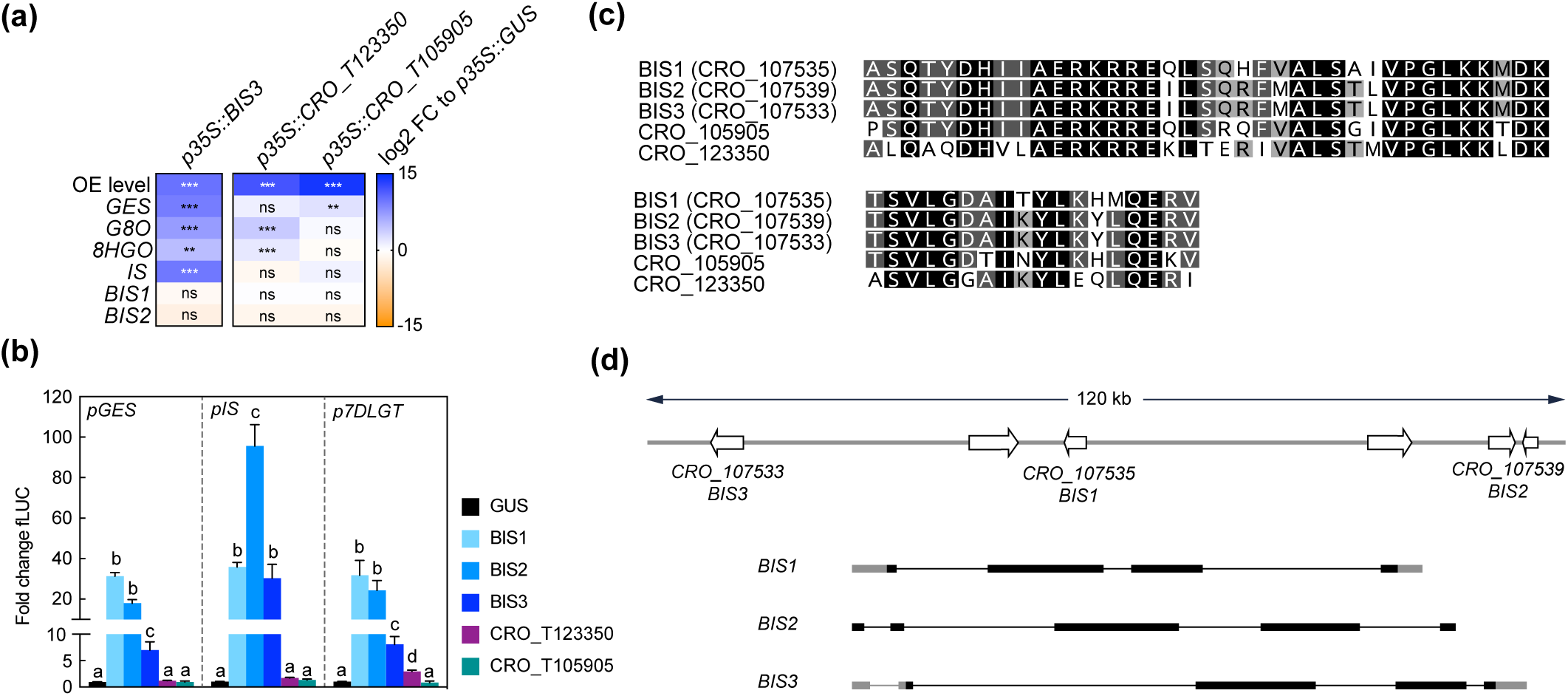
BIS3 up-regulates iridoid pathway genes. (a) Overexpression of *BIS3* leads to increased expression levels of iridoid pathway genes similar to those previously observed for *BIS1* and *2*, whereas overexpression of the other clade IVa TFs CRO_T123350 and CRO_T105905 leads to comparably much lower increases. *BIS1* and *2* themselves are not up-regulated by BIS3, suggesting that the iridoid pathway up-regulation is directly caused by BIS3 and that there is no cross up-regulation between the three BIS TFs. Asterisks indicate statistically significant differences in expression (*P < 0.05, **P < 0.01, ***P < 0.01) as calculated by Student’s *t*-test for each gene compared to GUS control (not included in this graph). (b) *N. tabacum* protoplasts were co-transfected with constructs containing the *firefly luciferase* gene (*fLUC*) expressed under control of the indicated promoter fragments and constructs for overexpression of *bHLHs* as indicated or *GUS* as a control. The y-axis shows fold change in normalized fLUC activity relative to the control transfection with GUS, set at 1. The error bars depict SEM of four biological replicates. For each promoter, columns labelled with different letters represent statistically significant differences (P<0.05, ANOVA with Tukey’s correction for multiple comparisons). BIS1-3 redundantly highly transactivate iridoid pathway promoters whereas CRO_T123350 and CRO_T105905 have no effect. (c) A MUSCLE alignment of the bHLH domain protein sequences reveals a high level of sequence identity between the five clade IVa bHLHs. However, BIS1, 2 and 3 show a slightly higher level of identity than CRO_T123350 and CRO_T105905 potentially explaining the differential level of up-regulation of iridoid pathway genes. Sites that are identical in all five sequences are colored in black; sites identical in four out of five sequences are colored in dark gray and sites identical in 3 out of 5 sequences are colored in light gray. (d) The latest version of the *C. roseus* genome revealed that the previously published BIS1 and 2, and BIS3 identified in this study, are all located in close proximity and have similarly organized gene structures suggesting that they probably arose from recent gene duplications. Exons are depicted as bars with untranslated regions (UTRs) in gray and CDS’ in black. Introns are depicted as black lines.

### The ORCA cluster: Functional diversification into branch-specific regulatory modules

In addition to the long-known *ORCA2* and *3*, *ORCA4, 5* and *6* have recently been found to be members of a gene cluster together with *ORCA3*, and their target gene profile has partly been elucidated (Paul *et al.*, 2016; Paul *et al.*, 2020). All ORCA TFs together form a cluster spreading over 70 kb, which can be subdivided into an ORCA3, 4 and 5 subcluster (around 25 kb) and an ORCA2 and 6 subcluster (around 10 kb) (Fig. 4a). A combination of different computational sequence analyses led us to hypothesize that the five ORCA TFs are possibly not fully functionally redundant. First, key amino acid residues (AARs) within the DNA-binding domain (DBD) differ between the ORCAs and are predicted to confer direct interaction with DNA differ (Fig. 5a). Second, it is generally accepted that the activation domains (ADs) of TFs are mostly poorly conserved at sequence level, but it is rather the structural properties defining the level of activation (Staby *et al.*, 2017). The combined information of different prediction programs suggested that the ORCA TFs differ significantly in their structural organization, further hinting towards a certain degree of diversification (Fig. S4). Third, ORCA3-5 contain a serine-rich C-terminal domain that ORCA2 and 6 lack. Fourth, the five ORCA TFs showed a differential expression profile (Fig. 4b and S5) and our co-expression analysis revealed that they co-express with different subsets of MIA pathway genes (Fig. 2a). Despite the occurrence of clusters of ORCA orthologs in other species, such as tomato and tobacco, it has not been systematically analyzed in any of these species if and to which level they are functionally redundant (Cárdenas *et al.*, 2016; Kajikawa *et al.*, 2017). Therefore, we sought out to decipher their specific regulatory roles by comparing their target gene and metabolite profile upon overexpression in flower petals. A PCA analysis of the MIA gene expression levels revealed that ORCA2-4 showed only partly overlapping target gene profiles (Fig. 4c). Particularly the expression of early and intermediate MIA pathway genes increased to similar levels upon overexpression of ORCA2-4 (Fig. 4d, for instance *SLS1*, *STR1*, *GS1*, *GS2*, *REDOX1*,*2*). However, in other cases, the fold changes differed considerably. For instance, *GO*, which encodes an enzyme producing the intermediate geissoschizine but also akuammicine (see Fig. 1a), as well as the iridoid pathway genes including the regulator BIS1, were considerably more highly up-regulated upon overexpression of *ORCA4*, and to a lesser extent of *ORCA2* compared to overexpression of the other *ORCAs* (Fig. 4d). Conversely, the *root MIA* genes *T19H* and *MAT* were more highly up-regulated when *ORCA3* or *ORCA5* were overexpressed. It should be noted that the overexpression level of the different *ORCA* paralogs differed considerably, with *ORCA3, 4* and *6* being highly overexpressed (around 510-, 440-, and 960-fold, respectively) and with *ORCA2* and in particular *ORCA5* somewhat less (93-, and 15-fold, respectively) overexpressed (Fig. 4d). Thus, the overall lower MIA biosynthesis gene up-regulation levels observed for *ORCA5* overexpression might be partly due to the lower overexpression level of *ORCA5*. We did not observe any “intra-cluster” up-regulation between the different ORCA TFs. We next performed metabolite profiling on samples from this experiment (Figs 4g-f, S6, Table S6). A PCA analysis of the Mass spectrometry (MS) data revealed a (partially) differential profile (Fig. 4e), which was caused by differential accumulation of MIA metabolites (Fig. 4f). In addition to MIAs identified in a previous study (Schweizer *et al.*, 2018), the loadings plot further showed differentially accumulating compounds that we identified as MIAs (Fig. 4f (red), Supporting Dataset S2). In particular, akuammicine, 16-hydroxytabersonine and 16-hydroxylochnericine accumulation increased more when *ORCA4* was overexpressed and that of the root MIA derivative 16-methoxyhörhammericine when *ORCA3* was overexpressed (Fig. 4g). Overexpression of *ORCA6* had only mild effects on MIA pathway gene expression and no noticeable effects on MIA abundance. Promoter transactivation assays in tobacco protoplasts with promoter fragments of *STR*, *GO*, *MAT*, and *TAT* expressed with and without the addition of CrMYC2a^D126N^ overall confirmed these observations (Fig. 7). ORCA4 transactivated *pGO* to a much higher extent than the other ORCA TFs, whereas ORCA3 transactivated *pTAT* higher than other ORCAs, wheras *pSTR* was similarly transactivated by ORCA3, 4 and 5. Together, these data support both redundant and specific functions of the different ORCA TFs in the regulation of MIA biosynthesis.

**Fig. 4.**
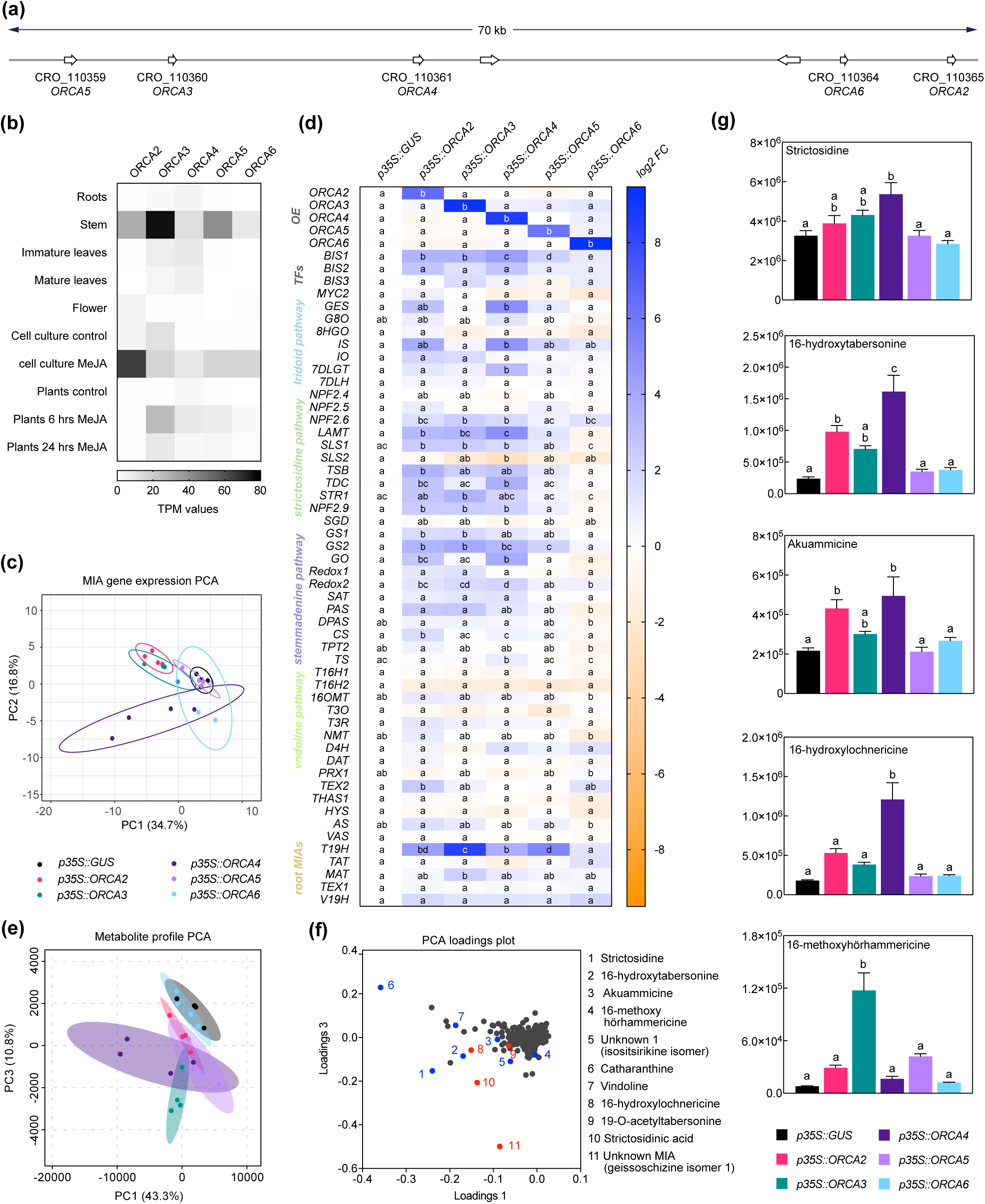
Members of the ORCA TF cluster differentially up-regulate MIA biosynthesis genes. (a) Localization of the *ORCA* genes within the cluster. (b) Expression pattern of *ORCA* paralogs extracted from publicly available RNA-Seq data from indicated tissues and conditions. TPM, Transcripts Per Kilobase Million. (c) A PCA of MIA pathway genes measured by qPCR upon *ORCA* overexpression (shown in d) suggests overlapping but also differential MIA target gene expression patterns in particular between *ORCA2, 3* and *4*. PCA was performed with ClustVis (Metsalu and Vilo, 2015). (d) Expression of MIA biosynthesis genes measured by qPCR upon *ORCA* overexpression shown as log2 fold changes (FC) compared to the overexpression of *GUS* as a control. For each pathway gene, cells labeled with different letters represent statistically significant differences (P<0.05, ANOVA with Tukey’s correction for multiple comparisons). Note that the overexpression levels of *ORCA3, 4* and *6* were higher than those of *ORCA2* and *5*. (e) PCA of metabolite profiling data from *ORCA* overexpression samples. (f) Loading plot of the PCA shown in (e), indicating that the differences found in the PCA are indeed predominantly caused by MIA metabolite levels. Selected known MIAs are labeled in blue; MIAs identified in this study (see Supporting Dataset S1) are labeled in red. (g) Levels of selected MIA metabolites expressed in average total ion current (TIC). The levels of all measured MIA metabolites can be found in the supporting information (Fig. S5; Table S6). Columns labeled with different letters represent statistically significant differences (P<0.05, ANOVA with Tukey’s correction for multiple comparisons). Error bars are standard error of the mean (SEM) of four biological replicates.

**Fig. 5.**
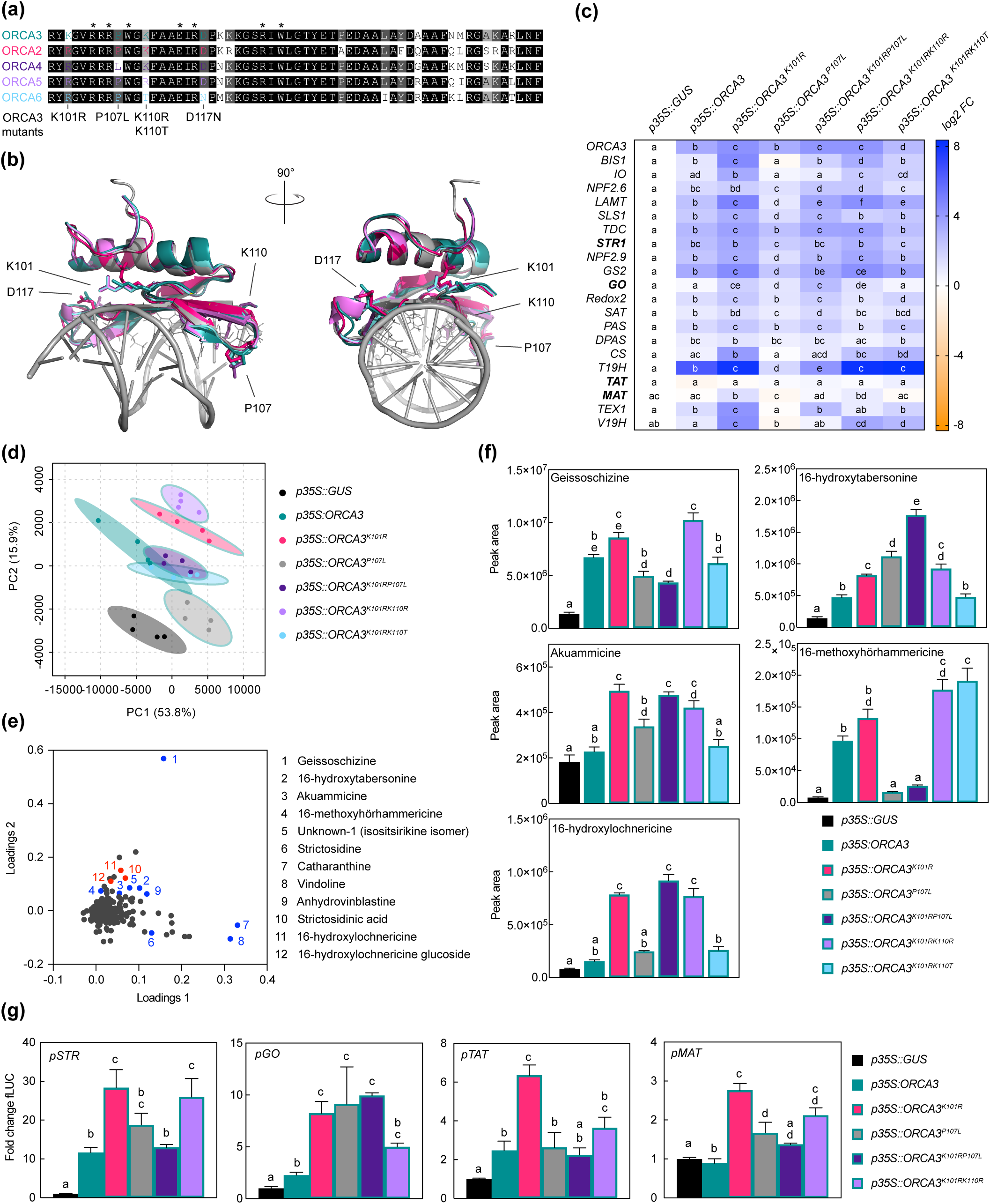
Specific amino acid residues (AARs) in the DNA-binding domain (DBD) of ORCA TFs influence target gene specificity. (a) Alignment of the DBDs of ORCAs. AARs labeled with an asterisk were identified previously as directly interacting with the DNA based on the solved crystal structure of the *Arabidopsis* AP2/ERF member AtERF1 DBD (Allen *et al.*, 1998; Shoji *et al.*, 2013). ORCA3 was used as a template to exchange AARs so that they would correspond to AARs in other ORCA TFs. The respective mutated AARs are indicated below the alignment. (b) Homology models of ORCA DBDs. The models (made by Phyre2 “intensive mode” (Kelley *et al.*, 2015); colored as in panel a) are overlaid with the AtERF1 structure bound to DNA (PDB 1GCC) colored in gray. AARs identified as directly interacting with the DNA (asterisks in panel a) are shown as lines of AtERF1. AARs differing in the ORCA DBDs and selected for site-directed mutagenesis are shown as sticks for all ORCA models, while the AAR numbering refers to ORCA3. The models suggest that the differing AARs are unlikely to interact directly with the DNA, but are in close proximity to the DNA and might thereby modulate DNA binding affinity. (c) Overexpression of *ORCA3* mutants in *C. roseus* flower petals and expression level of selected MIA target genes as measured by qPCR. Two AAR changes, K101R and P107L, seem to significantly modulate the MIA target gene profile. While K101R appears to increase the level of up-regulation, up-regulation is greatly reduced and in some cases nearly abolished when overexpressing the *ORCA3^P107L^* mutant. The double mutant *ORCA3^K101RP107L^*, corresponding to the native ORCA4 sequence, recovers some activation potential, and reaches similar levels than observed upon overexpression of *ORCA4* (Fig.4d). This suggests that the combination of these AARs may account for the differences between ORCA3 and 4. For each pathway gene, cells labeled with different letters represent statistically significant differences (P<0.05, ANOVA with Tukey’s correction for multiple comparisons). Results from overexpression of all *ORCA3* single AAR mutants (including *ORCA3^D117N^*) can be found in Fig. S6. (d) PCA of metabolite profiling data. (e) Loading plot indicating that sample separation shown in the PCA plot is largely due to changes in MIA metabolite levels. Selected known MIAs are labeled in blue; MIAs identified in this study are labeled in red. (f) Levels of selected MIA metabolites expressed in average total ion current (TIC). The levels of all measured MIA metabolites can be found in the supporting information (Fig. S7; Table S7). Columns labeled with different letters represent statistically significant differences (P<0.05, ANOVA with Tukey’s correction for multiple comparisons). Error bars are standard error of the mean (SEM) of four biological replicates. (g) Transactivation assays of *pSTR*, *pGO*, *pTAT* and *pMAT* performed in tobacco protoplasts. Columns labeled with different letters represent statistically significant differences (P<0.05, ANOVA with Tukey’s correction for multiple comparisons). Error bars are standard error of the mean (SEM) of four biological replicates.

**Fig. 6.**
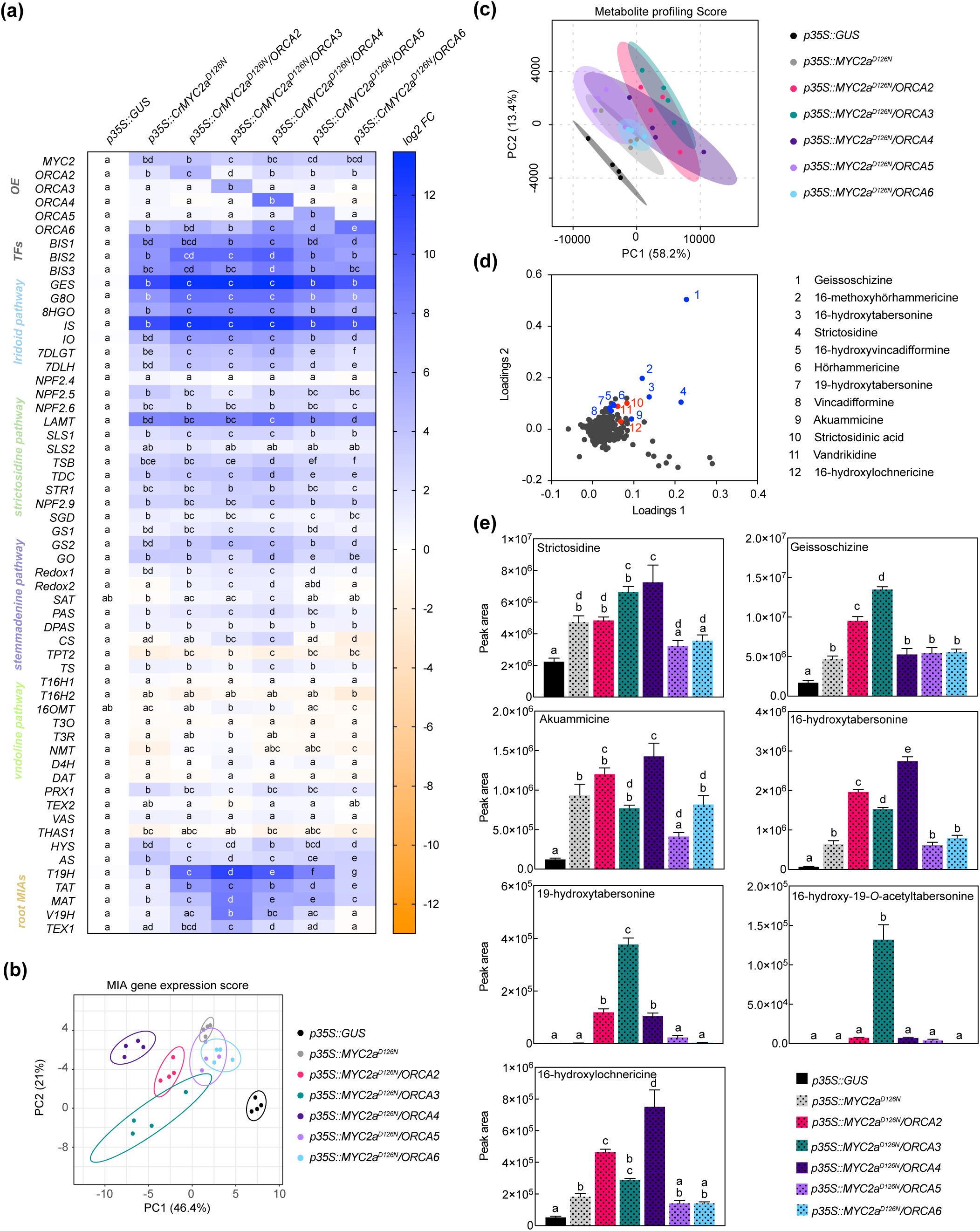
Combinatorial overexpression of *CrMYC2a^D216N^* and *ORCAs*. (a) Expression profile of MIA pathway genes and overexpression levels. For each pathway gene, cells labeled with different letters represent statistically significant differences (P<0.05, ANOVA with Tukey’s correction for multiple comparisons). Note that synergistic up-regulation of particularly root MIA genes is most pronounced by combinatorial overexpression of *CrMYC2a^D126N^* and *ORCA3*. (b) PCA of the MIA gene expression data is suggesting that in particular the combinatorial overexpression of *CrMYC2a^D126N^* and *ORCA2*, *3* and *4* lead to differential changes in the MIA gene expression pattern, whereas combinatorial overexpression with *ORCA5* or *6* does not differ substantially from overexpression of *CrMYC2a^D126N^* alone. PCA was performed with ClustVis (Metsalu and Vilo, 2015). (c) PCA of metabolite profiling data. (d) Loading plot of PCA with selected known MIAs (blue) and MIAs identified in this study (red). (e) Levels of selected MIA metabolites expressed in average total ion current (TIC). The levels of all measured MIA metabolites can be found in the supporting information (Fig. S7; Table S8). Columns labeled with different letters represent statistically significant differences (P<0.05, ANOVA with Tukey’s correction for multiple comparisons). Error bars are standard error of the mean (SEM).

**Fig. 7.**
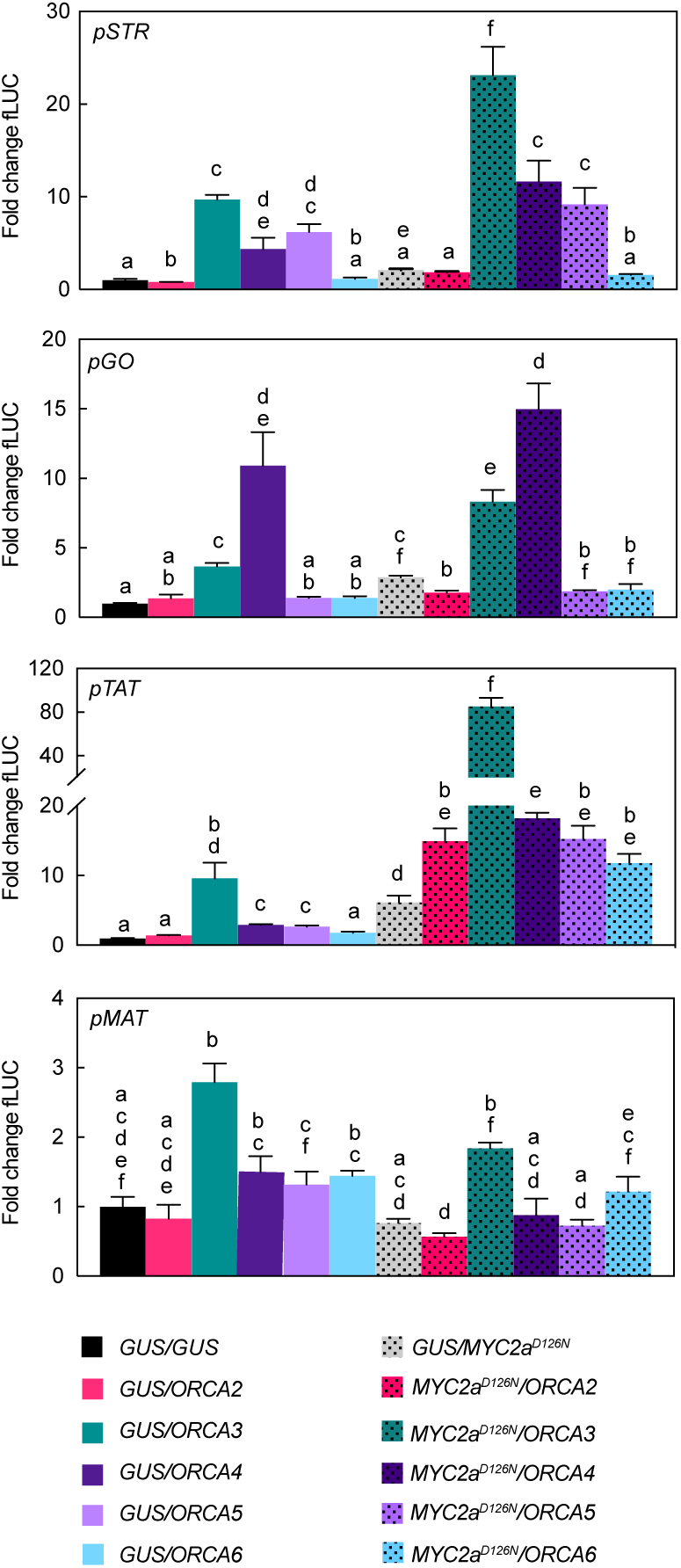
Transactivation of *STR*, *GO*, *TAT* and *MAT* promoter fragments by ORCAs and CrMYC2a^D126N^. *N. tabacum* protoplasts were co-transfected with constructs containing the *fLUC* gene expressed under control of the indicated promoter fragments and constructs for overexpression of *ORCAs, CrMYC2a^D126N^* or *GUS* as a control. Note in particular differential transactivation of *pGO* and *pTAT* upon *ORCA3* versus *ORCA4* expression and synergistic effects on *pTAT* upon combined overexpression of *ORCA3* and *CrMYC2a^D126N^* in combination. The y-axis shows fold change in normalized fLUC activity relative to the control transfection with GUS, set at 1. The error bars depict SEM of four biological replicates. For each promoter, columns labelled with different letters represent statistically significant differences (P<0.05, ANOVA with Tukey’s correction for multiple comparisons).

### Specific AARs within ORCA DBDs contribute to target gene specificity

Generally, the DBD is thought to define the specificity to the DNA motif, whereas the AD defines the level of up-regulation by recruiting the transcriptional machinery. In a previous study, the affinities of different orthologs of group IXa AP2/ERFs, i.e. ORCA3 and its orthologs from tobacco and *Arabidopsis thaliana*, to the DNA motif were compared (Shoji *et al.*, 2013). The results suggested that the specific binding affinity to different DNA motifs is indeed mediated by specific AARs in the DBD. However, specific AARs mediating target gene specificity among paralogs from the same species have never been characterized in detail to our knowledge. A protein sequence alignment of the DBDs of the ORCAs shows the expected overall high level of sequence conservation (Fig. 5a). Likewise, structural models of the ORCA DBDs are almost identical and highly align with the solved structure (PDB 1gcc) of the Arabidopsis ortholog AtERF1 (Allen *et al.*, 1998) (Fig. 5b). While the paralog-specific AARs corresponding to K101, P107, K110 and D117 in ORCA3 are not predicted to directly interact with DNA in the shown model, they are nevertheless predicted to be part of the beta-sheets in close proximity to the DNA (Fig. 4b). To compare a potential impact of these AARs on MIA target gene specificity, we performed site-directed mutagenesis on ORCA3 to obtain single (K101R, P107L, K110R, K110T and D117N) and double (K101R/P107L, K101R/K110R_K101R/K110T) mutants, whose sequence would correspond to that of the other paralogs. The mutated versions were then overexpressed alongside native *ORCA3* in flower petals in two independent infiltration series (Fig. Fig. 5c-f and Fig. S7, respectively). Overall, our results pointed to two key amino acid residues leading to changes in MIA target gene expression. Compared to overexpression of native *ORCA3*, overexpression of *ORCA3^K101R^* (corresponding to an AAR change occurring in ORCA2, 4, 5 and 6) generally enhanced the expression of MIA pathway genes (Figs 5c, S7). By contrast, overexpression of *ORCA3^P107L^* (corresponding to one of the AAR changes in ORCA4) led to reduced or abolished levels of MIA pathway genes that were partly recovered in the double mutant *ORCA3^K101RP107L^* (corresponding to AARs in ORCA4) (Fig. 5c). Noteworthy, overexpression of *ORCA3^K101RP107L^* still led to a lower up-regulation of *root MIA* genes than *ORCA3* or *ORCA3^K101R^*, suggesting that the combination of these two AARs contributes to the differential target gene profile of ORCA3 and 4. The impact of the other AARs appeared to be less specific. Overexpression of the *ORCA5*-like *ORCA3^K101RK110R^* led to an expression profile similar to that upon overexpression of *ORCA3^K101R^*. Also, the much lower level of MIA target gene up-regulation by ORCA6 compared to the other ORCAs was not explained by the selected AARs (compare the overexpression of *ORCA3^K101RK110T^* (Fig. 5c) and *ORCA3^D117N^* (Fig. S7)). However, we did not produce a triple mutant in which all three AARs would correspond to those of ORCA6. The observed differential target gene profiles upon overexpression of *ORCA3* mutants were also reflected at the metabolite level (Figs 5d-f, S8; Table S7). Noteworthy, in particular the levels of 16-hydroxytabersonine, akuammicine, 16-methoxyhörhammericine and 16-hydroxylochnericine (among others) upon overexpression of *ORCA3* versus the ORCA2-like *ORCA3^K101R^* and the ORCA4-like *ORCA3^K101RP107L^* were very similar to those upon overexpression of the actual *ORCA2, 3* and *4* (Fig. 4g and S6). We further performed transactivation assays with ORCA3, ORCA3^K101R^, ORCA3^P107L^, ORCA3^K101RP107L^ and ORCA3^K101K110R^ on *pSTR*, *pGO*, *pTAT* and *pMAT* (Fig. 5g). Similarly to what was observed in the flower petal overexpression experiments ORCA3^K101R^ appeared to have a higher overall transactivation effect than ORCA3 on the four promoters whereas the double mutant ORCA3^K101RP107L^ transactivated pGO at significantly higher levels than ORCA3 but not the other promoters.

### Synergistic up-regulation in combination with de-repressed CrMYC2a^D126N^ is both ORCA- and target gene-dependent

Previously, we have shown that combinatorial overexpression of *ORCA3* and the de-repressed *CrMYC2a^D126N^* variant leads to a synergistic up-regulation in particular of the *root MIA genes* compared to independent overexpression (Schweizer *et al.*, 2018). Based on this and our experiments above that suggest that the ORCA TFs to some extent up-regulate different subsets of MIA pathway genes, we next investigated whether the combined overexpression of *CrMYC2a^D126N^* with *ORCA2, 4, 5* or *6* leads to a synergistic up-regulation of target genes similar as with *ORCA3* and if different subsets of MIA pathway genes would be up-regulated (Fig. 6a). Indeed, a PCA depicted that in particular the combined overexpression of *CrMYC2a^D126N^* with *ORCA2*, *3* or *4* led to differences in the overlapping MIA pathway gene expression profiles (Fig. 6b). Accordingly, PCA analysis of the MS data (Fig. 6c) revealed somewhat differential metabolite profiles for the combination of *CrMYC2a^D126N^* with these three *ORCA* genes rather than with *ORCA5* or *6,* or upon overexpression of *CrMYC2a^D126N^* alone; these changes were due to differential accumulation of MIAs (Fig. 6d). With regard to synergistic effects, the combined overexpression of *CrMYC2a^D126N^* with *ORCA2* or *4*, synergistically up-regulated the *root MIA* genes, but to a lower extent than with *ORCA3* (Fig. 6a). Accordingly, at the metabolite level, the *root MIAs* 19-hydroxytabersonine and 16-hydroxy-19-O-acetyltabersonine accumulated to much higher levels upon *CrMYC2a^D126N^/ORCA3* overexpression than upon overexpression of any of the other combinations (Fig. 6e). Notably, no synergistic, if anything an additive up-regulation was observed for *GO*, although it was up-regulated upon overexpression of *CrMYC2a^D126N^* and *ORCA4* alone (Figs 4d, 6a). Likewise, the level of the respective metabolic product akuammicine was only less than 2-fold higher in the samples with combined overexpression of *CrMYC2a^D126N^* with *ORCA4*, compared to when *CrMYC2a^D126N^* was overexpressed alone (Fig. 6e). These two observations suggest that the ORCA-CrMYC2a^D126N^ synergistic module is partly TF-dependent, i.e. more pronounced for ORCA3 and, moreover, target gene-dependent, i.e. specific to some of the *root MIA* genes. To further confirm these observations promoter transactivation assays on *pSTR, pGO, pTAT* and *pMAT* were carried out (Fig. 7). Synergistic transactivation occurred particularly in the case of the *root MIA* gene promoter *pTAT* in combination with ORCA3 but could not be confirmed for the other root MIA promoter *pMAT*. However, transactivation levels for *pMAT* were generally low and it is possible that the 2-kB fragment did not contain all necessary cis regulatory sites. We were unable to clone the promoter region of *T19H* in several attempts. Transactivation of *pGO* by CrMYC2a^D126N^ and ORCA4 was not synergistic, again as observed in overexpression in flower petals.

To gain more insight into how a synergistic up-regulation could be mediated at the molecular level and to possibly shift the subset of target genes of this synergy, we next made chimeric constructs. Here, we focused on *ORCA3* and *4*, because their level of overexpression was most similar and the level of up-regulation of target genes was the strongest for these two paralogs. In these chimeric constructs, the N-terminal AD of ORCA3 was fused to the C-terminal DBD of ORCA4 (ORCA3-ORCA4) and *vice versa* (ORCA4-ORCA3) (Fig. 8a). Overexpression of these “*ORCA*-swaps” in combination with *CrMYC2a^D126N^* led to shifts in the up-regulation of root MIA genes (Fig. 8b). Overexpression of *ORCA3-ORCA4* led to a 2 to 4 fold higher up-regulation of *root MIA genes* than overexpression of native *ORCA4*, whereas overexpression of *ORCA4-ORCA3* led to around 4-fold lower up-regulation than overexpression of native *ORCA3.* These results indicate that the AD of ORCA3 might at least in part be necessary for the synergistic up-regulation. On the other hand, overexpression of *CrMYC2a^D126N^*/*ORCA3-ORCA4* did not lead to a higher up-regulation of *GO* than overexpression of *CrMYC2a^D126N^*/*ORCA4*, further supporting the previous observation that the synergistic up-regulation is specific to *root MIA* genes, i.e. target gene specific.

**Fig. 8.**
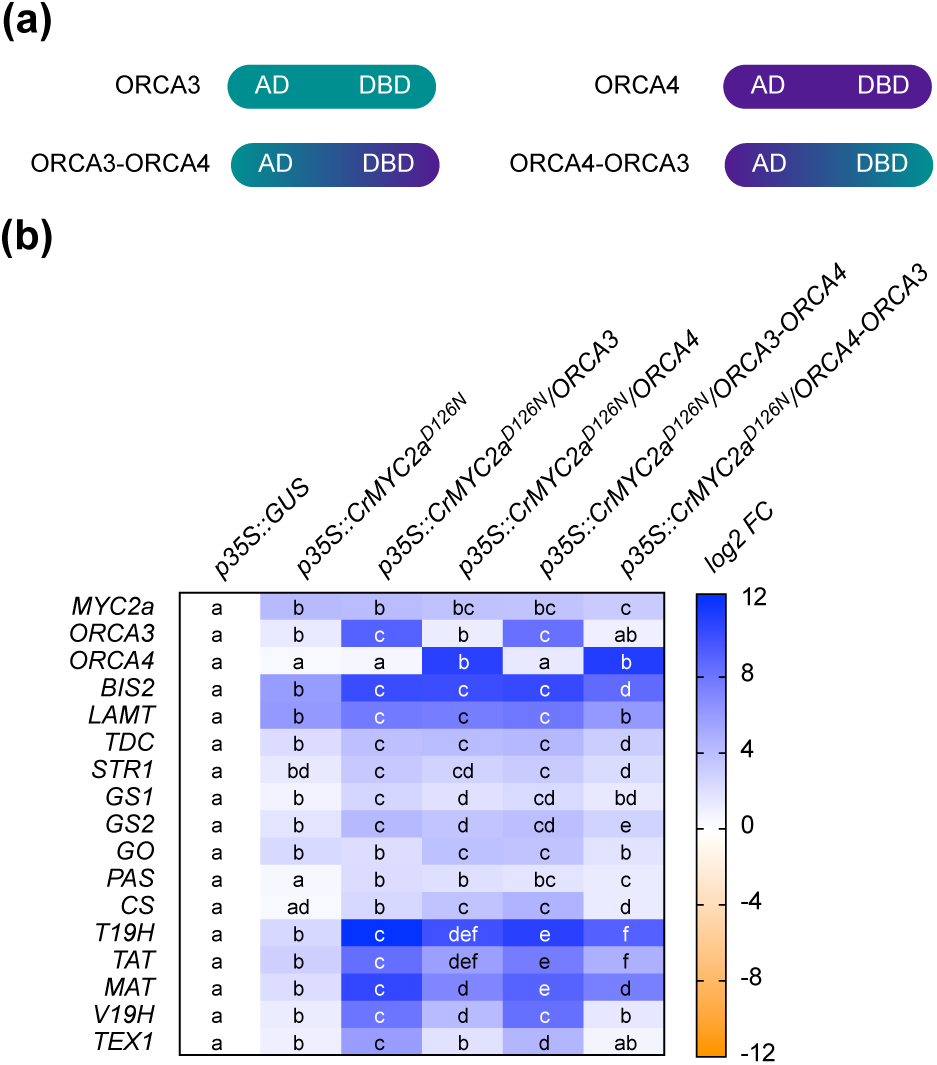
Swapping of DNA-binding domain (DBD) and activation domain (AD) of ORCA3 and 4. In order to get insight into the role of the AD in the synergistic up-regulation of MIA target genes, the respective DBDs and ADs of ORCA3 and 4 were swapped. (a) Cartoon showing the domain organization of ORCA3 and 4 and the respective chimeric proteins. (b) Combinatorial overexpression of *CrMYC2a^D126N^* with *ORCA3, 4* or chimeric proteins *ORCA3-ORCA4* or *ORCA4-ORCA3*. These results suggest that the AD of ORCA3 improves the synergistic up-regulation of target genes. Cells labeled with different letters represent statistically significant differences (P<0.05, ANOVA with Tukey’s correction for multiple comparisons).

### Putative cell-specific regulation of the *vindoline* branch

Data from the cell type-enriched transcriptomes pointed towards a number of epidermis-specific TFs that might be candidate regulators of the epidermis-specific expression of some MIA pathway genes (*strictosidine* and *stemmadenine* branches and parts of the *vindoline* branch and catharanthine biosynthesis). Among these potential candidates, two MYB TFs were highly expressed in the stem epidermis (Fig. S10b). Both MYB TFs, hereafter named MYB96 and MYB96b, appeared to be potential orthologs of AtMYB96 and 94 (Fig. S10a), which amongst others positively regulate cuticular wax biosynthesis genes in Arabidopsis (Seo *et al.*, 2011; Lee and Suh, 2015; Lee *et al.*, 2016). In a preliminary screen, both MYB TFs were found to transactivate the *T3R* promoter in tobacco protoplasts, albeit not additively (Fig. S10c). Overexpression of *MYB96* in flower petals also led to an up-regulation of expression of *T3R* and of a putative ortholog of the *Arabidopsis* cuticular wax biosynthesis gene *CER1*, but this was not the case for *MYB96b* (Fig. S10d). Nonetheless, the combined overexpression of *MYB96* and *MYB96b* led to a very small increase in catharanthine levels, which was not observed upon overexpression of *MYB96* alone (Fig. S10g,h). Notably however, because we neither detected any up-regulation of other *vindoline pathway* genes by *MYB96* (Fig. S10d) nor did MYB96 transactivate promoters of other vindoline pathway genes in tobacco protoplasts (Fig.S10c), MYB96 appears to be specific to *T3R*. In an attempt to enhance the vindoline content we further overexpressed *MYB96* in combination with the above-described *MYC2a^D126N^*-*ORCA4* module, which led to a higher availability of the vindoline precursor 16-hydroxytabersonine (Fig. S11). We observed that overexpression of *MYB96*, in addition to the observed up-regulation of *T3R* and *CER1,* also led to a slight but statistically significant up-regulation (around 2-fold) of *stemmadenine pathway* genes and to a significant up-regulation of the catharanthine exporter gene *TPT2* (at least 4-fold) (Fig. S11). However, at the metabolite level, the overexpression of *MYB96* solely or in combination with *ORCA4* and/or *CrMYC2a^D126N^* did not lead to increased vindoline or catharanthine content (Fig. S11d; Table S10). Furthermore, it should be noted that overexpression of *CrMYC2a^D126^* appeared to lead to a down-regulation of most vindoline pathway genes except *T3R* (Fig. S11a), further suggesting that a synergistic/additive action between these TFs seems unlikely.

## DISCUSSION

While *C. roseus* is traditionally known as a medicinal plant, it is also increasingly being established as a model species for plant specialized metabolism in general. Thanks to the extensive work in the past decades by several labs, including ours, our knowledge about the network of MIA pathway branches represents one of the most detailed known biosynthesis networks of specialized metabolites, both at the enzyme and regulator level. In this study, we first aimed to identify new regulators for the different MIA branches, using the currently available pathway information. Thereby, we further underscore the importance of specific ORCA and bHLH TFs as modular regulatory hubs for different MIA pathway branches. While these TF families have been already generally implicated in the regulation of specialized metabolism, our study additionally provides evidence for the functional specialization of TF paralogs due to specific AAR changes in the DBD.

### Functional redundancy and diversification among BIS and ORCA TF clusters

Clusters of TF paralogs appear among various species and families; the orthologs of *ORCA* genes occur for instance also as gene clusters in tobacco, tomato and potato (Shoji *et al.*, 2010; Cárdenas *et al.*, 2016; Kajikawa *et al.*, 2017) and so do the *Arabidopsis* BIS orthologs *bHLH18, 19* and *20* (Cui *et al.*, 2018). Our findings indicate that redundancy and sub-functionalization may occur at the same time for different target genes. In case of the ORCA cluster, we observed that they activate those pathway genes necessary to synthesize common precursors, i.e. “early pathway genes” in a redundant manner. Conversely, later MIA pathway branches are then to some extent controlled by individual members. ORCA3, for instance, is more specific to some of the root MIAs and ORCA4 to the akuammicine branch. ORCA6 seems to have a rather limited impact on known MIA pathway genes, with the exception of a moderate up-regulation of the root MIA genes. The latter results are consistent with a recent study (Singh *et al.*, 2020). These observations are similar to what has been described for the *Medicago truncatula* BIS orthologs, which redundantly activate early triterpene saponin biosynthesis genes but differentially activate downstream branches in the triterpene biosynthesis pathways and/or in different organs (Mertens *et al.*, 2016; Ribeiro *et al.*, 2020).

### Specific AARs in the DBDs of ORCA TFs modulate target gene specificity

Despite increasing knowledge about the DNA binding motifs of different TF families, much less is known about the specific binding affinities of TFs belonging to the same clade. A previous study compared the differential DNA binding affinities of ORCA3 and its orthologs from different species and defined some AARs conferring DNA binding specificity (Shoji *et al.*, 2013). Here, we specifically compare all members of clade XIa AP2/ERFs within the same species under the same experimental conditions and determine AARs conferring specificity that had not been identified previously. It will be interesting to further investigate whether similar AAR changes also confer target gene specialization in the other species with several paralogs and, moreover, determine paralog-specific DNA binding site sequences. Ultimately, further knowledge about such key AARs may offer more precise possibilities for metabolic engineering. For instance, the native version of a given TF might not be the most desirable one for flux into a specific pathway branch but an engineered version in which a key amino acid is substituted might do so. This smallest change for metabolic engineering purposes might be referred to as metabolic editing (Swinnen *et al.*, 2019). Similarly, once the respective DNA binding motifs of a TF are known, specific target genes could be brought under control of that TF by promoter base editing (Swinnen *et al.*, 2016).

### MYC2 – a long-known hub with remaining mechanistic mysteries

As described here and elsewhere, many MIA pathway genes are up-regulated by JA, which can be mimicked by overexpression of the de-repressed *CrMYC2a^D126N^* (Schweizer *et al.*, 2018). In addition, we have previously shown that the combinatorial overexpression with *ORCA3* leads to an up-regulation of the *root MIA pathway* genes and other target genes in a synergistic manner (Schweizer *et al.*, 2018). A similar synergistic up-regulation of target genes has previously been observed between the respective tomato and tobacco MYC2 and AP2/ERF orthologs, suggesting an evolutionary conservation of this combinatorial TF module (De Boer *et al.*, 2011; Cárdenas *et al.*, 2016). Here, we observe that in *C. roseus* the synergistic transactivation is to some extent paralog specific, as the level of synergy is higher in combination with ORCA3 than with ORCA4 for instance. Although we cannot rule out that this is (in part) invoked by different protein stabilities, our results further suggest that this is at least in part due to the AD as we could observe that the AD of ORCA3 significantly enhanced the level of target gene up-regulation when fused to the DBD of ORCA4, whereas the reverse swap rather led to a reduced up-regulation. Although ADs are generally little conserved at the sequence level, it has been shown that specific structural features or subdomains are essential for the TF activity, for instance for the recruitment of the transcriptional machinery (Staby *et al.*, 2017). Noteworthy, *in silico* analyses of the ORCA ADs predict structural differences, which further supports that they are diverse. It remains to be uncovered if and which of these structural differences confer the ORCA3-specific synergistic up-regulation of target genes and if this is for instance mediated by direct physical interaction between the TFs or by other mechanisms. Likewise, more investigation is needed to clarify how the synergistic up-regulation is restricted to specific target genes, i.e. the *root MIA* genes, whereas the up-regulation of other target genes, for instance the BIS TFs, does not appear synergistic.

### Outlook: touching the boundaries of co-expression analysis as a tool to find regulator candidates

Co-expression analysis, which is based on the assumption that the expression profiles of functionally related genes correlate, remains one of the most popular and successful strategies to identify both metabolic pathway genes and regulatory TFs (Movahedi *et al.*, 2012; Goossens, 2015; Wisecaver *et al.*, 2017). However, it has also been noted previously that a successful outcome is greatly dependent on the combination of datasets included in the analysis, the specific methodology used and others (Goossens, 2015; Uygun *et al.*, 2016). In the case of our study, there are a number of conceivable reasons, technical or mechanistic, why we were not able to unambiguously uncover regulators for all MIA branches yet, e.g. the *vindoline pathway*. First, the number of available expression datasets for *C. roseus* is still limited to around 80. Hence, if the pathway branch of interest does not show a (highly) differential expression profile among these datasets, the correlation with expression profiles of regulator candidates might be low. More specifically, the induction of regulatory TFs identified in past studies on *C. roseus* and of their target MIA pathway genes upon JA exposure is highly correlated because of the renowned amplification loops in the JA signaling cascade (De Geyter *et al.*, 2012), which facilitated their discoveries. By contrast, the *vindoline pathway* is not induced upon JA exposure and, apart from a higher expression in green tissues, it is not differentially expressed otherwise. Hence, this pathway branch is either not differentially expressed in response to a specific environmental condition, or a discriminating RNA-Seq dataset for this hypothetical condition is not yet available. Further, transcriptional regulation may involve independently acting TFs individually mediating the specific responses to environment, development, cell type and others. The expression of such a condition-related TF might thus only be strongly correlated to its MIA target genes in a subselection of datasets and not across all datasets. Mining of selected dataset combinations or application of a bi-clustering approach to identify significant co-expression in a subset of samples, could improve this and could be recommended for future co-expression analyses, but would possibly also yield a much longer list of candidates. Next, at the experimental level, it is possible that the candidate list indeed contains positive regulators but that (i) their cloning failed, (ii) they were not sufficiently overexpressed, (iii) their target gene induction required the combined action of TFs (analogous to the ORCA3-CrMYC2a^D126N^ module) but the respective TFs did not happen to be in the same co-overexpression group, (iv) they were not active in flower petals due to epigenetic or post-translational mechanisms.

Our study, together with the vast work previously published by different research groups extensively illustrates that MYC2 TFs globally up-regulate MIA biosynthesis together with the branch-specific ORCA and BIS TFs with the exception of the *vindoline* branch. This raises questions about the biological function of vindoline and/or the intermediates in this pathway branch. While MIA biosynthesis has been generally connected to defense against for instance caterpillars to some extent (Dugé de Bernonville *et al.*, 2017), the chemical ecology of the different specific MIAs is largely unknown. The decoupled expression profile of the *vindoline pathway* from other MIA branches and the absence of up-regulation by CrMYC2a^D126N^ (rather a potential down-regulation) concomitant with a light-induced regulation raise the possibility of other or additional biological functions of vindoline. Nevertheless, the biosynthesis (and/or putative transport) of precursors must be assured, a process that appears not to be light regulated. It is clear that additional studies, likely not based on transcriptome mining solely, are therefore necessary to further investigate how these different signals are integrated to regulate flux through the distinct MIA pathway branches.

## EXPERIMENTAL PROCEDURES

### Compilation and generation of RNA-Seq datasets

In total, 82 RNA-Seq samples were mapped to the latest *C. roseus* draft genome version (Kellner *et al.*, 2015) (http://medicinalplantgenomics.msu.edu/index.shtml) and compiled in our compendium. In addition to publicly available datasets, RNA-Seq data from additional in-house experiments were included (see Table S1). Two datasets consist of cell-type enriched transcriptomes obtained from macro-dissected stem (peeled) and leaf tissue (central vein compared to rest of leaf) as described previously (Van Moerkercke *et al.*, 2015). A third dataset consists of RNA-Seq data from flower petals infiltrated with *Agrobacterium tumefaciens* C58C1 or infiltration buffer as control. The fourth dataset consists of transcriptomes of stably transformed hairy root lines constitutively overexpressing *BIS1*, *ORCA3* or the *green fluorescent protein* (*GFP*) (Van Moerkercke *et al.*, 2015). In all cases, RNA was extracted from three biological replicates, as described previously (Van Moerkercke *et al.*, 2015) and in the Methods section of the main text, and sequenced (50 bp, single-end) with an Illumina HiSeq2500 by GATC Biotech (http://www.gatc-biotech.com/).

### RNA-Seq data processing

RNA-Seq data was obtained from eight publicly available and unpublished datasets, covering 81 samples listed in Table S1. Raw RNA-Seq data was processed with Prose from the Curse toolset (Vaneechoutte and Vandepoele, 2019). Prose uses FastQC (v0.11.7) (Andrews, 2012) and Trimmomatic (v0.38) (Bolger *et al.*, 2014) to perform adapter clipping and quality trimming, and Kallisto (v0.44.0) (Bray *et al.*, 2016) to quantify gene and transcript expression levels in Transcript Per Million (TPM) normalized values. For adapter clipping, Trimmomatic was run with the sliding window size set to 4 and the seed mismatches set to 2. Adapters were detected by FastQC and clipped in each sample separately. For quality trimming, Trimmomatic was run with a simple clip threshold of 10 and a minimum required quality of 15. Reads shorter than 32 bp after quality control were discarded. For Kallisto, an index of the *C. roseus* transcripts was created with a k-mer size of 31. The wrongly split transcript sequence for the *SGD* gene was manually corrected by merging the two CRO_T111319 and CRO_T111320. To process single-end reads with Kallisto, the mean fragment length was set to 200 and the standard deviation to 20. The *C. roseus* expression atlas is available in Supporting Dataset S1. Prose was used to generate k-nearest neighbor (KNN) networks from the expression atlas of 82 samples. In this network, each gene was connected to its top 500 most co-expressing genes. This co-expression information was used for network and heatmap visualization.

### Candidate selection

To search for potential regulators of gene expression, *C. roseus* transcripts were submitted to TRAPID (Van Bel *et al.*, 2013). TRAPID was used to generate functional annotation for these transcripts based on sequence similarity and the presences of protein domains. In total, TRAPID annotated 1,586 genes with the Gene Ontology term ‘regulation of gene expression’ (GO:0010468). Next, 44 genes with known involvement in the catharanthine and vindoline biosynthesis pathways were used as baits to select candidate regulators of these processes. For each bait gene, the top 500 most co-expressing genes in the KNN network were searched for the 1,586 genes that were predicted as regulators of gene expression by TRAPID. Genes that matched this selection were considered to be candidate regulators of the catharanthine or vindoline pathways, since they are both predicted regulators of gene expression and are co-expressed with known pathway genes. If more than 10 matches could be found for a single bait gene, only the top 10 most co-expressing candidates were kept. With 44 bait genes, this could result in at most 440 candidate genes, but due to overlap between the top 10s of different baits, a final list of 111 candidates was selected (Fig. 2a).

### Generation of DNA constructs

Coding sequences (CDSs) of TFs were amplified using cDNA from *C. roseus* var. “Little bright eyes” or *C. roseus* var. “Sunstorm apricot” as templates. Amplification was performed using Q5® High-Fidelity DNA Polymerase (New England BioLabs®) or iProof^TM^ (Bio-Rad) and gene-specific oligonucleotides containing attB Gateway® recombination sites (Table S2). Fragments were gel-purified (GeneJET Gel Extraction Kit, ThermoFisher) and recombined into pDONR^TM^207 using Gateway® BP clonase^TM^ II enzyme mix (ThermoFisher), and the resulting clones sequence verified. ENTRY clones containing *CrMYC2a^D126N^*, *BIS1, BIS2* were cloned previously (Van Moerkercke *et al.*, 2015; Van Moerkercke *et al.*, 2016; Schweizer *et al.*, 2018). ORCA3 protein mutants were constructed by site-directed mutagenesis using iProof^TM^ (Bio-Rad) and pDONR207_CrORCA3 as a template and complementary primers. Chimeric CrORCA3-4 and CrORCA4-3 were constructed by overlap extension PCR using chimeric primers. ENTRY clones containing TFs were recombined using LR Clonase^TM^ II enzyme mix (ThermoFisher) into pH7WG2D or pK7WG2D for overexpression in flower petals, or p2GW7 for promoter transactivation assays in tobacco protoplasts (Karimi *et al.*, 2002).

The promoter fragment of *STR* had been cloned previously (Vom Endt *et al.*, 2007). The promoter fragments of *GO*, *MAT*, *TAT*, *T3O*, *T3R*, *D4H* and *DAT* were amplified from *C. roseus* var. “Sunstorm apricot” genomic DNA using specific oligonucleotides (Table S3), recombined into pDONR^TM^207 and sequence verified, as described for the TF CDSs. The resulting ENTRY clones containing promoter fragments were recombined into pGWL7 (Karimi *et al.*, 2002) for transient expression assays in tobacco protoplasts.

### Transient expression assays in *Nicotiana tabacum* protoplasts

Promoter transactivation of *C. roseus* gene promoters by *C. roseus* TFs was measured via transient expression assays in *N. tabacum* ‘Bright Yellow-2’ protoplasts exactly as described previously (Vanden Bossche *et al.*, 2013; Schweizer *et al.*, 2018). Statistical analysis (ANOVA with Tukey’s test for multiple comparisons) was performed using PRISM GraphPad 8 software.

### Plant material and flower petal transformation

*Catharanthus roseus* var. “Little bright eyes” plants were grown in the greenhouse under 16-h/8-h light/dark conditions. Plants of the same age (6 months to 2 years) were used for flower petal infiltration with *A. tumefaciens* C58C1 harboring overexpression constructs as previously described (Schweizer *et al.*, 2018) with minor adaptations. Briefly, all opened flowers were removed two days prior to infiltration. *A. tumefaciens* pre-cultures were inoculated from glycerol stocks two days prior to infiltration to inoculate final overnight cultures one day prior to infiltration. Cultures were washed and re-suspended as described (Schweizer *et al.*, 2018). The suspensions were diluted and mixed (in case of combined overexpression of up to four TF overexpression constructs) to obtain a final concentration of OD_600_=0.4. Infiltration took place as described using four to five flowers, each from different individual plants. Samples were harvested in four biological replicates as described (Schweizer *et al.*, 2018). In case of parallel analysis of gene expression and metabolite profiling, each sample was divided in two immediately before freezing.

### Gene expression analysis by qPCR

RNA extraction and reverse transcription were performed as described (Schweizer *et al.*, 2018) with modifications. Up to 500 ng of RNA was used for reverse transcription using either the iScript^TM^ cDNA synthesis kit (Bio-Rad) or the qScript cDNA Synthesis Kit (Quantabio). qPCR was performed using a JANUS pipetting robot (PerkinElmer) and a LightCycler® 480 (Roche) using SYBR Green qPCR master Mix (Agilent) and gene-specific oligonucleotides (Table S5) in two technical replicates. The data was normalized to *N2227* and *SAND* as reference genes (Pollier *et al.*, 2014) using qBase (Hellemans *et al.*, 2007). Statistical analyses (ANOVA with Tukey’s test for multiple comparisons or Student’s *t*-test, as indicated) were performed on log2-transformed data using PRISM GraphPad 8 software. Principle component analyses (PCAs) of MIA target gene profiles were performed with ClustVis (Metsalu and Vilo, 2015).

### Metabolite profiling

For LC-MS, 10 μL of the sample was injected in an Acquity UPLC BEH C18 column (150×2.1 mm, 1.7 μm; Waters, Milford, MA) mounted on an Acquity UPLC system (Waters) connected to a Synapt HDMS qTOF-MS (Micromass) with an electrospray ionization source. A gradient was run at flow rate of 350 µL/min using acidified (0.1% (v/v) formic acid) solvents A (water/acetonitrile (99:1, v/v)) and B (acetonitrile/water; 99:1, v/v)): time 0 min, 5% B; 30 min, 50% B; 33 min, 100% B. The mass spectrometer was set to positive ionization mode with the following parameter values: capillary voltage, 2.5 kV; sampling cone, 40 V; extraction cone, 4 V; source temperature, 120°C; desolvation temperature, 400°C; cone gas flow, 50 L/h; and desolvation gas flow, 750 L/h. Centroid data were recorded between *m*/*z* 100 and *m*/*z* 1,200 at a scan speed of 0.2 s/scan. Peak areas were determined using Progenesis QI software (Waters). PCAs were performed with MetaboAnalyst 4.0 (https://www.metaboanalyst.ca/) using Pareto-scaled mass spectrometry data with standard settings (Chong *et al.*, 2018). Statistical analyses of identified MIAs (ANOVA with Tukey’s test for multiple comparisons) were performed using PRISM GraphPad 8 software. Identifications of the MIAs were done using FT-MS and MS*^n^* data generated as described (Schweizer *et al.*, 2018) and are shown in Supporting dataset S2.

### Gene IDs and data deposition

Gene IDs (*C. roseus* genome version 2) are listed with the corresponding primer sequences in Table S4. In some cases, the cloned sequences differed from publicly available genome and transcriptome data. All sequences cloned for this study have been deposited in the GenBank database; accession numbers are also listed in Table S4.

## ACKNOWLEDGEMENTS

We thank Robin Vanden Bossche for excellent technical assistance, and Alex Van Moerkercke for generating the RNA-Seq data of transformed *C. roseus* hairy root lines. M.C. and F.S. are indebted to the Swiss National Science Foundation (P300PA_177831 and P300PA_167772). This work was supported by European Union’s Horizon 2020 research and innovation program under Grant Agreement No 760331 (Newcotiana). This work was supported by the Agency for Innovation by Science and Technology (IWT) in Flanders (predoctoral fellowship to D.V.). J.G.G. is indebted to the Portuguese Fundação para a Ciência e Tecnologia (FCT) and FEDER Funds (SFRH/BD/97590/2013).

## AUTHOR CONTRIBUTIONS

M.C. and A.G. designed the research. M.C. performed the research and data analysis with contributions from D.M., F.S, L.D.M., R.D.C, J.G.G, T.M.-C. F.J.M.H and M.S.. D.V. and K.V. performed RNA-Seq data and co-expression analysis and inferred candidate regulators. J.P. collected and analyzed metabolite data and performed compound identification. M.C. and A.G. wrote the manuscript with consent from all other authors.

## SUPPORTING INFORMATION

**Fig. S1** MIA gene expression levels upon bulk overexpression of TF candidates.

**Fig. S2** Separate overexpression of positive TF candidates.

**Fig. S3** Promoter transactivation of iridoid pathway genes by combinations of BIS.

**Fig. S4** Comparison of structural features and motifs of ORCA activation domains (ADs).

**Fig. S5** Expression profiles of MIA biosynthesis genes and regulators in analyzed RNA-seq data.

**Fig. S6** MIA levels upon overexpression of *ORCAs*.

**Fig. S7** MIA gene expression levels upon overexpression of *ORCA3* single mutants.

**Fig. S8** MIA levels upon overexpression of *ORCA3* mutants.

**Fig. S9** MIA levels upon combinatorial overexpression of *CrMYC2a^D126N^* and *ORCAs*.

**Fig. S10** Orthologs of cuticular wax regulators up-regulate the vindoline pathway gene *T3R*.

**Fig. S11** Combinatorial overexpression of *ORCA4, CrMYB96, CrMYC2a^D126N^*.

**Table S1** List of RNA-Seq datasets used.

**Table S2** Oligonucleotide primers used for cloning of coding sequences.

**Table S3** Oligonucleotide primers used for cloning of promoter fragments.

**Table S4** List of all cloned TFs and promoter fragments and Genebank IDs.

**Table S5** Oligonucleotide primers used for qPCR.

**Table S6** MIA levels upon overexpression of *ORCAs*.

**Table S7** MIA levels upon overexpression of *ORCA3* mutants.

**Table S8** MIA levels upon combinatorial overexpression of *CrMYC2a^D126N^* and *ORCAs*.

**Table S9** MIA levels upon overexpression of *MYB96/b*.

**Table S10** MIA levels upon overexpression of *MYB96/b, CrMYC2a^D126N^* and *ORCA4*

**Supporting Dataset I** *C. roseus* expression atlas.

**Supporting Dataset II** MIAs identified in this study.

