## Supporting Dataset II for "Subfunctionalization of paralog transcription factors contributes to regulation of alkaloid pathway branch choice in *Catharanthus roseus*"

**Supplemental Dataset 1. Identification of *C. roseus* alkaloids.**

Metabolites identified in *C. roseus* flowers by LC-FT-ICR-MS. For each metabolite it is indicated whether the metabolite was identified based on an authentic standard or whether it was tentatively identified based on its accurate mass and MS$^n$ fragmentation.

| Compound | Identification | Fragmentation spectra |
|---|---|---|
| Catharanthine | Standard | Schweizer *et al.*, 2018 |
| Vindoline | Standard | Schweizer *et al.*, 2018 |
| Strictosidine | Standard | Schweizer *et al.*, 2018 |
| Secologanin | Standard | Schweizer *et al.*, 2018 |
| 16-Hydroxytabersonine | Standard | Schweizer *et al.*, 2018 |
| 19-Hydroxytabersonine | Standard | Schweizer *et al.*, 2018 |
| Hörhammericine | Standard | Schweizer *et al.*, 2018 |
| Vinblastine | Standard | Schweizer *et al.*, 2018 |
| Vincristine | Standard | Schweizer *et al.*, 2018 |
| Akuammicine | Standard | Schweizer *et al.*, 2018 |
| Vincadifformine | Standard | Schweizer *et al.*, 2018 |
| Loganin | Standard | Schweizer *et al.*, 2018 |
| Isositsirikine | Standard | Schweizer *et al.*, 2018 |
| 16-Methoxyhörhammericine | Predicted | Schweizer *et al.*, 2018 |
| 16-Hydroxyhörhammericine | Predicted | Schweizer *et al.*, 2018 |
| Anhydrovinblastine | Predicted | Schweizer *et al.*, 2018 |
| Geissoschizine | Predicted | Schweizer *et al.*, 2018 |
| Serpentine | Predicted | Schweizer *et al.*, 2018 |
| 16-Hydroxyvincadifformine | Predicted | Schweizer *et al.*, 2018 |
| Perivine | Predicted | Schweizer *et al.*, 2018 |
| *O*-acetylstemmadenine | Predicted | Schweizer *et al.*, 2018 |
| Minovincinine | Predicted | Schweizer *et al.*, 2018 |
| Unknown 1 (isositsirikine isomer) | Predicted | Schweizer *et al.*, 2018 |
| Unknown 2 (strictosidine aglycone isomer) | Predicted | Schweizer *et al.*, 2018 |
| 16-hydroxy-19-*O*-acetyltabersonine | Predicted | Schweizer *et al.*, 2018 |
| Strictosidine secologanoside | Predicted | Schweizer *et al.*, 2018 |
| Deacetylvindoline | Predicted | This study |
| Desacetoxyvindoline | Predicted | This study |
| Demethoxyvindoline = vindorosine | Predicted | This study |
| Desacetoxyvindorosine | Predicted | This study |
| Deacetylvindorosine = catharosine | Predicted | This study |
| Strictosidinic Acid | Predicted | This study |
| 19-*O*-acetyltabersonine | Predicted | This study |

**Supplemental Dataset 1. Identification of *C. roseus* alkaloids.**

Metabolites identified in *C. roseus* flowers by LC-FT-ICR-MS. For each metabolite it is indicated whether the metabolite was identified based on an authentic standard or whether it was tentatively identified based on its accurate mass and $MS^n$ fragmentation.

| Compound | Identification | Fragmentation spectra |
|---|---|---|
| 16-Hydroxylochnericine | Predicted | This study |
| 16-Hydroxylochnericine glucoside | Predicted | This study |
| Unknown MIA (geissoschizine isomer 1) | Predicted | This study |
| Vandrikidine | Predicted | This study |
| Unknown MIA (geissoschizine isomer 2) | Predicted | This study |
| 16-Hydroxytabersonine glucoside | Predicted | This study |
| Unknown MIA (catharanthine isomer 1) | Predicted | This study |
| Unknown MIA (catharanthine isomer 2) | Predicted | This study |
| Unknown MIA (catharanthine isomer 3) | Predicted | This study |
| Unknown MIA (serpentine isomer) | Predicted | This study |

Below, for each metabolite identified in this study a detailed description of its identification is given.

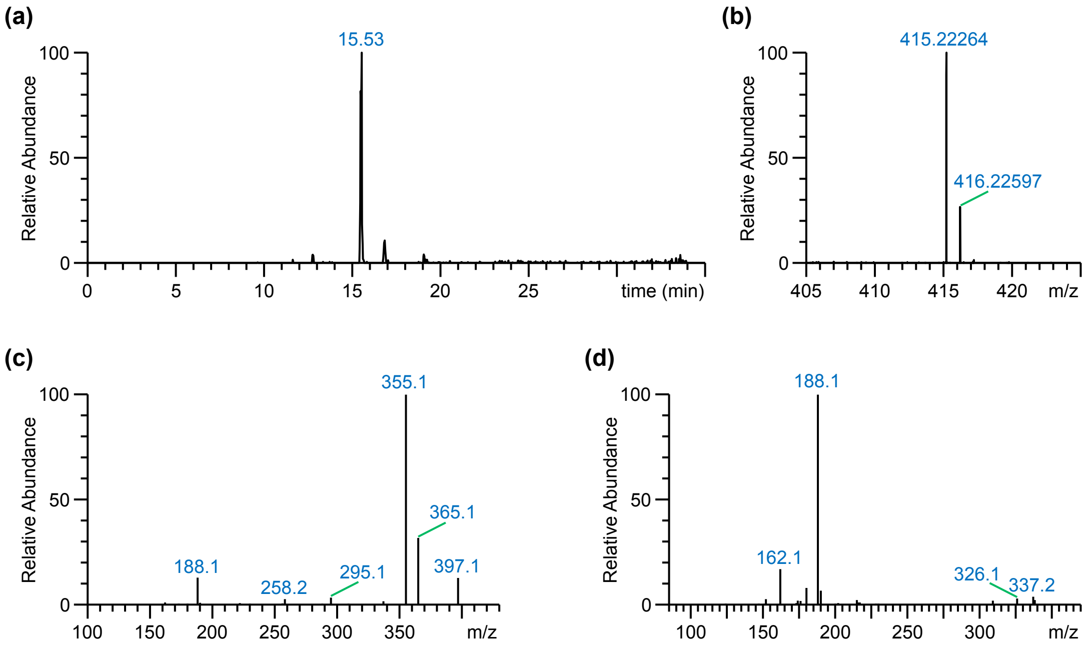


**Figure 1.** Identification of deacetylvindoline. (**a**) Extracted Ion Current (EIC = 415.22 ± 0.01 Da) chromatogram of a *C. roseus* flower sample. Only one major peak with a mass corresponding to the mass of deacetylvindoline was observed. (**b**) FT-MS scan of the compound eluting at 15.53 min revealed an accurate mass of the $[M+H]^+$ ion at *m*/*z* 415.22264, corresponding to the chemical formula of deacetylvindoline, $C_{23}H_{30}N_2O_5$ (δ ppm = -0.261). (**c**) $MS^2$ fragmentation spectrum of the $[M+H]^+$ ion at *m*/*z* 415.22. Major daughter ions are observed at *m*/*z* 397 (loss of $H_2O$), *m*/*z* 365 (loss of $H_2O$ + MeOH), *m*/*z* 355 (loss of methyl formate), and *m*/*z* 188. (**d**) $MS^3$ fragmentation spectrum of the daughter ion at *m*/*z* 355.

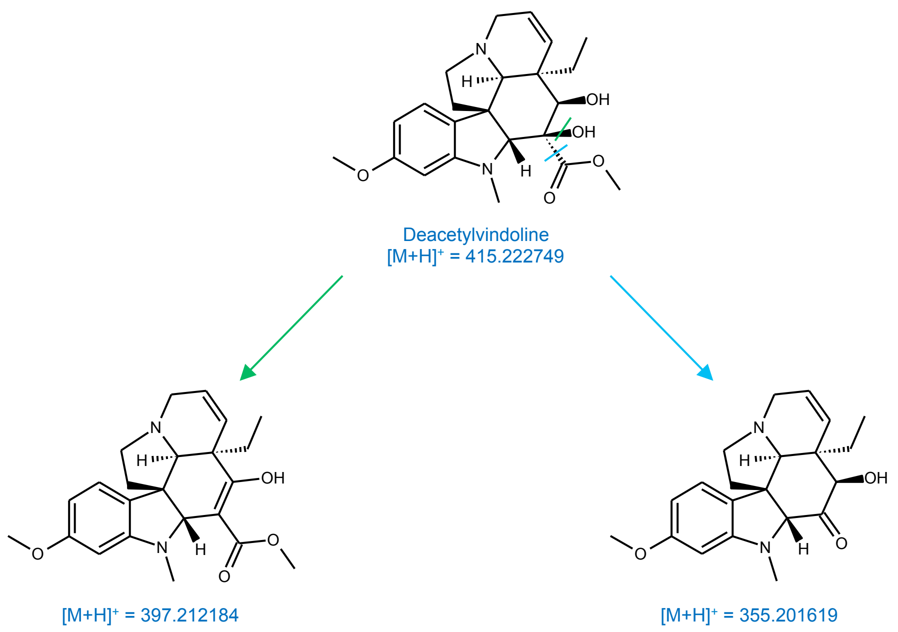


**Figure 2.** Chemical structure and proposed fragmentation of deacetylvindoline.

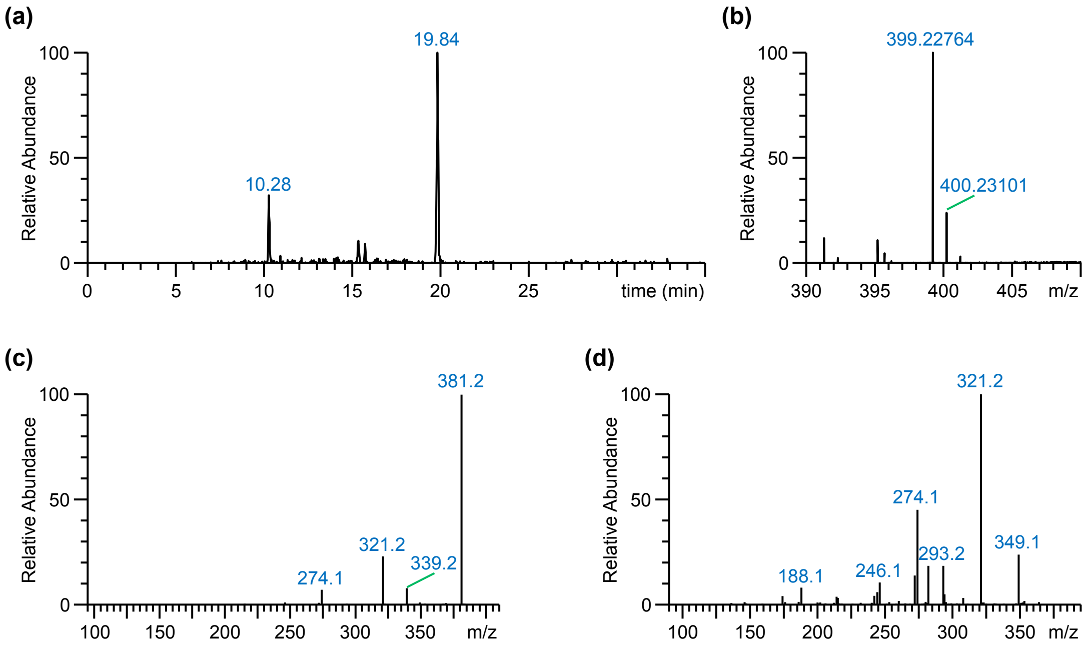


**Figure 3.** Identification of desacetoxyvindoline. (**a**) Extracted Ion Current (EIC = 399.22 ± 0.01 Da) chromatogram of a *C. roseus* flower sample. Two major peaks with this mass were observed in the chromatogram; the first peak eluting at 10.28 min was previously identified as 16-methoxyhörhammericine (Schweizer et al., 2018). (**b**) FT-MS scan of the compound eluting at 19.84 min revealed an accurate mass of the $[M+H]^+$ ion at *m*/*z* 399.22764, corresponding to the chemical formula of desacetoxyvindoline, $C_{23}H_{30}N_2O_4$ ($\delta$ ppm = -0.486). (**c**) $MS^2$ fragmentation spectrum of the $[M+H]^+$ ion at *m*/*z* 399.23. Major daughter ions are observed at *m*/*z* 381 (loss of $H_2O$), and *m*/*z* 321 (loss of $H_2O$ + methyl formate). (**d**) $MS^3$ fragmentation spectrum of the daughter ion at *m*/*z* 381.

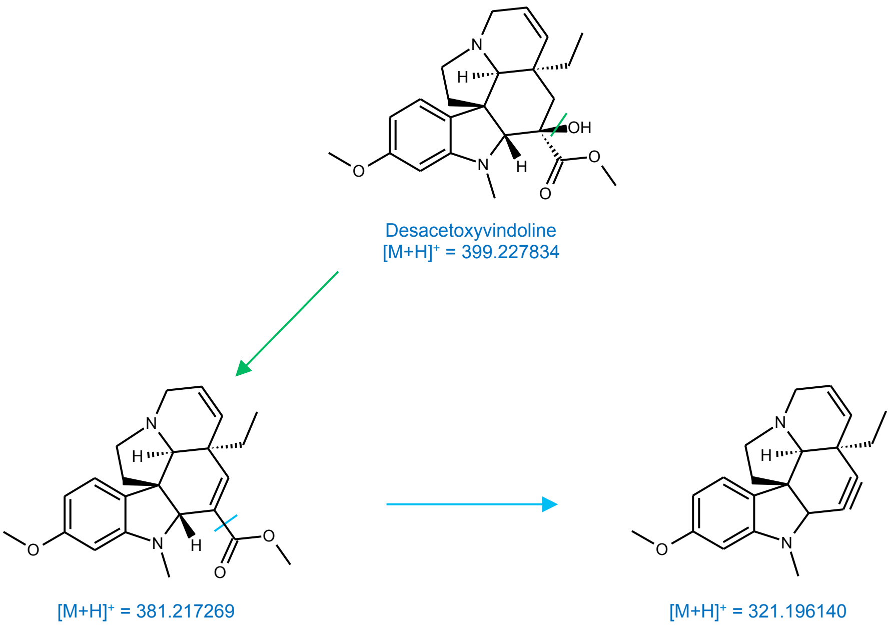


**Figure 4.** Chemical structure and proposed fragmentation of desacetoxyvindoline.

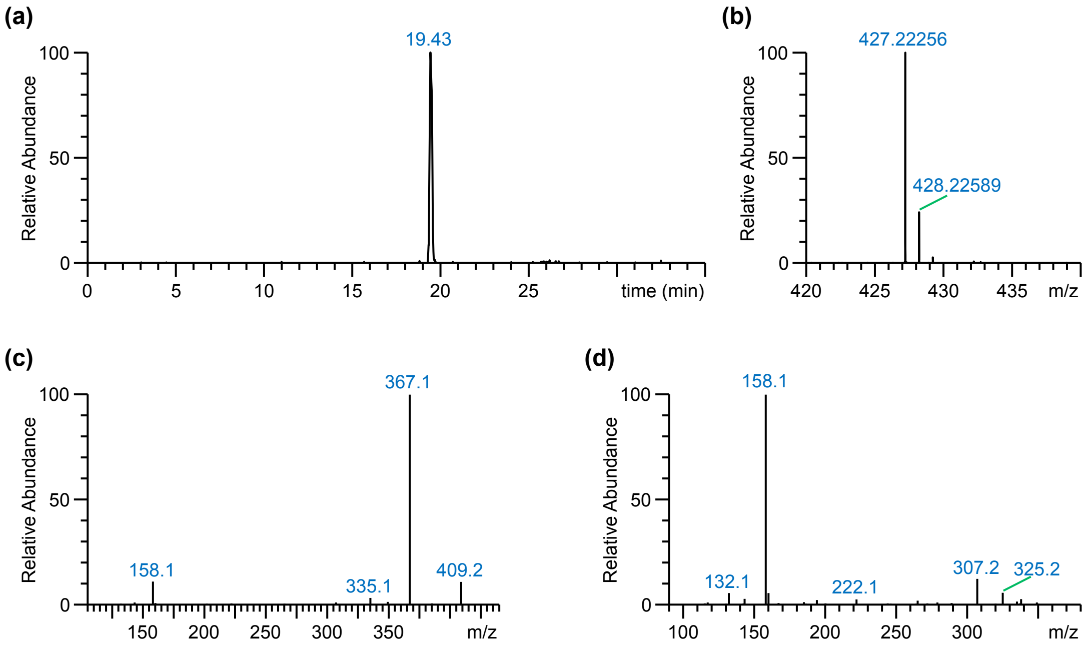


**Figure 5.** Identification of vindorosine. (**a**) Extracted Ion Current (EIC = 427.22 ± 0.01 Da) chromatogram of a *C. roseus* flower sample. Only one major peak with a mass corresponding to the mass of vindorosine was observed. (**b**) FT-MS scan of the compound eluting at 19.43 min revealed an accurate mass of the $[M+H]^+$ ion at *m*/*z* 427.22256, corresponding to the chemical formula of vindorosine, $C_{24}H_{30}N_2O_5$ (δ ppm = -0.441). (**c**) $MS^2$ fragmentation spectrum of the $[M+H]^+$ ion at *m*/*z* 427.22. Major daughter ions are observed at *m*/*z* 409 (loss of MeOH), *m*/*z* 367 (loss of acetate), and *m*/*z* 158. These fragments ions are similar to those of vindoline (Schweizer et al., 2018), which has an additional *O*-methyl group compared to vindorosine, leading to a difference of 30 amu for all fragment ions. (**d**) $MS^3$ fragmentation spectrum of the daughter ion at *m*/*z* 367.

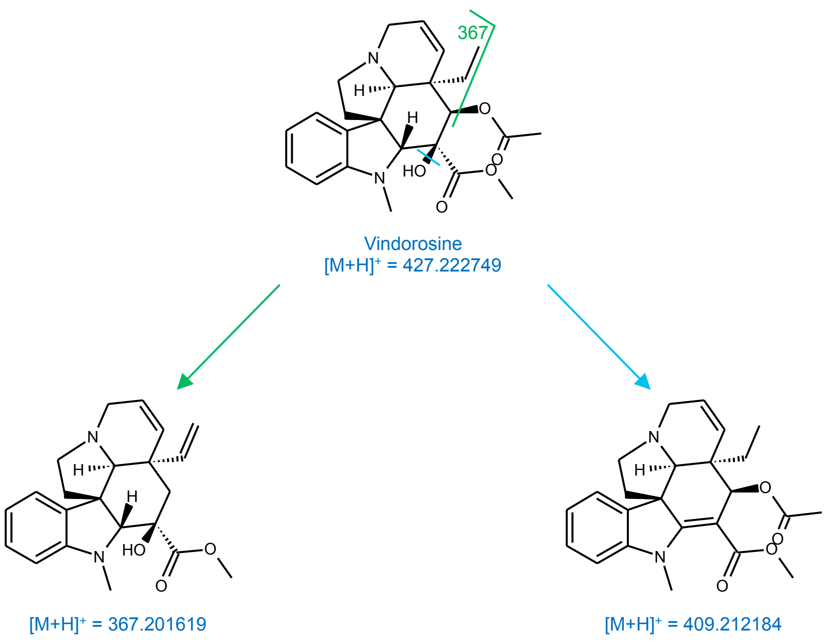


**Figure 6.** Chemical structure and proposed fragmentation of vindorosine.

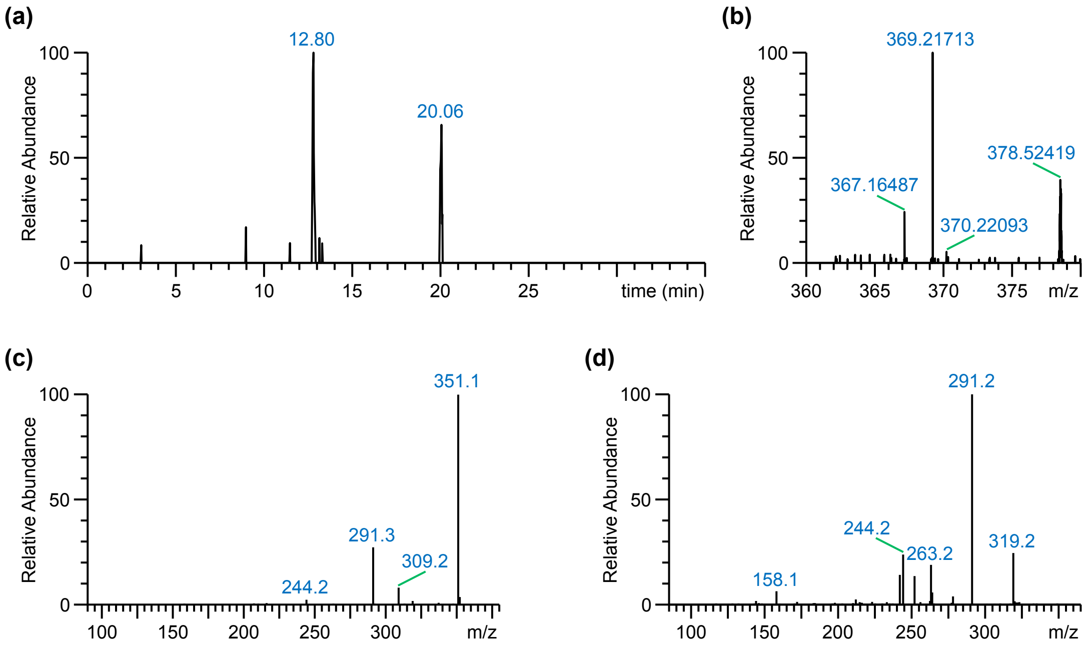


**Figure 7.** Identification of desacetoxyvindorosine. (**a**) Extracted Ion Current (EIC = 369.22 ± 0.01 Da) chromatogram of a *C. roseus* flower sample. Two low abundant peaks with this mass were observed in the chromatogram. As the very similar desacetoxyvindoline elutes at 19.84 min, the compound eluting at 20.06 minutes is our prime candidate. (**b**) FT-MS scan of the compound eluting at 20.06 min revealed an accurate mass of the $[M+H]^+$ ion at *m*/*z* 369.21713, corresponding to the chemical formula of desacetoxyvindorosine, $C_{22}H_{28}N_2O_3$ (δ ppm = -0.377). (**c**) $MS^2$ fragmentation spectrum of the $[M+H]^+$ ion at *m*/*z* 369.22. Major daughter ions are observed at *m*/*z* 351 (loss of $H_2O$) and *m*/*z* 291 (loss of $H_2O$ + methyl formate). (**d**) $MS^3$ fragmentation spectrum of the daughter ion at *m*/*z* 351.

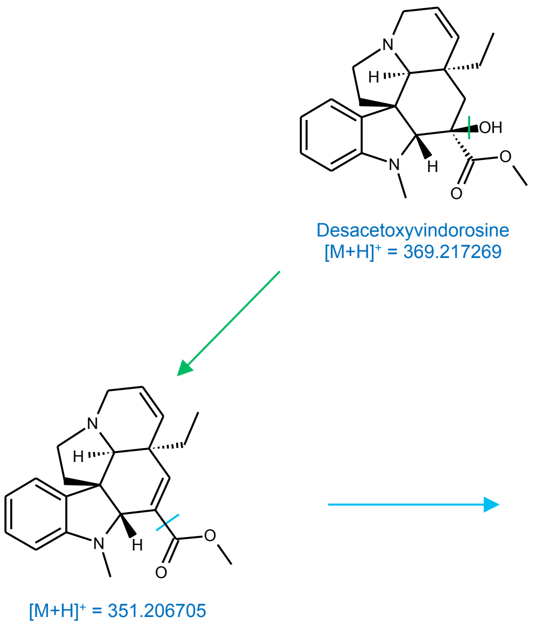


**Figure 8.** Chemical structure and proposed fragmentation of desacetoxyvindorosine.

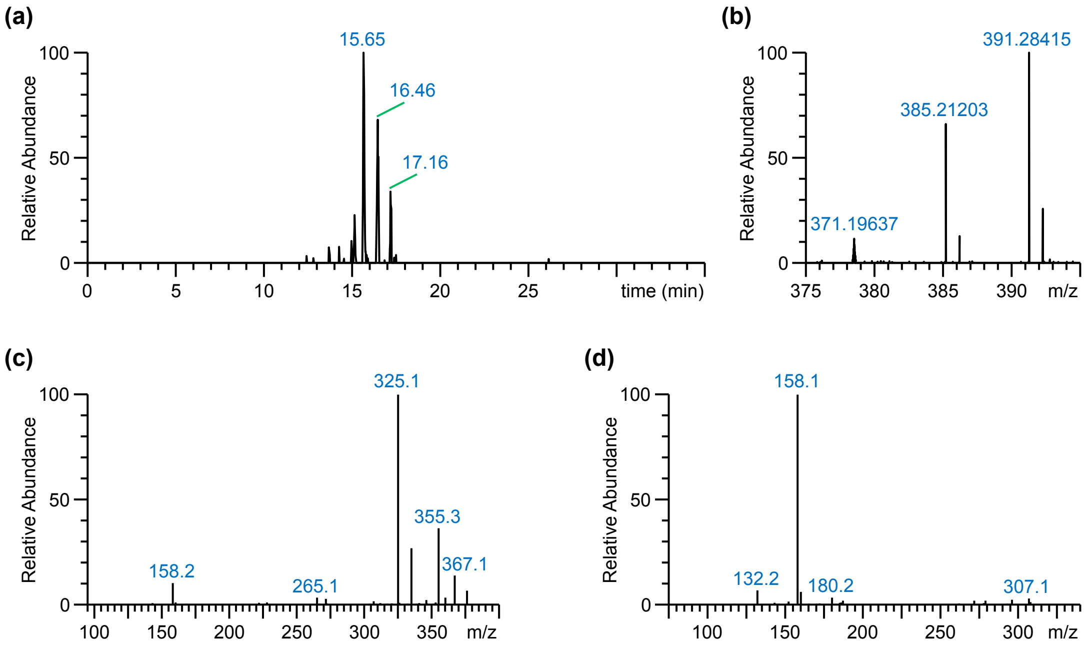


**Figure 9.** Identification of deacetylvindorosine. (**a**) Extracted Ion Current (EIC = 385.21 ± 0.01 Da) chromatogram of a *C. roseus* flower sample. A few low abundant peaks with this mass were observed in the chromatogram. As the very similar deacetylvindoline elutes at 15.53 min, the compound eluting at 15.65 minutes is our prime candidate. (**b**) FT-MS scan of the compound eluting at 15.65 min revealed an accurate mass of the $[M+H]^+$ ion at *m*/*z* 385.21201, corresponding to the chemical formula of deacetylvindorosine, $C_{22}H_{28}N_2O_4$ (δ ppm = -0.451). (**c**) $MS^2$ fragmentation spectrum of the $[M+H]^+$ ion at *m*/*z* 385.21. Major daughter ions are observed at *m*/*z* 355 and *m*/*z* 325 (loss of methyl formate). (**d**) $MS^3$ fragmentation spectrum of the daughter ion at *m*/*z* 325.

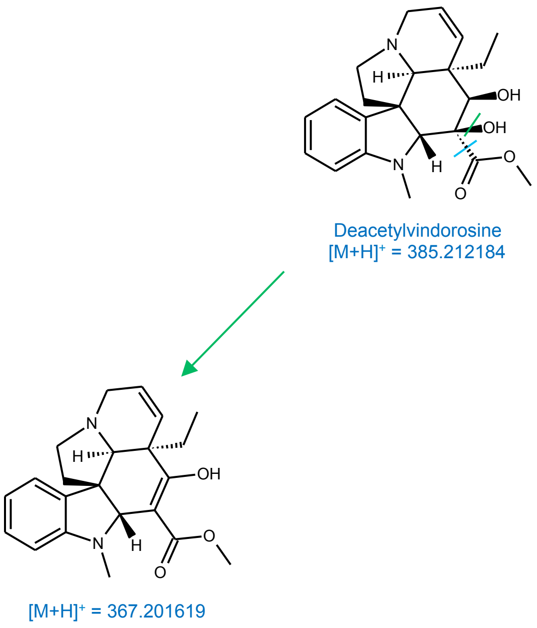


**Figure 10.** Chemical structure and proposed fragmentation of deacetylvindorosine.

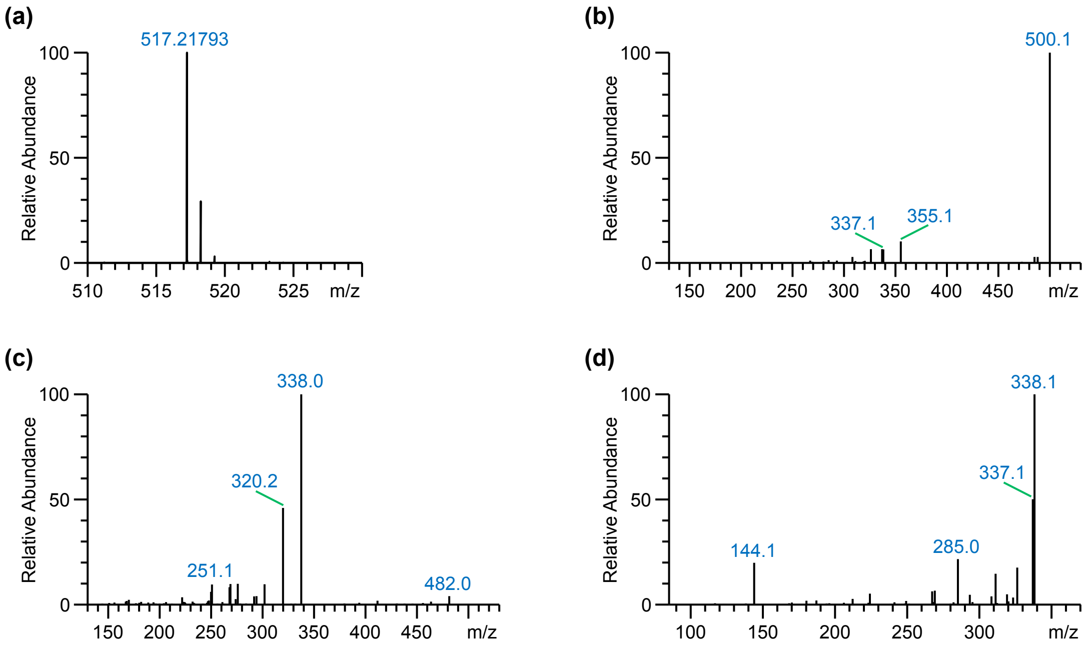


**Figure 11.** Identification of strictosidinic acid. (**a**) FT-MS scan of the unknown peak revealed an accurate mass of the $[M+H]^+$ ion at *m/z* 517.21793, corresponding to the chemical formula $C_{26}H_{32}N_2O_9$ (δ ppm = -0.246). (**b**) $MS^2$ fragmentation spectrum of the ion at *m/z* 517.22. The major daughter ion observed at *m/z* 500 arises from the loss of an $NH_3$ molecule (17 Da), a neutral loss previously observed for strictosidine (Schweizer et al., 2018). Additional fragment ions were observed at *m/z* 355 and *m/z* 337, resulting from the loss of a hexose moiety (162 Da), or a hexose and a $H_2O$ molecule (162 + 18 Da), respectively. (**c**) $MS^3$ fragmentation spectrum of the daughter ion at *m/z* 500. In analogy with the fragmentation of strictosidine, major granddaughter ions were observed at *m/z* 338 and *m/z* 320, resulting from the loss of a hexose moiety (162 Da), or a hexose and a $H_2O$ molecule (162 + 18 Da), respectively. (**d**) $MS^3$ fragmentation spectrum of the daughter ion at *m/z* 355. Major granddaughter ions were observed at *m/z* 338 (loss of $NH_3$), *m/z* 337 (loss of $H_2O$), and *m/z* 144, the latter corresponding to an indole molecule.

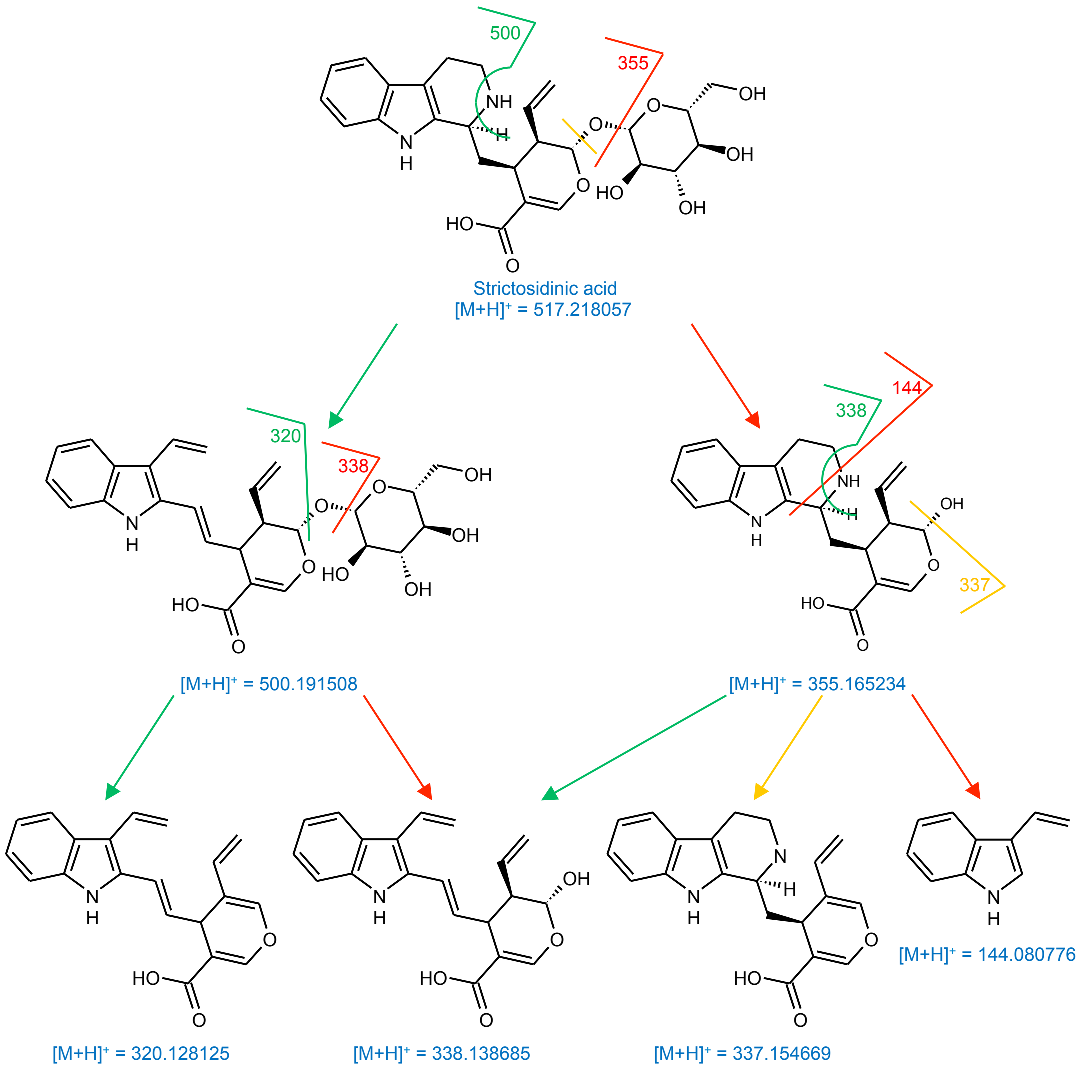


**Figure 12.** Chemical structure and proposed fragmentation of strictosidinic acid.

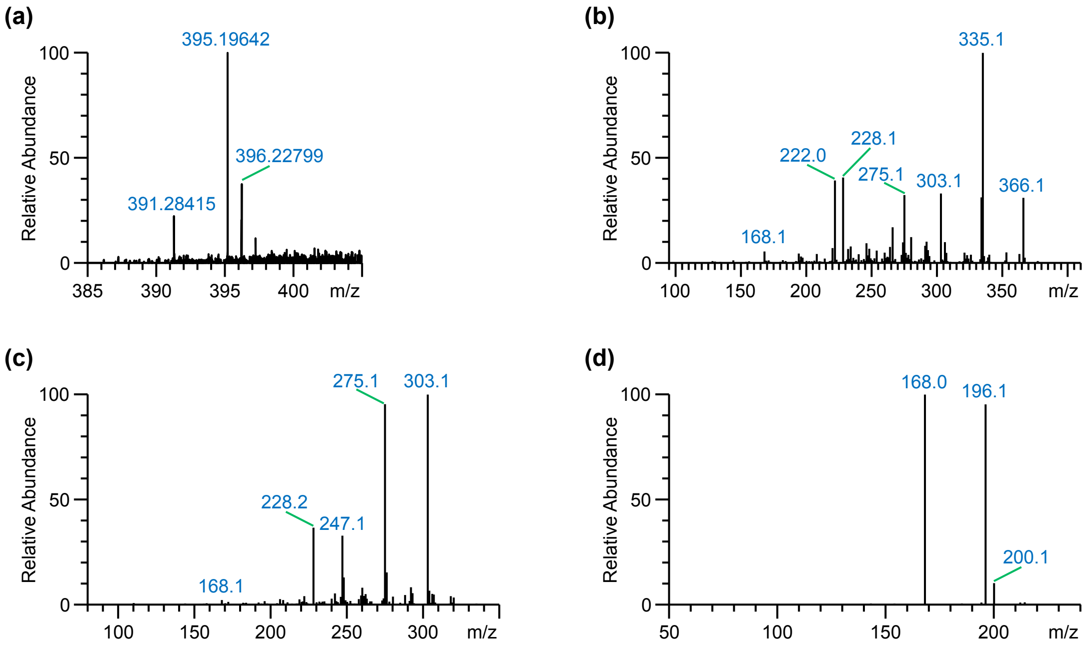


**Figure 13.** Identification of 19-*O*-acetyltabersonine. (**a**) FT-MS scan of the unknown peak revealed an accurate mass of the $[M+H]^+$ ion at *m*/*z* 395.19642, corresponding to the chemical formula $C_{23}H_{26}N_2O_4$ (δ ppm = -0.288). (**b**) $MS^2$ fragmentation spectrum of the ion at *m*/*z* 395.20. The major daughter ion observed at *m*/*z* 335 arises from the loss of either an acetate moiety, which would point to an acetylated molecule, or the loss of methyl formate. Additional daughter ions are observed at *m*/*z* 366, *m*/*z* 303, *m*/*z* 275, *m*/*z* 228, and *m*/*z* 222. (**c**) $MS^3$ fragmentation spectrum of the daughter ion at *m*/*z* 335. Major granddaughter ions are observed at *m*/*z* 303 (loss of MeOH, and indicating an O-methyl group), *m*/*z* 275 (loss of methyl formate), *m*/*z* 247, and *m*/*z* 228. (**d**) $MS^3$ fragmentation spectrum of the daughter ion at *m*/*z* 228. Two major granddaughter ions were observed at *m*/*z* 168, and *m*/*z* 196. The occurrence of a daughter ion at *m*/*z* 228 and its $MS^3$ fragmentation into these two major ions was also observed for tabersonine (Schweizer et al., 2018). Combined with the observed loss of an acetate moiety, this fragmentation indicates the unknown metabolite corresponds to 19-*O*-acetyltabersonine.

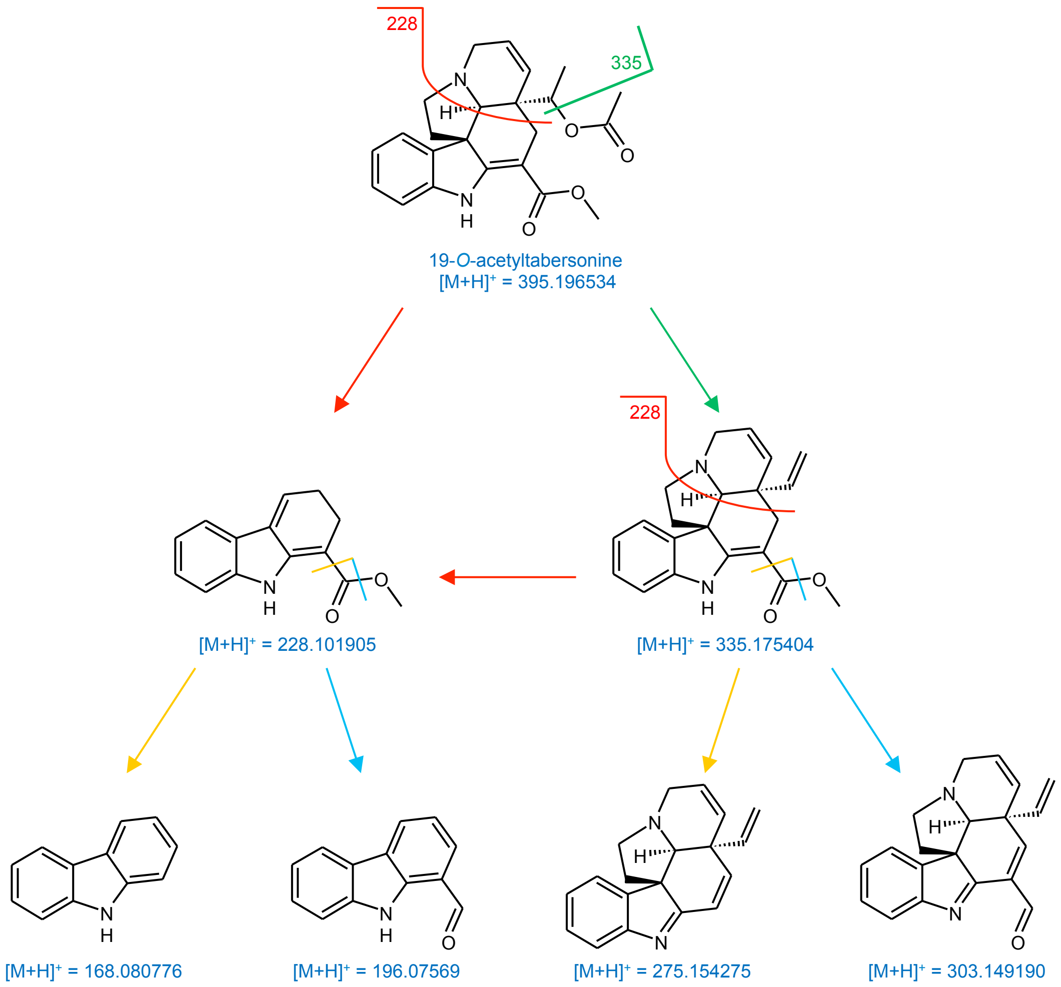


**Figure 14.** Chemical structure and proposed fragmentation of 19-*O*-acetyltabersonine.

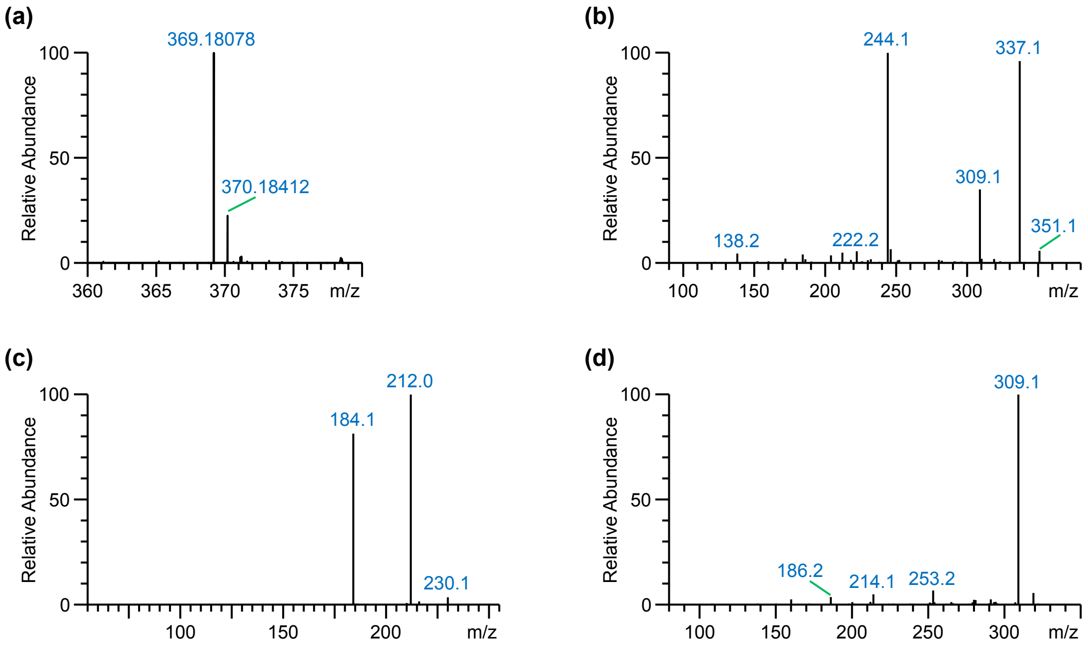


**Figure 15.** Identification of 16-hydroxylochnericine. (**a**) FT-MS scan of the unknown peak revealed an accurate mass of the $[M+H]^+$ ion at *m*/*z* 369.18078, corresponding to the chemical formula $C_{21}H_{24}N_2O_4$ (δ ppm = -0.281). (**b**) $MS^2$ fragmentation spectrum of the $[M+H]^+$ ion at *m*/*z* 369.18. The fragmentation spectrum is similar to that of lochnericine, which has major daughter ions at *m*/*z* 228, *m*/*z* 293, and *m*/*z* 321 (Schweizer et al., 2018). The ions observed for this unknown metabolite are 16 Da heavier, pointing to a lochnericine-like molecule with an additional hydroxy group. (**c**) $MS^3$ fragmentation spectrum of the daughter ion at *m*/*z* 244. Two major granddaughter ions were observed at *m*/*z* 184 and *m*/*z* 212. The occurrence of a daughter ion at *m*/*z* 244 and its $MS^3$ fragmentation into these two major granddaughter ions was also observed for 16-hydroxytabersonine (Schweizer et al., 2018). As such, the unknown molecule is likely lochnericine, with an additional hydroxy group at the C-16 position: 16-hydroxylochnericine (**d**) $MS^3$ fragmentation spectrum of the daughter ion at *m*/*z* 337.

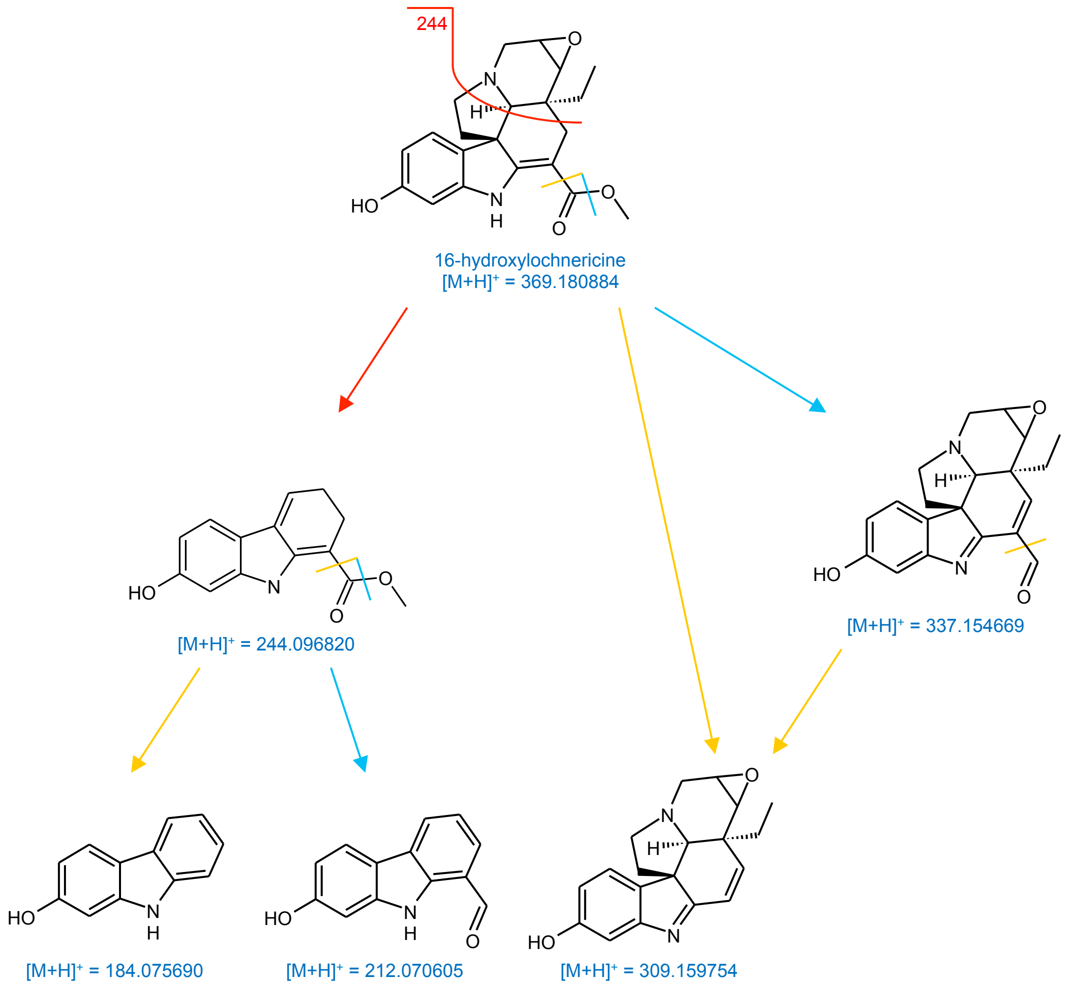


**Figure 16.** Chemical structure and proposed fragmentation of 16-hydroxylochnericine.

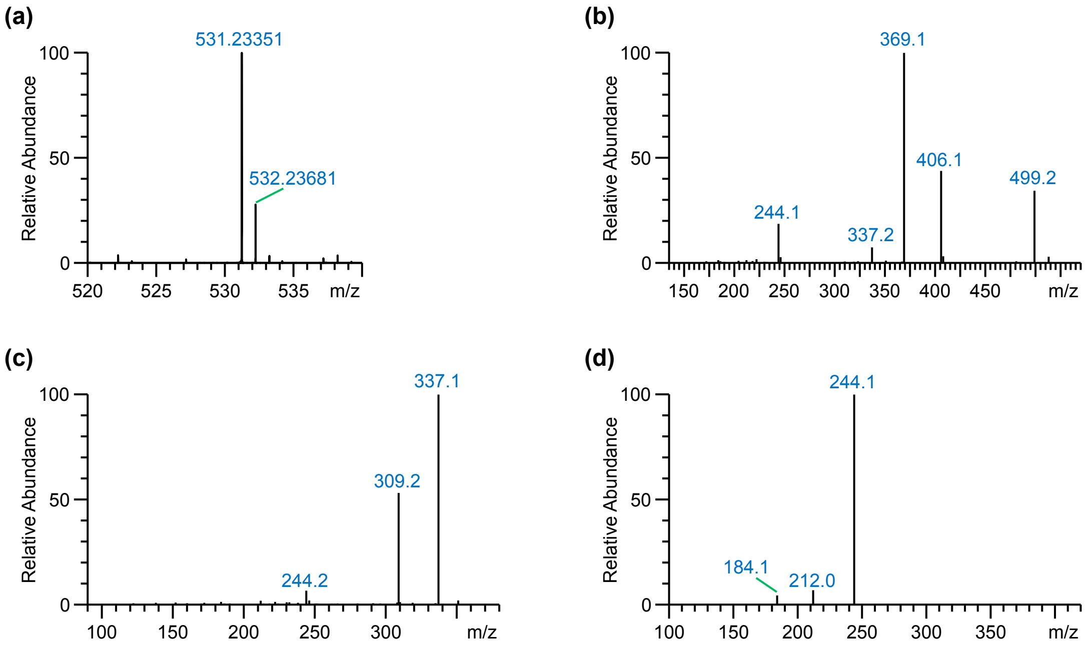


**Figure 17.** Identification of 16-hydroxylochnericine glucoside. (**a**) FT-MS scan of the unknown peak revealed an accurate mass of the $[M+H]^+$ ion at *m*/*z* 531.23351, corresponding to the chemical formula $C_{27}H_{34}N_2O_9$ (δ ppm = -0.371). (**b**) $MS^2$ fragmentation spectrum of the $[M+H]^+$ ion at *m*/*z* 531.23. The major daughter ion at *m*/*z* 369 points to the loss of a hexose residue. Other major daughter ions are observed at *m*/*z* 499 (loss of MeOH), *m*/*z* 406, *m*/*z* 337, and *m*/*z* 244. This fragmentation pattern indicates the unknown metabolite corresponds to a MIA glucoside, with the MIA having a $[M+H]^+$ of 369.18 Da. (**c**) $MS^3$ fragmentation spectrum of the daughter ion at *m*/*z* 369. (**d**) $MS^3$ fragmentation spectrum of the daughter ion at *m*/*z* 406. the presence of a granddaughter ion at m/z 244 and two minor granddaughter ions at *m*/*z* 184 and *m*/*z* 212 indicates a 16-hydroxylochnericine backbone. As such, the unknown glucoside likely corresponds to 16-hydroxylochnericine glucoside.

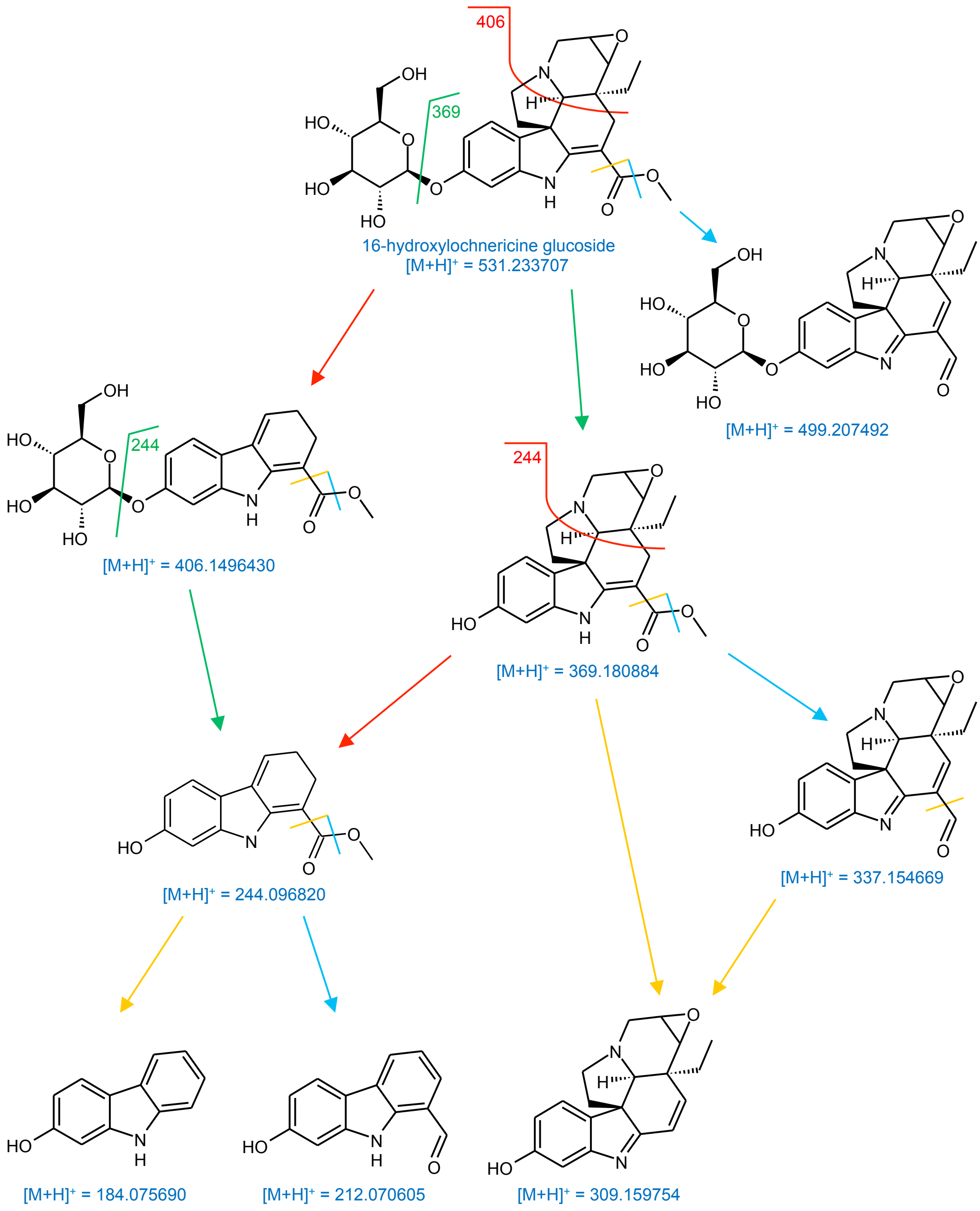


**Figure 18.** Chemical structure and proposed fragmentation of 16-hydroxylochnericine glucoside.

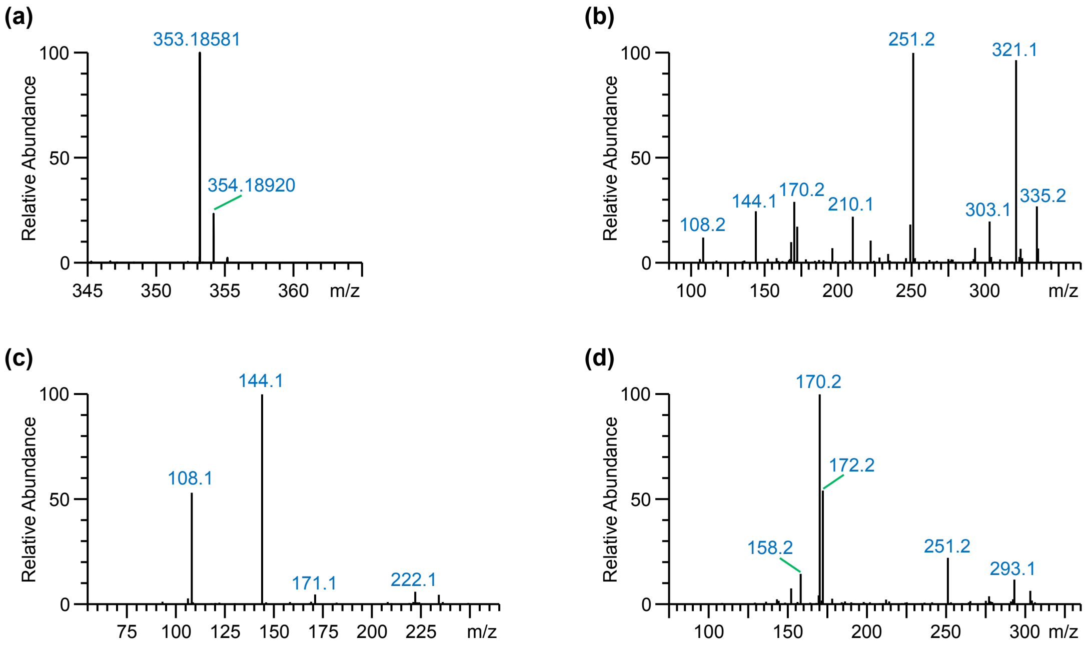


**Figure 19.** Identification of an unknown MIA (geissoschizine isomer 1). (**a**) FT-MS scan of the unknown peak revealed an accurate mass of the $[M+H]^+$ ion at *m/z* 353.18581, corresponding to the chemical formula $C_{21}H_{24}N_2O_3$ ($\delta$ ppm = -0.451). (**b**) $MS^2$ fragmentation spectrum of the $[M+H]^+$ ion at *m/z* 353.19. Major daughter ions are observed at *m/z* 321 (loss of MeOH) and *m/z* 251. A major daughter ion at *m/z* 251 was also observed for isositsirikine and geissoschizine (Schweizer et al., 2018), indicating the unknown metabolite likely corresponds to a corynanthe-type MIA with an open E-ring. (**d**) $MS^3$ fragmentation spectrum of the daughter ion at *m/z* 251. Like for isositsirikine (Schweizer et al., 2018), fragmentation of this ion leads to two major granddaughter ions at *m/z* 108 and *m/z* 144. (**d**) $MS^3$ fragmentation spectrum of the daughter ion at *m/z* 321.

Based on this fragmentation spectrum, we propose the unknown metabolite to be an isomer of geissoschizine with an open E-ring. This is further supported by the observed loss of a $H_2O$ molecule in the $MS^2$ fragmentation spectrum, which indicates the presence of a free hydroxy group. The isomer proposed in Figure 20, 19*Z*-geissoschizine, is only illustrative for the fragmentation of the unknown metabolite.

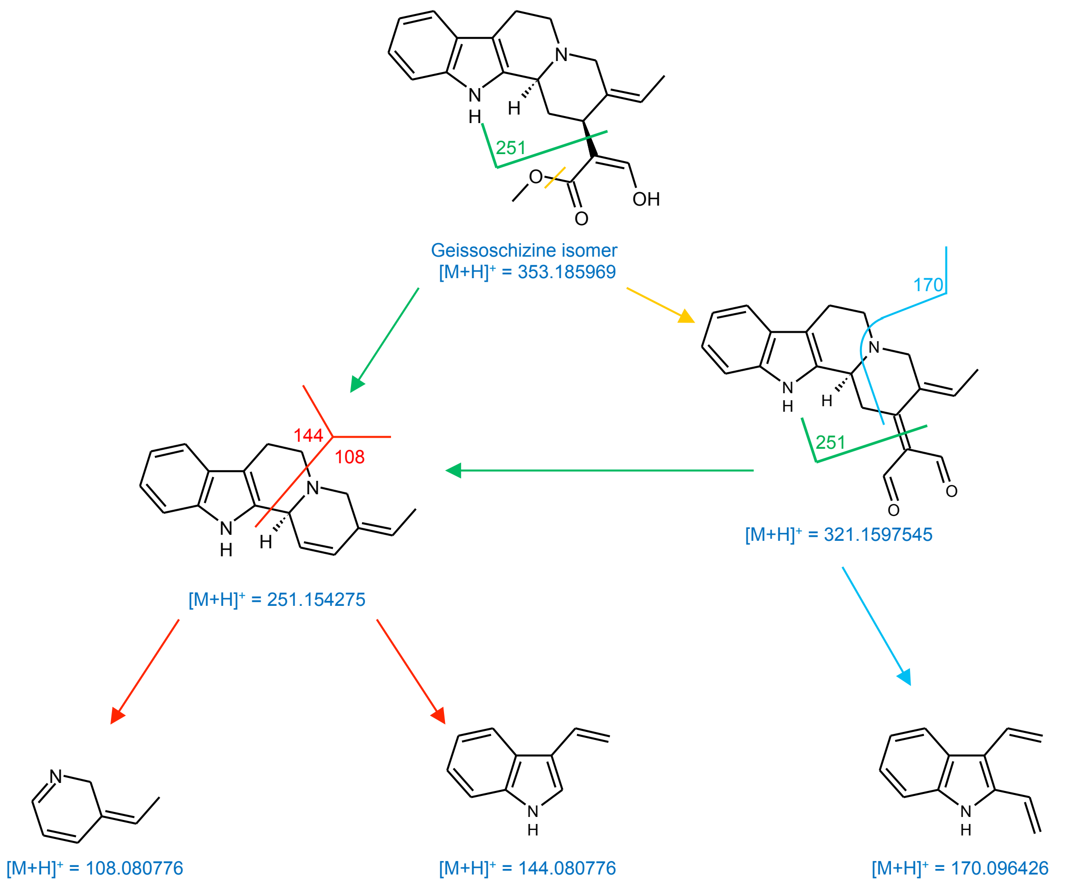


**Figure 20.** Proposed fragmentation of the unknown MIA (geissochizine isomer 1). The isomer depicted here, 19*Z*-geissoschizine, is only illustrative for the possible fragmentation of the unknown metabolite.

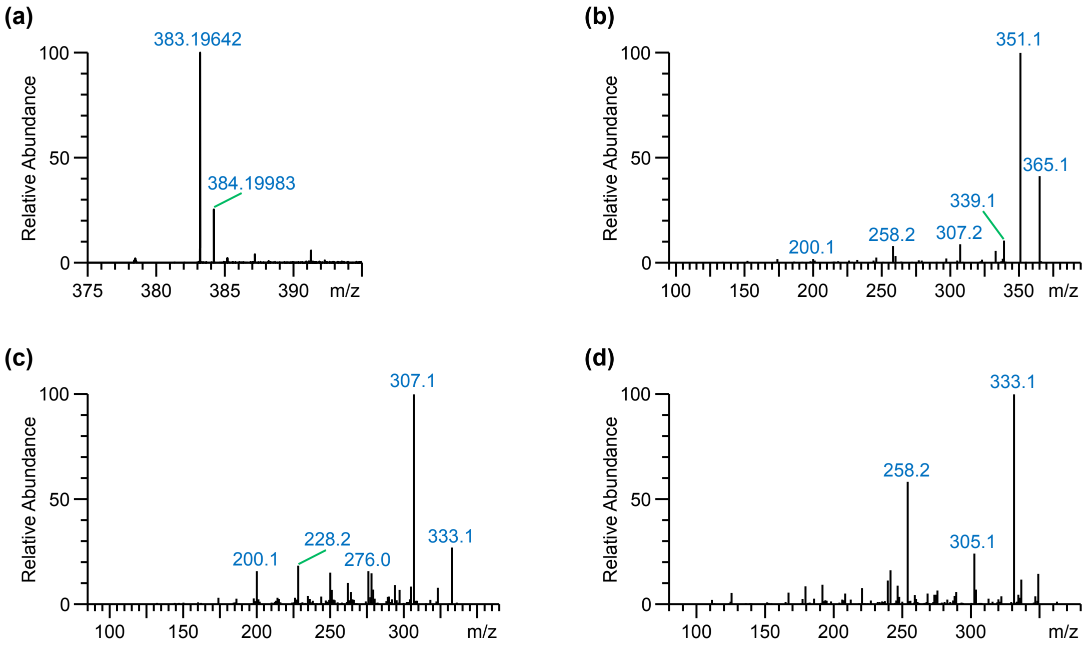


**Figure 21.** Identification of vandrikidine. (**a**) FT-MS scan of the unknown peak revealed an accurate mass of the $[M+H]^+$ ion at *m*/*z* 383.19642, corresponding to the chemical formula $C_{22}H_{26}N_2O_4$ ($\delta$ ppm = -0.297). (**b**) $MS^2$ fragmentation spectrum of the $[M+H]^+$ ion at *m*/*z* 383.20. Major daughter ions are observed at *m*/*z* 365 (loss of $H_2O$) and *m*/*z* 351 (loss of MeOH). A less abundant daughter ion was observed at *m*/*z* 258, a daughter ion previously observed for 16-methoxyhörhammericine (Schweizer et al., 2018), suggesting an aspidosperma-type MIA backbone with a methoxy group at the C-16 position as is the case for vandrikidine. (**c**) $MS^3$ fragmentation spectrum of the daughter ion at *m*/*z* 351. (**d**) $MS^3$ fragmentation spectrum of the daughter ion at *m*/*z* 365. Both $MS^3$ fragmentation spectra support that the unknown metabolite corresponds to vandrikidine.

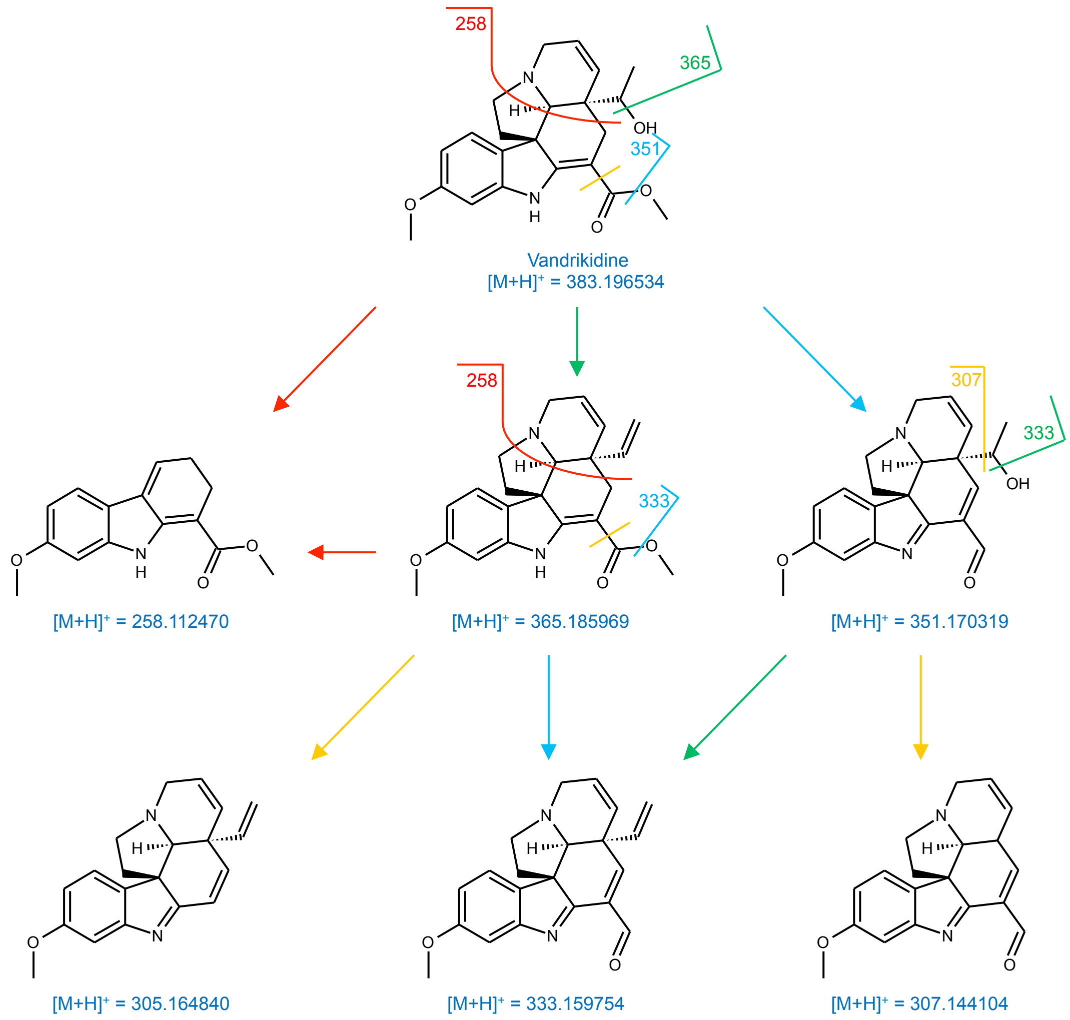


**Figure 22.** Chemical structure and proposed fragmentation of vandrikidine.

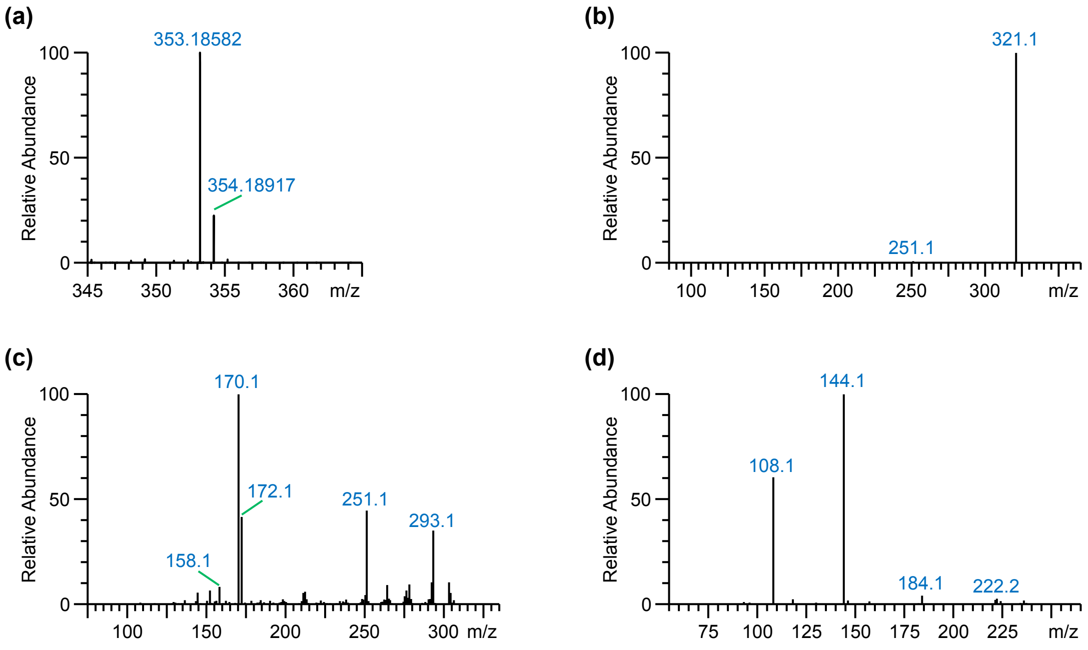


**Figure 23.** Identification of an unknown MIA (geissoschizine isomer 2). (**a**) FT-MS scan of the unknown peak revealed an accurate mass of the $[M+H]^+$ ion at *m*/*z* 353.18582, corresponding to the chemical formula $C_{21}H_{24}N_2O_3$ ($\delta$ ppm = -0.422). (**b**) $MS^2$ fragmentation spectrum of the $[M+H]^+$ ion at *m*/*z* 353.19. A single major daughter ion was observed at *m*/*z* 321 (loss of MeOH). A minor ion was observed at *m*/*z* 251. (**c**) $MS^3$ fragmentation spectrum of the daughter ion at *m*/*z* 321. (**d**) $MS^3$ fragmentation spectrum of the daughter ion at *m*/*z* 251 yields two major granddaughter ions at *m*/*z* 108 and *m*/*z* 144.

Fragmentation of a daughter ion at *m*/*z* 251 into two major granddaughter ions at *m*/*z* 108 and *m*/*z* 144 was observed before for isositsirikine. Hence, it is likely that the unknown metabolite also corresponds to a corynanthe-type MIA with an open E-ring. Since only one major fragment ion, resulting from the loss of a MeOH moiety was observed in the $MS^2$ fragmentation spectrum, only one functional group that can be easily lost upon collision induced dissociation is present on the molecule. To illustrate the fragmentation of the unknown geissoschizine isomer, a methoxylated dioxolene was used (Figure 24).

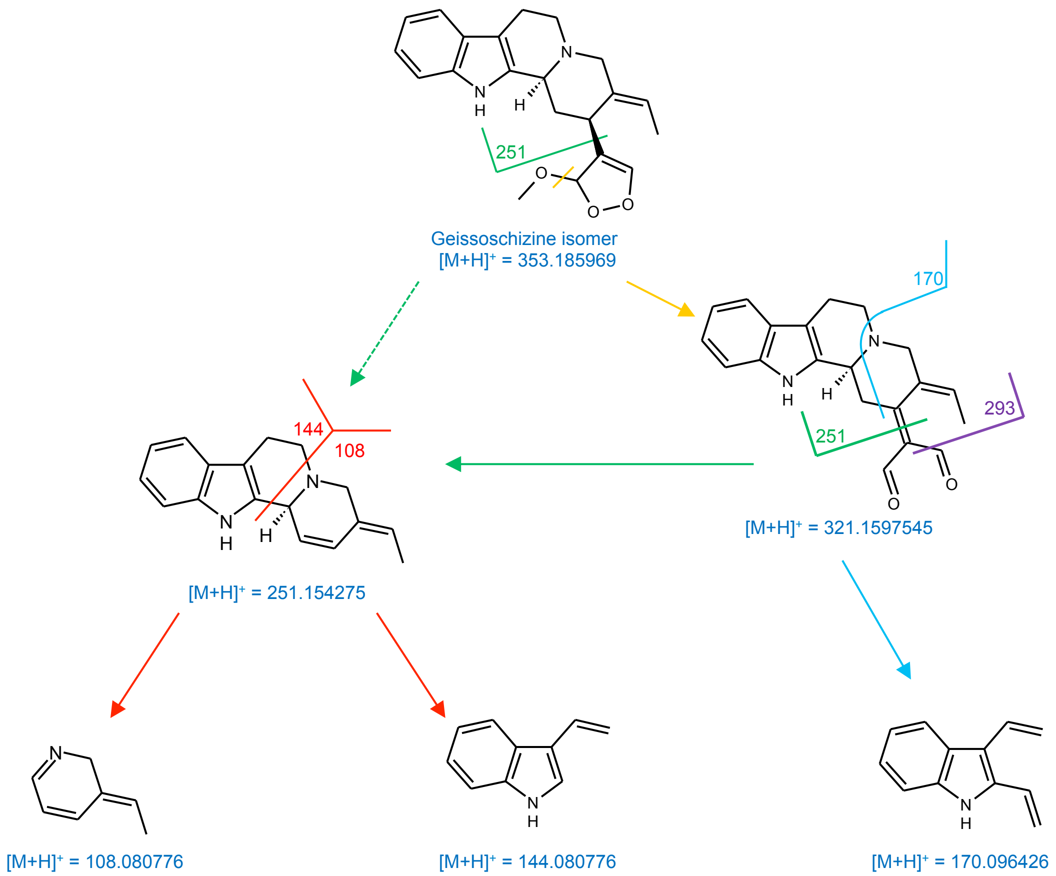


**Figure 24.** Proposed fragmentation of the unknown MIA (geissochizine isomer 2). The isomer depicted here is only illustrative for the possible fragmentation of the unknown metabolite.

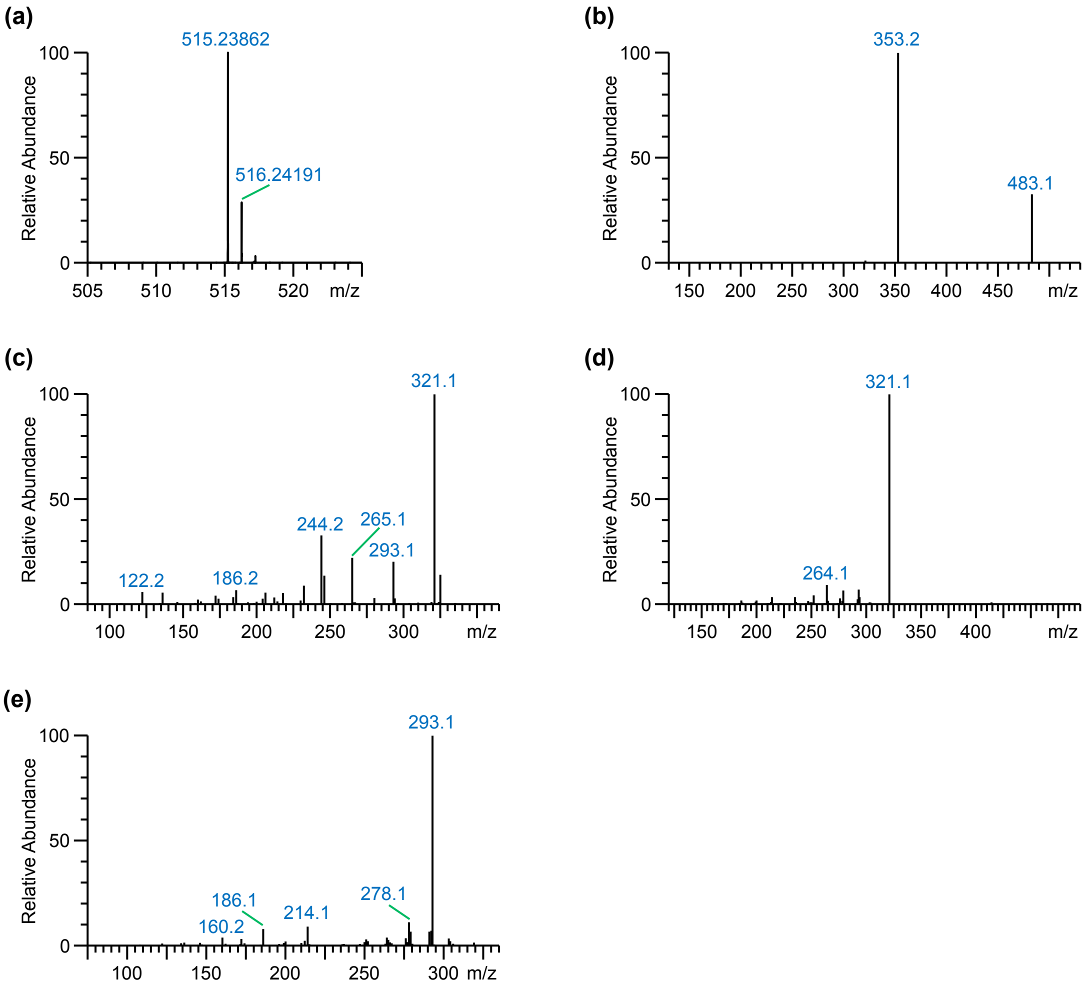


**Figure 25.** Identification of 16-hydroxytabersonine glucoside. (**a**) FT-MS scan of the unknown peak revealed an accurate mass of the $[M+H]^+$ ion at *m*/*z* 515.23862, corresponding to the chemical formula $C_{27}H_{34}N_2O_8$ ($\delta$ ppm = -0.335). (**b**) $MS^2$ fragmentation spectrum of the ion at *m*/*z* 515.24. The major daughter ion observed at *m*/*z* 353 arises from the loss of a hexose moiety, whereas the ion observed at *m*/*z* 483 is derived from the neutral loss of MeOH. Together this indicates the unknown compound is a glycoside of a MIA with an *O*-methyl group. (**c**) $MS^3$ fragmentation spectrum of the daughter ion at *m*/*z* 353. Major granddaughter ions are observed at *m*/*z* 321 (loss of MeOH, and again indicating an *O*-methyl group), *m*/*z* 293 (loss of methyl formate), *m*/*z* 265, and *m*/*z* 244. (**d**) $MS^3$ fragmentation spectrum of the daughter ion at *m*/*z* 483. The major granddaughter ion observed at *m*/*z* 321 results from the loss of a hexose moiety. (**e**) $MS^4$ fragmentation spectrum of the granddaughter ion at *m*/*z* 321. This $MS^4$ fragmentation spectrum is identical to that of the $MS^3$ fragmentation spectrum of the daughter ion at *m*/*z* 321 of 16-hydroxytabersonine (Schweizer et al., 2018). This indicates the unknown metabolite is likely a glucoside of 16-hydroxytabersonine.

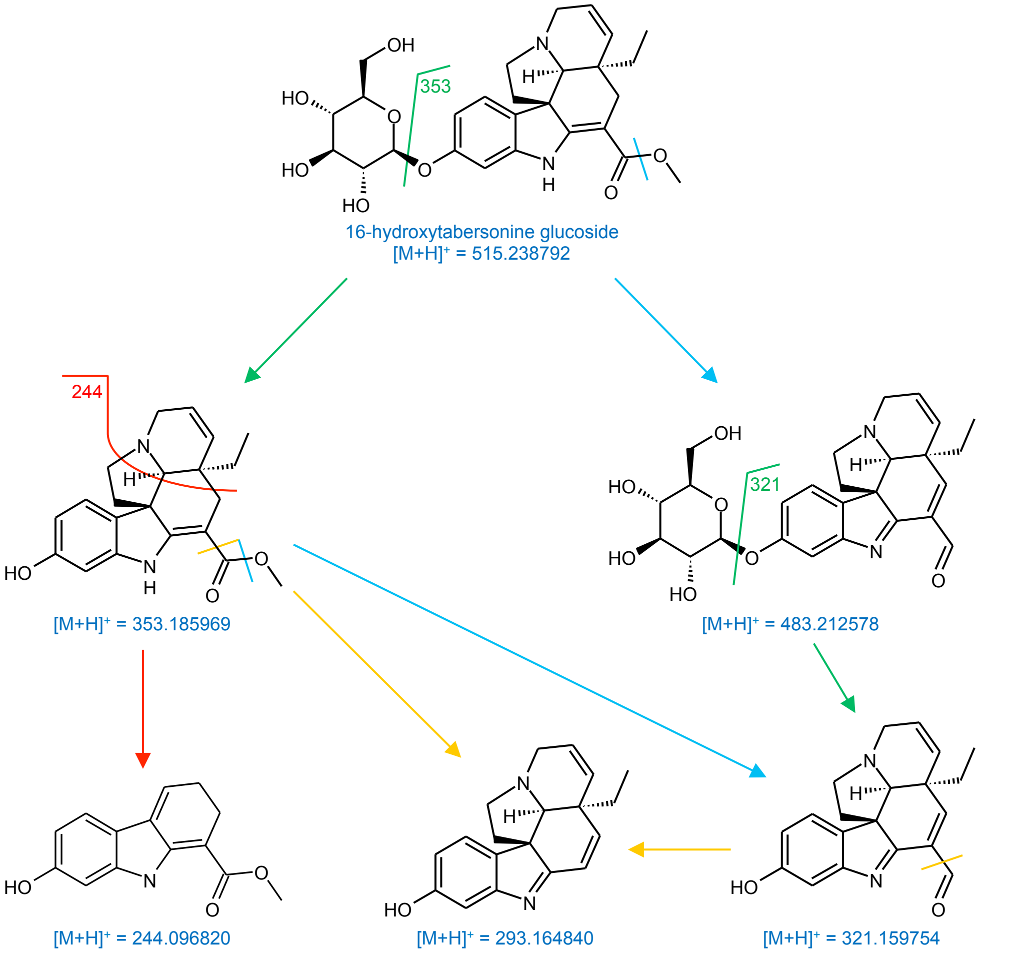


**Figure 26.** Chemical structure and proposed fragmentation of 16-hydroxytabersonine glucoside.

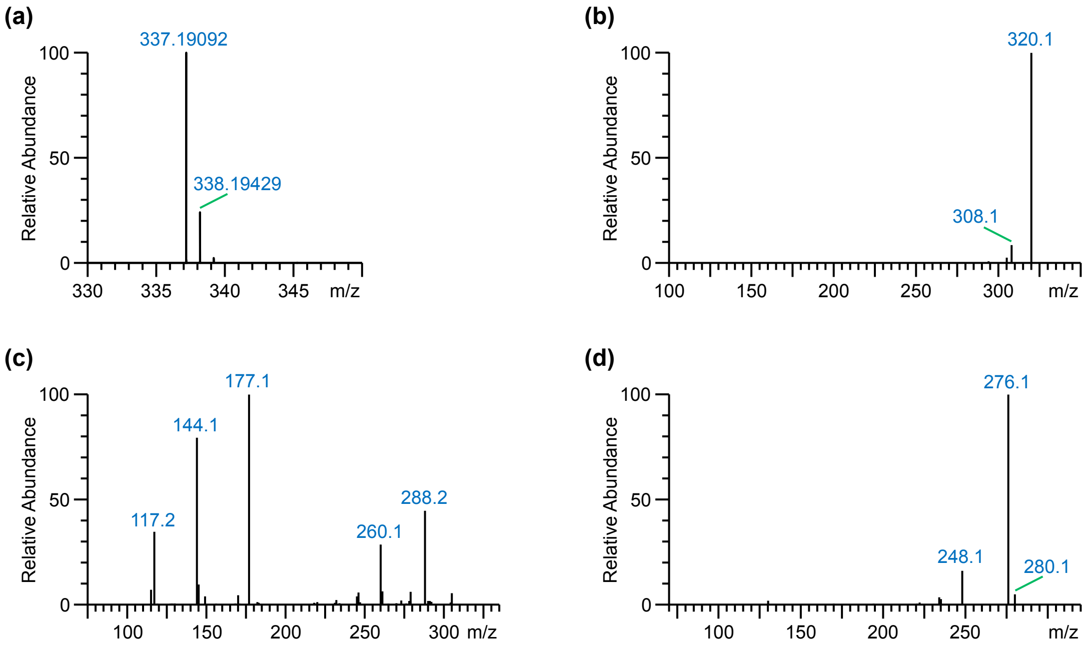


**Figure 27.** Identification of an unknown MIA (catharanthine isomer 1). (**a**) FT-MS scan of the unknown peak revealed an accurate mass of the $[M+H]^+$ ion at *m*/*z* 337.19092, corresponding to the chemical formula $C_{21}H_{24}N_2O_2$ (δ ppm = -0.399). (**b**) $MS^2$ fragmentation spectrum of the $[M+H]^+$ ion at *m*/*z* 337.19. A single major daughter ion was observed at *m*/*z* 320 (loss of $NH_3$). A minor ion was observed at *m*/*z* 308. (**c**) $MS^3$ fragmentation spectrum of the daughter ion at *m*/*z* 320. (**d**) $MS^3$ fragmentation spectrum of the daughter ion at *m*/*z* 308.

The unknown metabolite has a calculated chemical formula that is identical to catharanthine. The observed loss of ammonia ($NH_3$) was also observed for strictosidine and may point to a similar A-B-C-ring structure in the unknown catharanthine isomer.

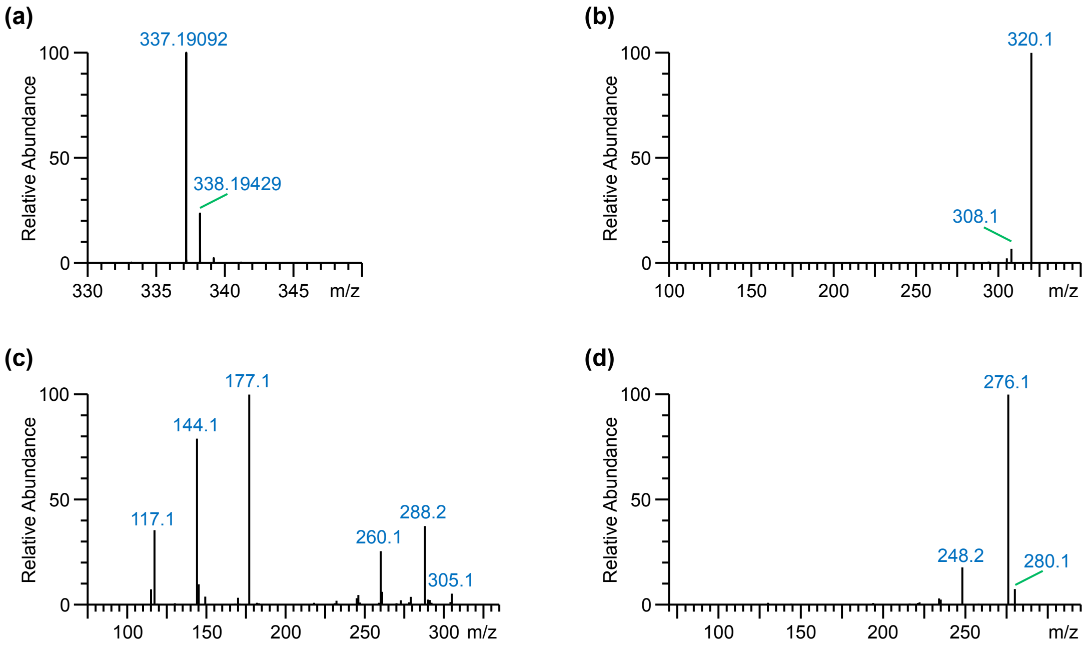


**Figure 28.** Identification of an unknown MIA (catharanthine isomer 2). (**a**) FT-MS scan of the unknown peak revealed an accurate mass of the $[M+H]^+$ ion at *m*/*z* 337.19092, corresponding to the chemical formula $C_{21}H_{24}N_2O_2$ ($\delta$ ppm = -0.399). (**b**) $MS^2$ fragmentation spectrum of the $[M+H]^+$ ion at *m*/*z* 337.19. A single major daughter ion was observed at *m*/*z* 320 (loss of $NH_3$). A minor ion was observed at *m*/*z* 308. (**c**) $MS^3$ fragmentation spectrum of the daughter ion at *m*/*z* 320. (**d**) $MS^3$ fragmentation spectrum of the daughter ion at *m*/*z* 308.

The calculated chemical formula and fragmentation spectrum of this unknown metabolite is nearly identical to that of catharanthine isomer 1.

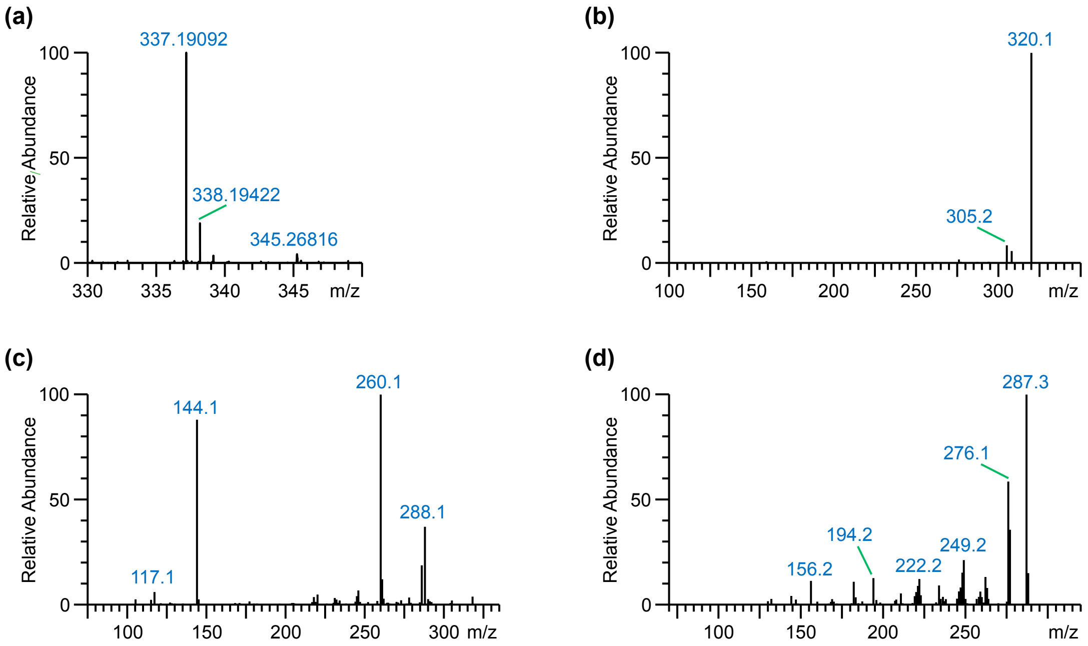


**Figure 29.** Identification of an unknown MIA (catharanthine isomer 3). (**a**) FT-MS scan of the unknown peak revealed an accurate mass of the $[M+H]^+$ ion at *m*/*z* 337.19092, corresponding to the chemical formula $C_{21}H_{24}N_2O_2$ (δ ppm = -0.399). (**b**) $MS^2$ fragmentation spectrum of the $[M+H]^+$ ion at *m*/*z* 337.19. A single major daughter ion was observed at *m*/*z* 320 (loss of $NH_3$). A minor ion was observed at *m*/*z* 305. (**c**) $MS^3$ fragmentation spectrum of the daughter ion at *m*/*z* 320. (**d**) $MS^3$ fragmentation spectrum of the daughter ion at *m*/*z* 305.

The calculated chemical formula and fragmentation spectrum of this unknown metabolite is similar to that of the unknown metabolites that were annotated as catharanthine isomers 1 and 2.

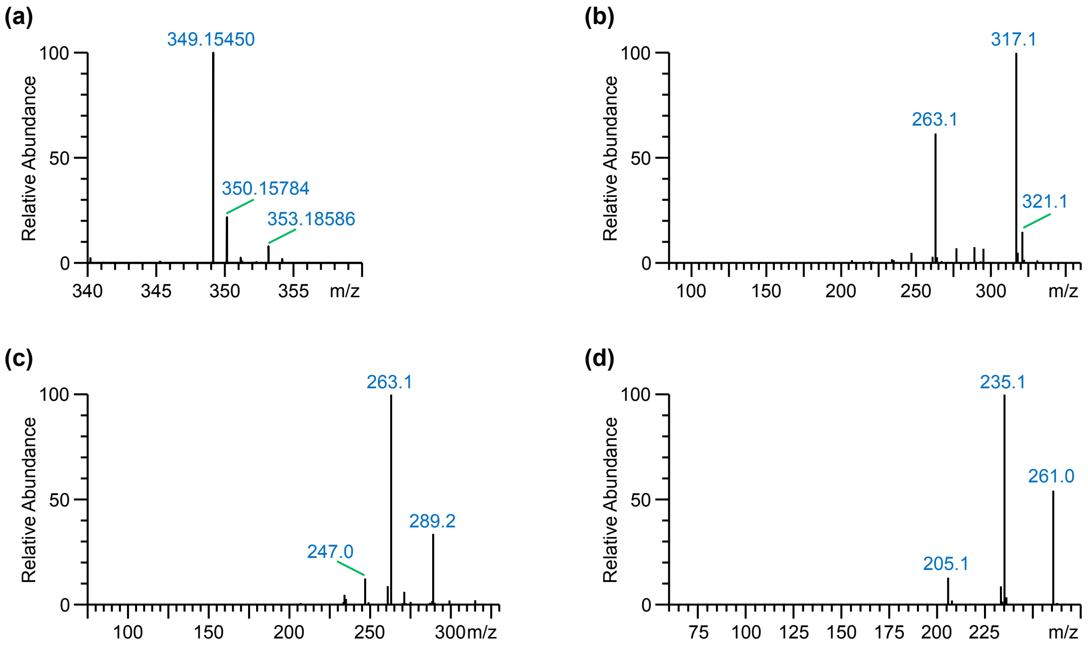


**Figure 30.** Identification of an unknown MIA (serpentine isomer). (**a**) FT-MS scan of the unknown peak revealed an accurate mass of the $[M+H]^+$ ion at *m*/*z* 349.15450, corresponding to the chemical formula of serpentine, $C_{21}H_{20}N_2O_3$ (δ ppm = -0.484). (**b**) $MS^2$ fragmentation spectrum of the $[M+H]^+$ ion at *m*/*z* 349.15. Major daughter ions are observed at *m*/*z* 317 (loss of MeOH) and *m*/*z* 263. (**c**) $MS^3$ fragmentation spectrum of the daughter ion at *m*/*z* 317. (**d**) $MS^3$ fragmentation spectrum of the daughter ion at *m*/*z* 263.

The observed chemical formula and fragmentation spectrum is nearly identical to that of serpentine (Schweizer et al., 2018), hence, this unknown metabolite was annotated as a serpentine isomer.
