## Supporting Information for "Subfunctionalization of paralog transcription factors contributes to regulation of alkaloid pathway branch choice in *Catharanthus roseus*"

***The plant journal* Supporting Information**

Article title: **Subfunctionalization of paralog transcription factors contributes to regulation of terpenoid indole alkaloid branch choice in *Catharanthus roseus***

Authors: Maite Colinas, Jacob Pollier, Dries Vaneechoutte, Deniz G. Malat, Fabian Schweizer, Rebecca De Clercq, Joana G. Guedes, Teresa Martínez-Cortés, Francisco J. Molina Hidalgo, Mariana Sottomayor, Klaas Vandepoele, and Alain Goossens

The following supporting information is available:

**Fig. S1** MIA gene expression levels upon bulk overexpression of TF candidates.

**Fig. S2** Separate overexpression of positive TF candidates.

**Fig. S3** Promoter transactivation of iridoid pathway genes by combinations of BIS.

**Fig. S4** Comparison of structural features and motifs of ORCA activation domains (ADs).

**Fig. S5** Expression profiles of MIA biosynthesis genes and regulators in analyzed RNA-seq data.

**Fig. S6** MIA levels upon overexpression of *ORCAs*.

**Fig. S7** MIA gene expression levels upon overexpression of *ORCA3* single mutants.

**Fig. S8** MIA levels upon overexpression of *ORCA3* mutants.

**Fig. S9** MIA levels upon combinatorial overexpression of *CrMYC2a^D126N^* and *ORCAs*.

**Fig. S10** Orthologs of cuticular wax regulators up-regulate the vindoline pathway gene *T3R*.

**Fig. S11** Combinatorial overexpression of *ORCA4, CrMYB96, CrMYC2a^D126N^*.

**Table S1** List of RNA-Seq datasets used.

**Table S2** Oligonucleotide primers used for cloning of coding sequences.

**Table S3** Oligonucleotide primers used for cloning of promoter fragments.

**Table S4** List of all cloned TFs and promoter fragments and Genebank IDs.

**Table S5** Oligonucleotide primers used for qPCR.

**Table S6** MIA levels upon overexpression of *ORCAs*.

**Table S7** MIA levels upon overexpression of *ORCA3* mutants.

**Table S8** MIA levels upon combinatorial overexpression of *CrMYC2a^D126N^* and *ORCAs.*

**Table S9** MIA levels upon overexpression of *MYB96/b*.

**Table S10** MIA levels upon overexpression of *MYB96/b, CrMYC2a^D126N^* and *ORCA4*

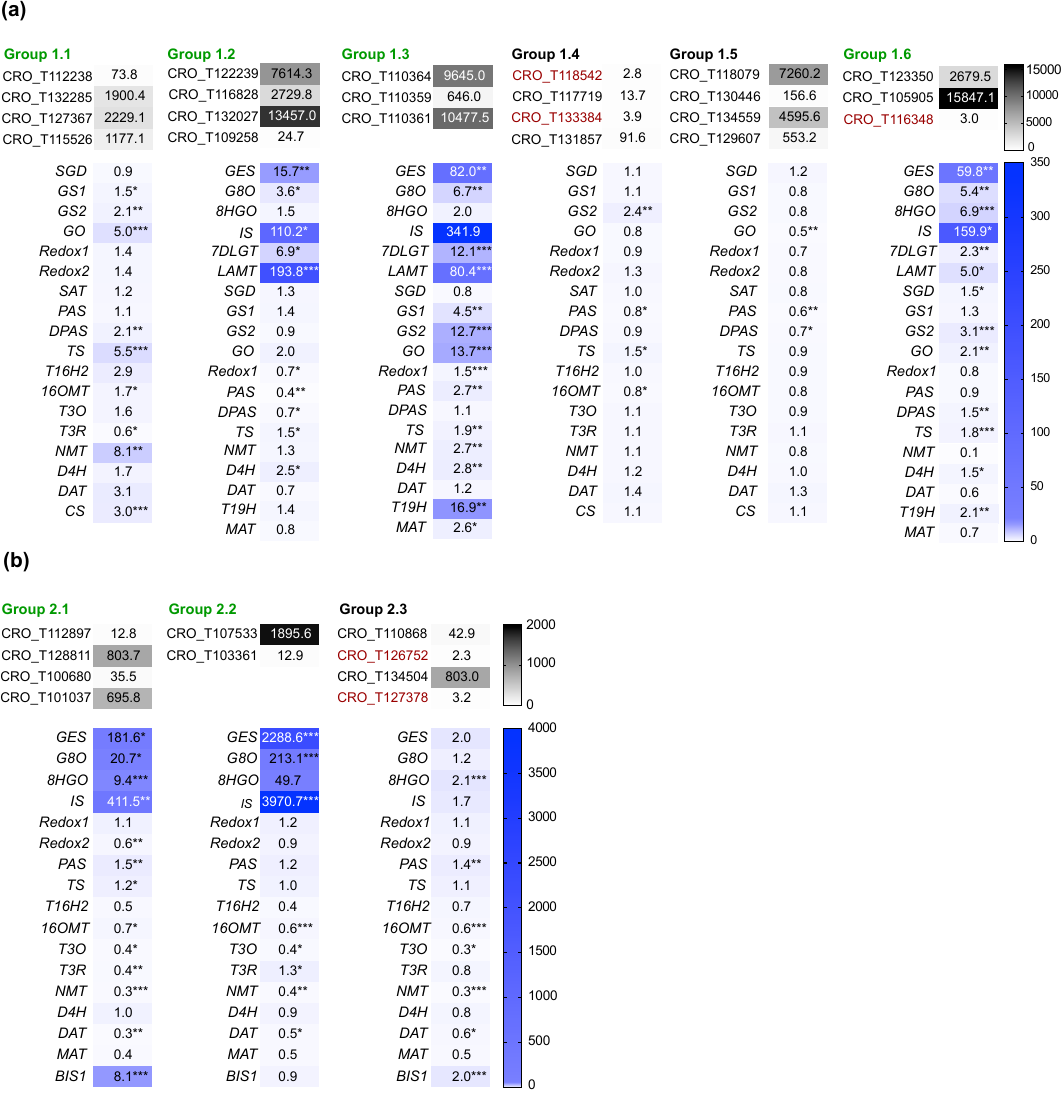

**Fig. S1 MIA gene expression levels upon bulk overexpression of TF candidates.** Up to four TF candidates were bulk overexpressed via flower petal infiltration with *A. tumefaciens* in two rounds (a and b), and the expression level of MIA genes subsequently measured by qPCR. Gene expression levels are expressed as fold changes relative to that of the *p35S::GUS* control. *P<0.05, **P<0.01, ***P<0.001 calculated by two-tailed Student’s *t*-test of four biological replicates. Groups marked in green were chosen for subsequent separate infiltration of TF candidates. Gene IDs labeled in red were only mildly overexpressed (<5-fold) and not taken along for further analyses.

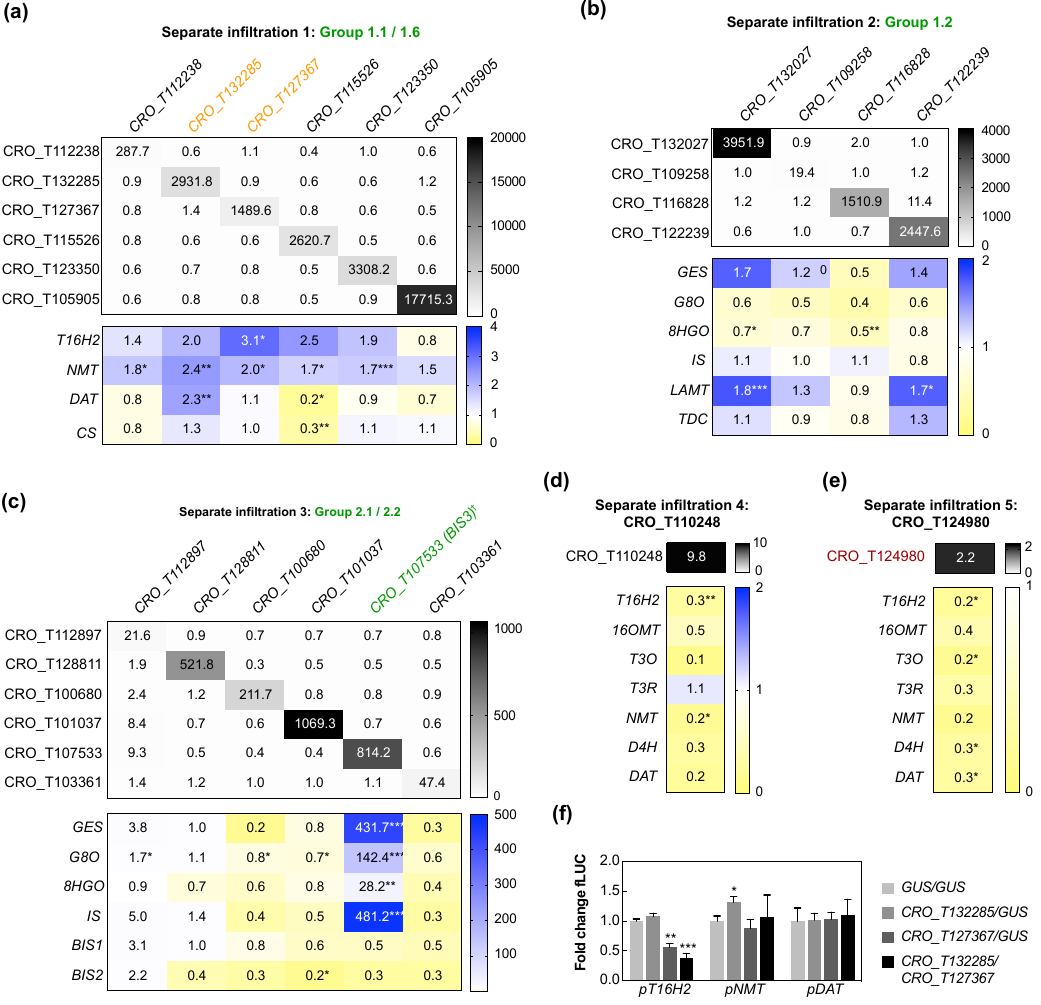

**Fig. S2 Separate overexpression of positive TF candidates.** TF candidates selected from results from bulk overexpression (Fig. S1) were infiltrated separately in independent infiltration series. Gene expression levels measured by qPCR are expressed as fold changes relative to the expression in the *p35S::GUS* control. *P<0.05, **P<0.01, ***P<0.001 calculated by two-tailed Student’s *t*-test, four biological replicates. Gene IDs labeled in red were only mildly overexpressed (<5-fold).

**(a)** Genes from screening group 1.1 and 1.6 that were well overexpressed (see Fig. S1) were overexpressed separately. Overexpression of CRO_T132285 and CRO_T127367 (orange colored) led to a statistically significant >2-fold up-regulation of some vindoline genes. CRO_T123350 and CRO_T105905 can be classified as bHLH Clade IVa TFs due to sequence homology, and were therefore additionally tested for up-regulation of iridoid pathway genes (Fig. 2).

**(b)** Separate overexpression of genes from screening group 1.2. The previously observed up-regulation of early MIA pathway genes was not confirmed.

**(c)** Separate overexpression of TF candidates from bulk overexpression groups 2.1 and 2.2. † CRO_T107533 (re-named *BIS3*) was identified as a *BIS* paralog (see Fig. 3).

**(d)** and **(e)** These TF candidates had been previously selected as potential MIA pathway regulators by independent co-expression analyses and overexpressed separately.

**(f)** In order to clarify their up-regulation effect on MIA genes upon overexpression in flower petals (see a), candidates CRO_T132285 and CRO_T127367 were tested in promoter transactivation assays in *N. tabacum* protoplasts. The absence of a consistent activation of these promoter fragments does not support a direct involvement in vindoline pathway regulation. *P<0.05, **P<0.01, ***P<0.001 was calculated by two-tailed Student’s *t*-test. Error bars depict SEM of four biological replicates.

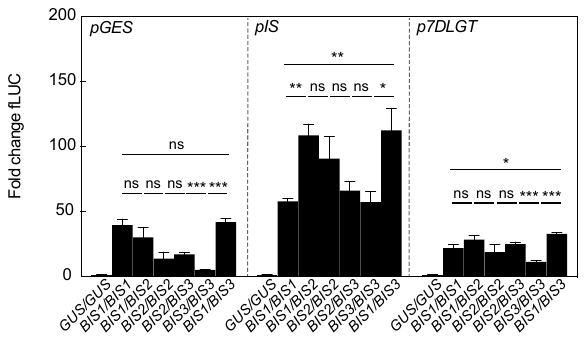

**Fig. S3 Promoter transactivation of iridoid pathway genes by combinations of BIS’.**

It was previously shown that BIS1 and BIS2 physically interact and to a certain extent synergistically up-regulate iridoid pathway target gene (Van Moerkercke et al. 2016). Here, transactivation assays of selected iridoid pathway promoter fragments (*pGES*, *pIS* and *p7DLGT*) were tested combinations of BIS’. An increase in transactivation by different BIS TFs was limited to transactivation of *pIS* and *p7DLGT* by BIS1 and 3 only. *N. tabacum* protoplasts were co-transfected with constructs containing the *fLUC* gene to be expressed under control of the indicated promoter fragments and constructs for overexpression of *BIS’* or *GUS* as a control. For individual overexpression of BIS1, 2 or 3 protoplasts were transfected with the double amounts of BIS1, 2 or 3 expression plasmids in order to compare effects of potential homo-dimers versus hetero-dimers. The y-axis shows fold change in normalized fLUC activity relative to the control transfection with GUS, set at 1. The error bars depict SEM of four biological replicates. Asterisks indicate statistically significant differences in transactivation (*P < 0.05, **P < 0.01, ***P < 0.01) as calculated by Student’s *t*-test.

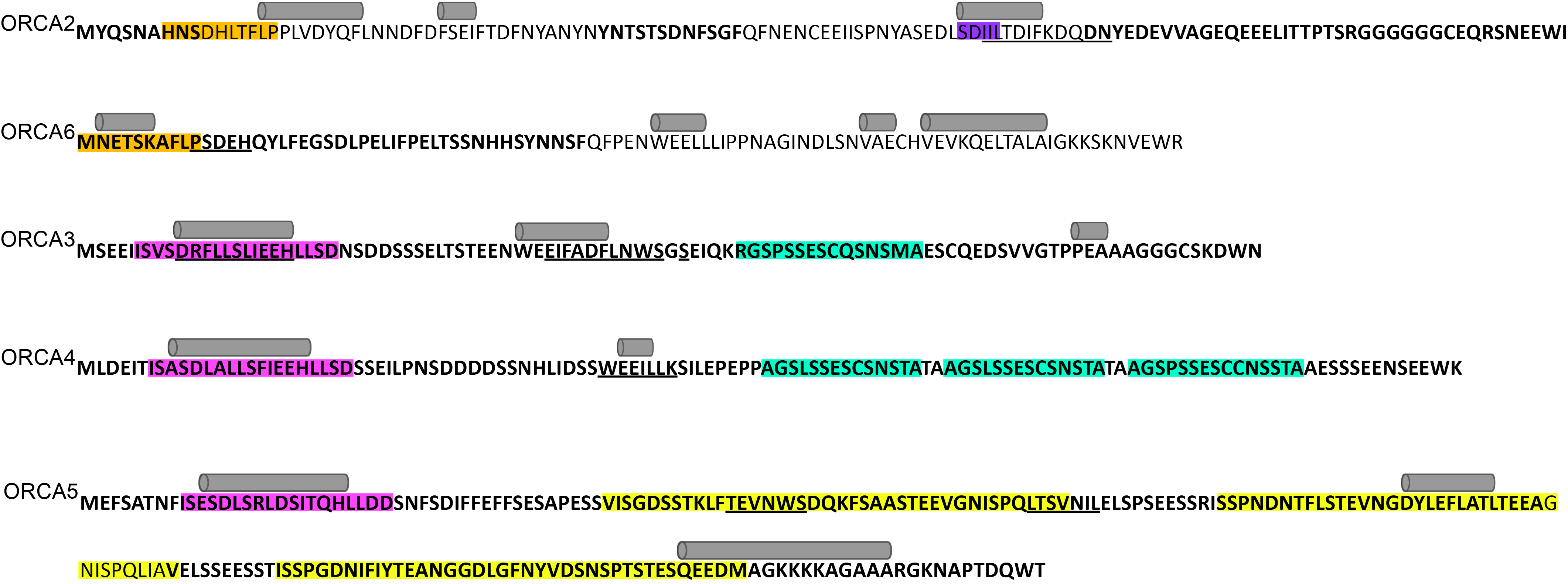

**Fig. S4 Comparison of structural features and motifs of ORCA activation domains (ADs).** While the DNA binding domain (DBD) is assumed to determine binding activity to specific DNA motifs, the AD is thought to confer the level of activation for instance by recruiting the transcriptional machinery and/or by interaction with other TFs or co-regulators. The observed differences in MIA target gene activation are only partly explained by differences in specific amino acid residues (AARs) in the DBD. In particular, a general lower activity (such as observed for the ORCA6 paralog) or the observed differences in synergistic up-regulation (i.e. higher for ORCA3 than for the other ORCAs) might be also caused by differences in the ADs. ADs commonly show a high level of disorder so that structural information by protein crystallization is scarce, and a very low level of AAR sequence conservation compared to the DBD, however structural features and motifs can be revealed using a combination of *in silico* methods (O'Shea *et al.*, 2017). To compare ORCA AD sequences despite their low homology, we used different prediction and motif search programs. As expected, PONDR® VSL2 predicts long stretches of the ORCA ADs to be disordered (bold font); in case of ORCA3-5 practically the entire AD is predicted to be disordered (Xue *et al.*, 2010). Within disordered regions, molecular recognition features (MoRFs) can occur. These short regions can undergo disorder-to-order transitions and bind protein partners, and can be predicted by MoRFpred (underlined sequence) (Disfani *et al.*, 2012). The 𝛼-helices (gray cylinders above the sequence) predicted by PSIPRED 4.0 often coincide with MoRFs, further indicating that these short regions may indeed form secondary structures (Buchan and Jones, 2019). Moreover, we performed a MEME analysis to uncover motifs with sequence similarity (depicted by colors) (Bailey *et al.*, 2015). In summary, the ADs of ORCA3, 4 and 5 share for instance in particular a motif at the N-terminus (pink), but they do not share any of the predicted motifs or structural features with those of ORCA2 and 6. The observed differences in MIA gene up-regulation between the ORCA3 and 4 domain swaps (Fig. 7) may be due to specific motifs but this requires further future experimentation.

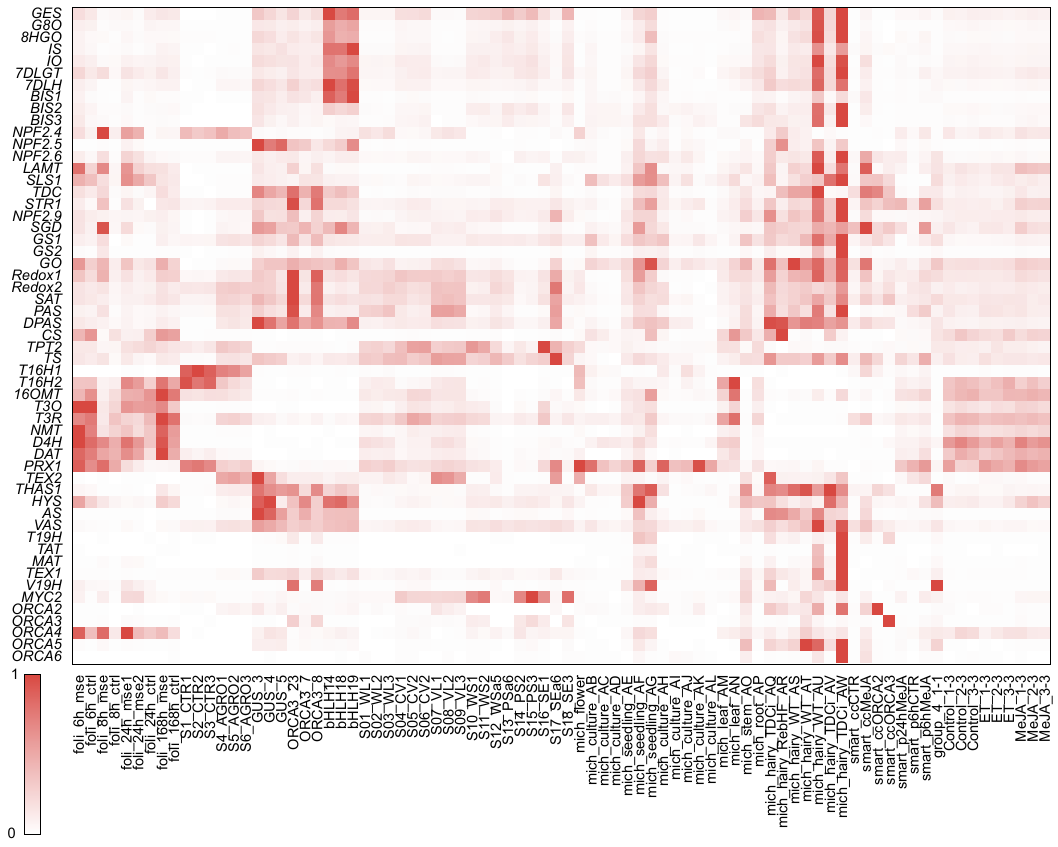

**Fig. S5 Expression profiles of MIA biosynthesis genes and regulators in analyzed RNA-seq data.**

Expression levels of MIA pathway genes and BIS and ORCA transcription factors are shown for all 81 RNA-seq samples included in the candidate selection analysis. TPM values were extracted from the expression atlas (Supporting dataset 1) and expression levels scaled from 0 (lowest TPM value) to 1 (highest TPM value) for each gene across all RNA-seq samples.

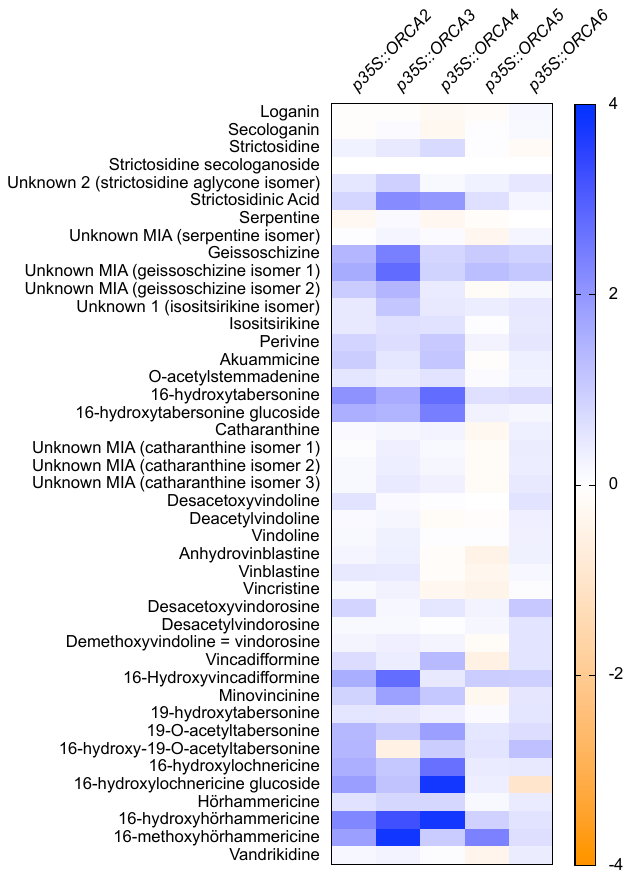

**Fig. S6 MIA levels upon overexpression of *ORCAs*.** Levels are expressed as log2 fold changes (FC) compared to the *p35S::GUS* control. Total ion current (TIC) values from this experiment, including statistical analysis, can be found in Table S6.

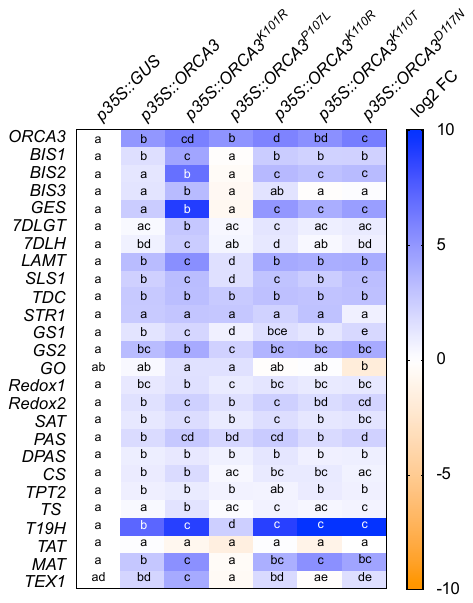

**Fig. S7 MIA gene expression levels upon overexpression of *ORCA3* single mutants.** The indicated AARs of ORCA3 were mutated to correspond to those of other ORCA TFs (see Fig. 5a). Gene expression levels measured by qPCR are expressed as average log2 FC relative to the *p35S::GUS* control. For each gene, different letters represent statistically significant differences (P<0.05, ANOVA with Tukey’s correction for multiple comparisons).

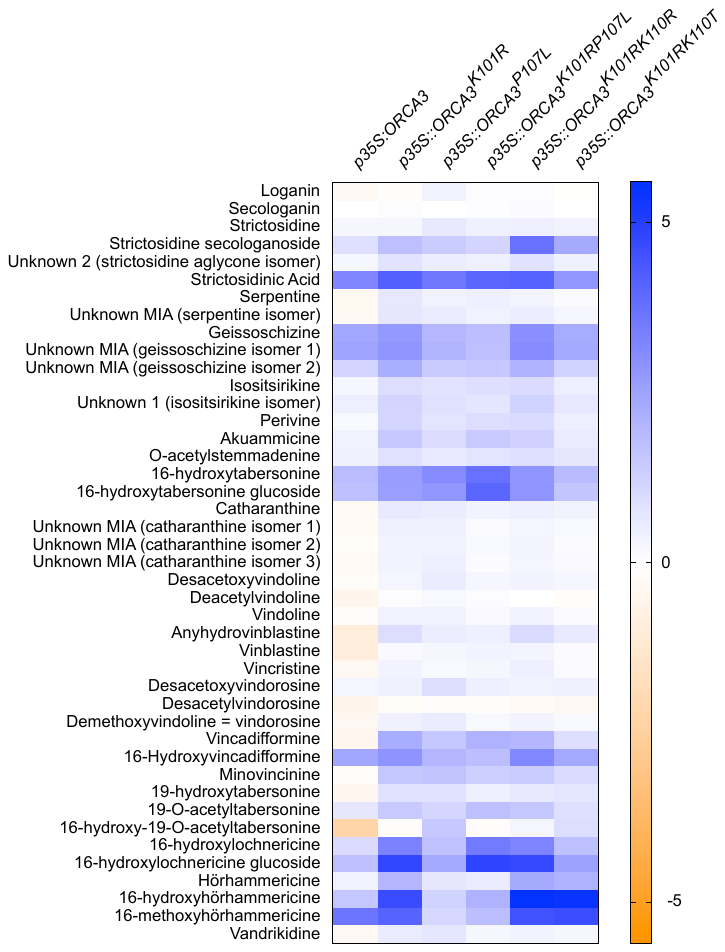

**Fig. S8 MIA levels upon overexpression of *ORCA3* mutants.** Levels are expressed as log2 FC compared to the *p35S::GUS* control. Total ion current (TIC) values from this experiment including statistical analysis can be found in Table S7.

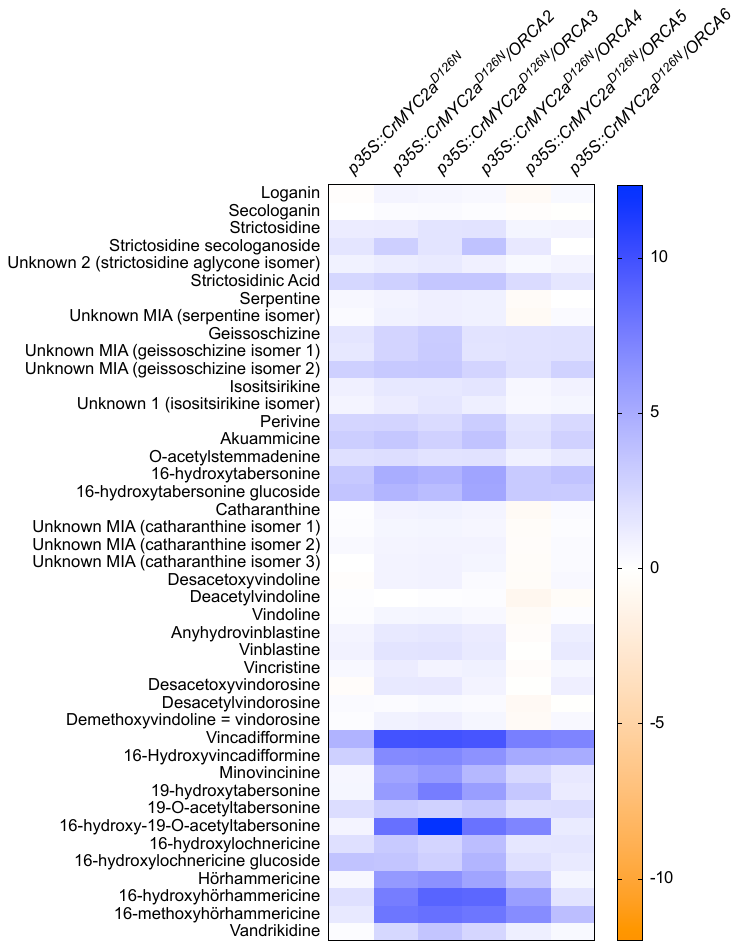

**Fig. S9 MIA levels upon combinatorial overexpression of *CrMYC2a^D126N^* and *ORCAs*.** Levels are expressed as log2 FC compared to the *p35S::GUS* control. Total ion current (TIC) values from this experiment including statistical analysis can be found in Table S8.

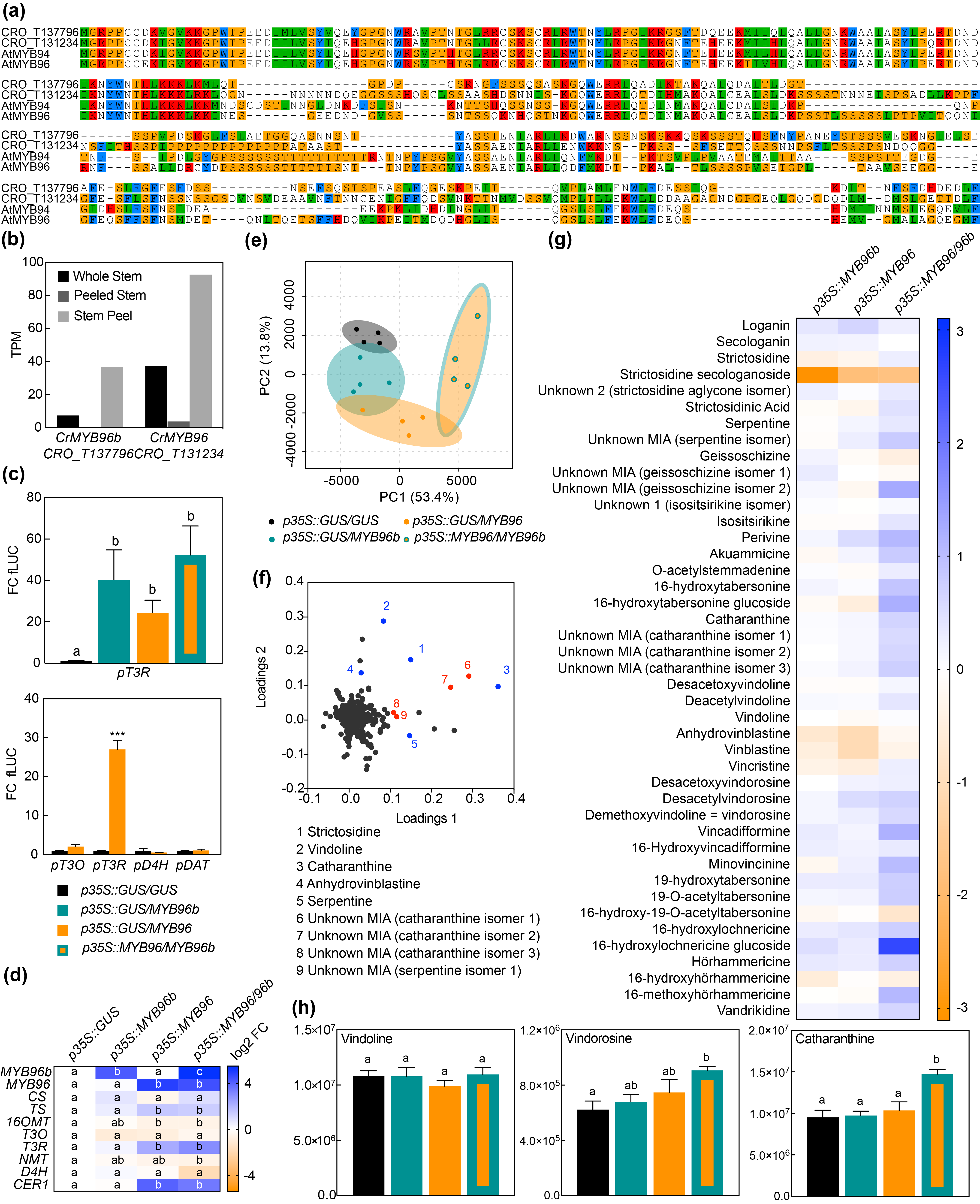

**Fig. S10 Orthologs of cuticular wax biosynthesis regulators up-regulate the vindoline pathway gene *T3R*.** The *A. thaliana* TFs MYB94 and 96 have been implicated in up-regulation of cuticular wax biosynthesis. When screening epidermis-enriched transcriptome data for potential epidermis-specific MIA biosynthesis regulators, the *C. roseus* orthologs CRO_T137796 and CRO_T131234 appeared and were renamed CrMYB96b and CrMYB96, respectively.

**(a**) MUSCLE protein sequence alignment of *A. thaliana* MYB94 and 96 and *C. roseus* orthologs.

**(b)** Transcripts Per Kilobase Million (TPM) values for *CrMYB96* and *96b* extracted from transcriptome data of macro-dissected stems of *C. roseus* (see Methods), showing that these transcripts are more abundant in the epidermis-enriched peel.

**(c)** Promoter transactivation assays in *N. tabacum* protoplasts by measuring luciferase activity upon co-transfection with constructs containing promoter fragments fused to *fLUC* and constructs for overexpression of Cr*MYB96/b* or *GUS* as a control. The y-axis shows fold change in normalized fLUC activity relative to the GUS control. The error bars depict SEM of four biological replicates. Upper panel: columns labelled with different letters represent statistically significant differences (P<0.05, ANOVA with Tukey’s correction for multiple comparisons). Lower panel: MYB96 was tested for transactivation of other vindoline pathway promoters; for every promoter statistical significance was determined by the Student's *t*-test (*P < 0.05, **P < 0.01, ***P < 0.001) compared to GUS control.

**(d) – (h)** Transient overexpression of *CrMYB96* and *96b* in *C. roseus* flower petals.

**(d)** Overexpression level and MIA biosynthesis gene expression levels were measured by qPCR and are expressed as average log2 FC relative to the *p35S::GUS* control. For each gene, different letters represent statistically significant differences (P<0.05, ANOVA with Tukey’s correction for multiple comparisons). In this experiment, overexpression of *CrMYB96* (or combination with *CrMYB96b*) leads to higher up-regulation of *T3R* than overexpression of *CrMYB96b* alone; therefore CrMYB96 was used in further experiments. Moreover, an ortholog of the cuticular wax gene *CER1* was also up-regulated, suggesting that the role in cuticular wax biosynthesis regulation is evolutionary conserved.

**(e)** PCA plot of metabolite profile data.

**(f)** Loading plot of PCA showing selected MIAs identified in previous studies (blue) and in this study (red).

**(g)** MIA levels are expressed as log2 FC compared to the *p35S::GUS* control. Total ion current (TIC) values from this experiment including statistical analysis can be found in Table S9.

**(h)** TIC values for vindoline, vindorosine and catharanthine. Columns labelled with different letters represent statistically significant differences (P<0.05, ANOVA with Tukey’s correction for multiple comparisons).

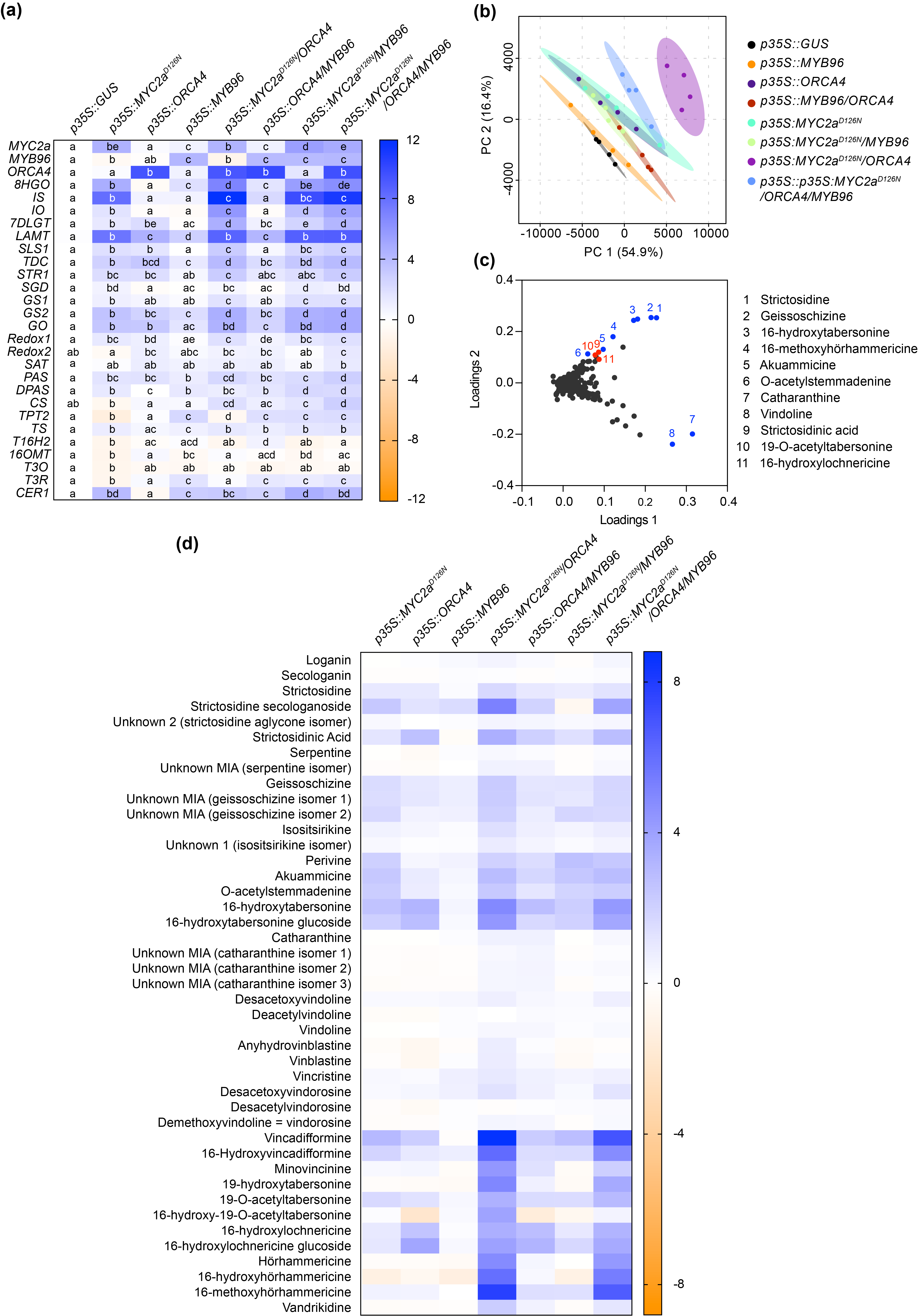

**Fig. S11 Combinatorial overexpression of *ORCA4*, *CrMYB96* and *CrMYC2a^D126N^*.**

**(a)** Gene expression levels were measured by qPCR and are expressed as average log2 FC relative to the *p35S::GUS* control. For each gene, different letters represent statistically significant differences (P<0.05, ANOVA with Tukey’s correction for multiple comparisons).

**(b)** PCA metabolite profile data.

**(c)** Loading plot of PCA showing selected MIAs previously identified (blue) and in this study (red).

**(d)** MIA levels are expressed as log2 FC compared to the *p35S::GUS* control. Consistent with other experiments, the MIA profile significantly changes upon overexpression of *ORCA4* or *CrMYC2a^D126N^* alone or in combination, as opposed to overexpression of *MYB96*. Total ion current (TIC) values from this experiment including statistical analysis can be found in Table S10.

**Table S1** List of RNA-Seq datasets used.

| **Dataset ID**  **(Reference)** | **Sample ID** | **Sample name (as in Supporting Dataset 1)** | **Short sample description** |
| --- | --- | --- | --- |
| SRA030483  (Gongora-Castillo *et al.*, 2012) | SRR122239 | mich_flower | flowers |
|  | SRR122240 | mich_culture_AB | Suspension culture (yeast extract (YE) 0.3 mg/mL for 12 hours) |
|  | SRR122241 | mich_culture_AC | Suspension culture (YE 0.3 mg/mL for 24 hours) |
|  | SRR122242 | mich_culture_AD | Suspension culture control |
|  | SRR122243 | mich_seedling_AE | Sterile seedlings |
|  | SRR122244 | mich_seedling_AF | Sterile seedlings (Methyl jasmonate(MeJA) 6uM 12 days) |
|  | SRR122245 | mich_seedling_AG | Sterile seedlings |
|  | SRR122246 | mich_culture_AH | Suspension culture (YE 0.3 mg/mL for 6 hours) |
|  | SRR122247 | mich_culture_AI | Suspension culture (MeJA 100uM for 6 hours) |
|  | SRR122248 | mich_culture_AJ | Suspension culture (MeJA 100uM for 12 hours) |
|  | SRR122249 | mich_culture_AK | Suspension culture (MeJA 100uM for 24 hours) |
|  | SRR122250 | mich_culture_AL | Suspension culture |
|  | SRR122251 | mich_leaf_AM | Mature leaves |
|  | SRR122252 | mich_leaf_AN | Immature leaves |
|  | SRR122253 | mich_stem_AO | Stem |
|  | SRR122254 | mich_root_AP | Root |
|  | SRR122255 | mich_hairy_TDCi_AQ | Hairy roots (TDCi) |
|  | SRR122256 | mich_hairy_RebHF_AR | Hairy roots (RebH_F) |
|  | SRR122257 | mich_hairy_WT_AS | Wild type (wt) hairy roots |
|  | SRR122258 | mich_hairy_WT_AT | Wt hairy roots (MeJA 250uM for o hours) |
|  | SRR122259 | mich_hairy_WT_AU | Wt hairy roots (MeJA 250uM for 24 hours) |
|  | SRR122260 | mich_hairy_TDCi_AV | TDCi hairy roots (MeJA 250uM for 0 hours) |
|  | SRR122261 | mich_hairy_TDCi_AW | TDCi hairy roots (MeJA 250uM for 24 hours) |
| SRA064724  (Van Moerkercke *et al.*, 2013) | SRS385814 | smart_ccCTR | Cell suspension culture (CC) control |
|  | SRS385815 | smart_ccMeJA | CC exposed to MeJA |
| SRP026417  (Van Moerkercke *et al.*, 2015) | SRR924147 | smart_ccORCA2 | CC overexpressing *ORCA2* |
|  | SRR924148 | smart_ccORCA3 | CC overexpressing *ORCA3* |
| SRA064076  (Van Moerkercke *et al.*, 2013) | SRS383709 | smart_p24hMeJA | Plants 24 hours MeJA |
|  | SRS383671 | smart_p6hCTR | Control plants 6 hours |
|  | SRS383701 | smart_p6hMeJA | Plants 6 hours MeJA |
| ERA669805  (Dugé de Bernonville *et al.*, 2017) | ERR1512369 | foli_6h_mse | Plants exposed to *Manduca sexta* (*Ms*) 6 hours |
|  | ERR1512370 | foli_6h_ctrl | *Ms* control plants 6 hours |
|  | ERR1512371 | foli_8h_mse | *Ms* plants 8 hours |
|  | ERR1512372 | foli_8h_ctrl | *Ms* control plants 8 hours |
|  | ERR1512373 | foli_24h_mse1 | *Ms* plants 24 hours |
|  | ERR1512374 | foli_24h_mse2 | *Ms* plants 24 hours |
|  | ERR1512375 | foli_24h_ctrl | *Ms* control plants 24 hours |
|  | ERR1512376 | foli_168h_mse | *Ms* plants 168 hours, new leaves |
|  | ERR1512377 | foli_168h_ctrl | *Ms* control plants 168 hours, new leaves |
| SRP095740  (Pan *et al.*, 2018) | SRR5133632 | ET_1-3 | Seedlings exposed to ethylene (ET) |
|  | SRR5133633 | ET_2-3 |  |
|  | SRR5133635 | ET_3-3 |  |
|  | SRR5133629 | MeJA_2-3 | Seedlings exposed to MeJA |
|  | SRR5133628 | MeJA_1-3 |  |
|  | SRR5133636 | MeJA_3-3 |  |
|  | SRR5133634 | Control_3-3 | Control seedlings |
|  | SRR5133631 | Control_2-3 |  |
|  | SRR5133630 | Control_1-3 |  |
|  | SRR5944780 | group_4_1-1 | Seedlings |
| SRP055543  (Van Moerkercke *et al.*, 2015) | SRS858191 | GUS_3 | Control hairy root lines overexpressing *GUS* |
|  | SRS858189 | GUS_4 |  |
|  | SRS858216 | GUS_5 |  |
| SRP055543  (Van Moerkercke *et al.*, 2015) | SRS858143 | bHLH_14 | Hairy root lines overexpressing *BIS1* |
|  | SRS858141 | bHLH_18 |  |
|  | SRS858111 | bHLH_19 |  |
| PRJEB40213 (This study) | ERR4567494 | ORCA3_23 | Hairy root lines overexpressing *ORCA3* |
|  | ERR4567495 | ORCA3_7 |  |
|  | ERR4567496 | ORCA3_8 |  |
| PRJEB40216 (This study) | ERR4567574 | S01_WL1 | Whole leaves |
|  | ERR4567575 | S02_WL2 |  |
|  | ERR4567576 | S03_WL3 |  |
|  | ERR4567577 | S04_CV1 | Central vein |
|  | ERR4567578 | S05_CV2 |  |
|  | ERR4567579 | S06_CV2 |  |
|  | ERR4567580 | S07_VL1 | Leaf without central vein |
|  | ERR4567581 | S08_VL2 |  |
|  | ERR4567582 | S09_VL3 |  |
|  | ERR4567583 | S10_WS1 | Whole stem |
|  | ERR4567584 | S11_WS2 |  |
|  | ERR4567585 | S12_WSa5 |  |
|  | ERR4567586 | S13_PSa6 | Peeled stem |
|  | ERR4567587 | S14_PS2 |  |
|  | ERR4567588 | S15_PS3 |  |
|  | ERR4567589 | S16_SE1 | Stem epidermis |
|  | ERR4567590 | S17_SEa6 |  |
|  | ERR4567591 | S18_SE3 |  |
| PRJEB40214  (This study) | ERR4567511 | S1_CTR1 | Mock infiltrated flower petals |
|  | ERR4567512 | S2_CTR2 |  |
|  | ERR4567513 | S3_CTR3 |  |
|  | ERR4567514 | S4_AGRO1 | *A. tumefaciens* C58C1 infiltrated flower petals |
|  | ERR4567515 | S5_AGRO2 |  |
|  | ERR4567516 | S6_AGRO3 |  |

**Table S2** Oligonucleotide primers used for cloning of coding sequences.

| CRO_ID/  Gene or TF family | Sequence fw/  rev |
| --- | --- |
| CRO_T110359 | GGGGACAAGTTTGTACAAAAAAGCAGGCTTAATGGAGTTTTCAGCTACTAATTTTATC |
| ORCA5 | GGGGACCACTTTGTACAAGAAAGCTGGGTATTACAATATTGTCTCCTGTTTCATCT |
| CRO_T110361 | GGGGACAAGTTTGTACAAAAAAGCAGGCTTAATGCTTGATGAAATTACTATTTCAGCC |
| ORCA4 | GGGGACCACTTTGTACAAGAAAGCTGGGTATTAATTTCTTCTCTTCTTTCGTCCCT |
| CRO_T110364 | GGGGACAAGTTTGTACAAAAAAGCAGGCTTAATGAATGAAACTTCAAAAGCCTTCCT |
| ORCA6 | GGGGACCACTTTGTACAAGAAAGCTGGGTACTATTGTTTACGGCCAAGATGATT |
| CRO_T112238 | GGGGACAAGTTTGTACAAAAAAGCAGGCTTAATGGAGTGCTTCAAAAATCATCTTCC |
| GATA | GGGGACCACTTTGTACAAGAAAGCTGGGTACTAAGAATTAGATGGTGGGTGGGA |
| CRO_T132285 | GGGGACAAGTTTGTACAAAAAAGCAGGCTTAATGGAGATGGAGAAATCTTGTGA |
| BBX | GGGGACCACTTTGTACAAGAAAGCTGGGTATCACTGAACGTTGGACAAACA |
| CRO_T112897 | GGGGACAAGTTTGTACAAAAAAGCAGGCTTAATGTCTTCTTGTTTAACTGGAGGTG |
| CCT | GGGGACCACTTTGTACAAGAAAGCTGGGTACTATCCTTGCTCGCTCTCTGG |
| CRO_T127367 | GGGGACAAGTTTGTACAAAAAAGCAGGCTTAATGGTACAATCAAAGAAGTTCAGAGG |
| AP2/ERF | GGGGACCACTTTGTACAAGAAAGCTGGGTATCAATTTCTGTTCAAGAGTTCTTCA |
| CRO_T130446 | GGGGACAAGTTTGTACAAAAAAGCAGGCTTAATGGTATCTTCTGATAATAATAAAAAG |
| BBX | GGGGACCACTTTGTACAAGAAAGCTGGGTATCATTTATTTTGCAGGGGGAAAACA |
| CRO_T118079 | GGGGACAAGTTTGTACAAAAAAGCAGGCTTAATGTCTCACATAGCAGTGGAAAGA |
| bHLH | GGGGACCACTTTGTACAAGAAAGCTGGGTACTACGGTTCTATTTGTTTGGAGCA |
| CRO_T128811 | GGGGACAAGTTTGTACAAAAAAGCAGGCTTAATGGCTGCCAAGGTCAGG |
| C2H2 ZF | GGGGACCACTTTGTACAAGAAAGCTGGGTATTATCTCGACCGGCCTAATCTC |
| CRO_T107533 | GGGGACAAGTTTGTACAAAAAAGCAGGCTTAATGGATATGATGACGGATAATTCAGGA |
| bHLH | GGGGACCACTTTGTACAAGAAAGCTGGGTACTACCCAAGAAGTGACCGAAGA |
| CRO_T132027 | GGGGACAAGTTTGTACAAAAAAGCAGGCTTAATGTATTGTAATTTTTGGTCAAGTAGT |
| bHLH | GGGGACCACTTTGTACAAGAAAGCTGGGTATCATAGATTGAATATCAAGCCCATCA |
| CRO_T109258 | GGGGACAAGTTTGTACAAAAAAGCAGGCTTAATGGAAAGCTTAGATATTCATGAAGAA |
| bHLH | GGGGACCACTTTGTACAAGAAAGCTGGGTATTACATACTCATGGGACTGTCTGG |
| CRO_T118542 | GGGGACAAGTTTGTACAAAAAAGCAGGCTTAATGGCTGCCCAAGTTTTAGA |
| ARR | GGGGACCACTTTGTACAAGAAAGCTGGGTATTATGAGAATATCTGATTGTATCTTT |
| CRO_T103361 | GGGGACAAGTTTGTACAAAAAAGCAGGCTTAATGATGAAAATGAAGTTACTCAATCGT |
| TPR | GGGGACCACTTTGTACAAGAAAGCTGGGTACTATTGCAGATTATCTACAGTCAAGAG |
| CRO_T117719 | GGGGACAAGTTTGTACAAAAAAGCAGGCTTAATGGCTGCTGAATCATCAAGC |
| AP2/ERF | GGGGACCACTTTGTACAAGAAAGCTGGGTATCAGCTGACCCATTTTACTGTGT |
| CRO_T133384 | GGGGACAAGTTTGTACAAAAAAGCAGGCTTAATGTCACTGAGGAGAAATTTTCTTT |
| TLP | GGGGACCACTTTGTACAAGAAAGCTGGGTATCACTCGCAGGCCAGTTTT |
| CRO_T122239 | GGGGACAAGTTTGTACAAAAAAGCAGGCTTAATGCAAAACCCTTCTCATAAACAT |
| NYF-YA | GGGGACCACTTTGTACAAGAAAGCTGGGTATCATTTTGGTGCACCTCCCT |
| CRO_T116828 | GGGGACAAGTTTGTACAAAAAAGCAGGCTTAATGGGTAGTACTAGATTCATGAACA |
| WRKY | GGGGACCACTTTGTACAAGAAAGCTGGGTATCATCCGGTGGTGCCACA |
| CRO_T123350 | GGGGACAAGTTTGTACAAAAAAGCAGGCTTAATGGATCTTTTAGGAGCACATGAGT |
| bHLH | GGGGACCACTTTGTACAAGAAAGCTGGGTATTATTGCATGAGCTTTTGGATGGC |
| CRO_T105905 | GGGGACAAGTTTGTACAAAAAAGCAGGCTTAATGGAAGCTCCATCAACTTCTTG |
| bHLH | GGGGACCACTTTGTACAAGAAAGCTGGGTACTATCTGTTTTTCTTTAGTGCTGA |
| CRO_T115526 | GGGGACAAGTTTGTACAAAAAAGCAGGCTTAATGGAGAAGAGTAGTACTAGTACT |
| ZF | GGGGACCACTTTGTACAAGAAAGCTGGGTACTAGAGATGAAGGTCTAAACTCACA |
| CRO_T129607 | GGGGACAAGTTTGTACAAAAAAGCAGGCTTAATGGGGAAGAAAATGAAGCTCC |
| OFP | GGGGACCACTTTGTACAAGAAAGCTGGGTACTAATTTCTCAGACACGAAGAAGAAGA |
| CRO_T131857 | GGGGACAAGTTTGTACAAAAAAGCAGGCTTAATGTACGAAAACGGTGATTTTGA |
| AP2/ERF | GGGGACCACTTTGTACAAGAAAGCTGGGTATCACCATTCACTAAACAATCTCTGT |
| CRO_T134559 | GGGGACAAGTTTGTACAAAAAAGCAGGCTTAATGAAGAAAATGATGAGAAGTCAACT |
| Homeodomain | GGGGACCACTTTGTACAAGAAAGCTGGGTATTAGCTAATTTGACTACAAAGTTGATT |
| CRO_T116348 | GGGGACAAGTTTGTACAAAAAAGCAGGCTTAATGGATAATTGTGAAATTAGCGAAAGA |
| HF | GGGGACCACTTTGTACAAGAAAGCTGGGTATTATTTGCACGATTGAAAGAGAAAAA |
| CRO_T100680 | GGGGACAAGTTTGTACAAAAAAGCAGGCTTAATGAAAGAGAGACAACGGTGGC |
| MYB | GGGGACCACTTTGTACAAGAAAGCTGGGTACTAATTAGGCTCATCCATTCGAGG |
| CRO_T101037 | GGGGACAAGTTTGTACAAAAAAGCAGGCTTAATGGTCAAATTTTCCAAGTTT |
| TCP | GGGGACCACTTTGTACAAGAAAGCTGGGTACTAATTTTGAAAATGGATTTGATTTTG |
| CRO_T110868 | GGGGACAAGTTTGTACAAAAAAGCAGGCTTAATGGACTCTAAGAAGCTTGTTAATGG |
| AP2/B3 | GGGGACCACTTTGTACAAGAAAGCTGGGTACTACCTACAAGTGACACCCTTGC |
| CRO_T126752 | GGGGACAAGTTTGTACAAAAAAGCAGGCTTAATGATCTCAAAAGGGTCGTCGT |
| ANAC | GGGGACCACTTTGTACAAGAAAGCTGGGTATTACTGAGCACAAATTAAGGTTCCA |
| CRO_T134504 | GGGGACAAGTTTGTACAAAAAAGCAGGCTTAATGGGTAGAAAGTGCTCACATTG |
| MYB | GGGGACCACTTTGTACAAGAAAGCTGGGTATCAGATGACGCTGATAATTGGTC |
| CRO_T127378 | GGGGACAAGTTTGTACAAAAAAGCAGGCTTAATGGCTGCAGAGTTGCAATT |
| ANAC | GGGGACCACTTTGTACAAGAAAGCTGGGTATTAAAATGGCTTCTGCAAGAACA |
| CRO_T110248 | GGGGACAAGTTTGTACAAAAAAGCAGGCTCGATGTTATCGAGAGTTAACAGCATGG |
| bHLH | GGGGACCACTTTGTACAAGAAAGCTGGGTTTAAACCATCCCTTGGAAGCC |
| CRO_T124980 | GGGGACAAGTTTGTACAAAAAAGCAGGCTCGATGGCTTTAGAAGCCCTTTCTTCC |
| bHLH | GGGGACCACTTTGTACAAGAAAGCTGGGTTCAACAAGTACGGGGTGGC |
| CRO_T110360 | GGGGACAAGTTTGTACAAAAAAGCAGGCTTAATGTCCGAAGAAATCATTTCCGT |
| ORCA3 | GGGGACCACTTTGTACAAGAAAGCTGGGTATTAATATCGTCTCTTCTTCCTTCCTCC |
| CRO_T110360 | GATTGGAACCGGTATAGAGGCGTTAGACGGCGG |
| ORCA3^K101R^ | CCGCCGTCTAACGCCTCTATACCGGTTCCAATC |
| CRO_T110360 | GTTAGACGGCGGCTTTGGGGGAAGTTCG |
| ORCA3^P107L^ | CGAACTTCCCCCAAAGCCGCCGTCTAAC |
| CRO_T110360 | GCCGTGGGGGAGATTCGCGGCGGAG |
| ORCA3^K110R^ | CTCCGCCGCGAATCTCCCCCACGGC |
| CRO_T110360 | GCCGTGGGGGACTTTCGCGGCGGAG |
| ORCA3^K110T^ | CTCCGCCGCGAAAGTCCCCCACGGC |
| CRO_T110360 | CGGCGGAGATAAGGAATCCGAAAAAGAAAG |
| ORCA3^D117N^ | CTTTCTTTTTCGGATTCCTTATCTCCGCCG |
| CRO_T110360/1 | GAGGAGGTTGTTCGAAGGATTGGAACCGGTACAGAGGCGTTAGACGG |
| ORCA3/4 swap | CCGTCTAACGCCTCTGTACCGGTTCCAATCCTTCGAACAACCTCCTC |
| CRO_T110360/1 | CGCCGTCTAACGCCCTTATACCGCTTCCATTCCTCAGAATTTTCTTCACTAC |
| ORCA4/3 swap | GTAGTGAAGAAAATTCTGAGGAATGGAAGCGGTATAAGGGCGTTAGACGGCG |
| CRO_T131234 | GGGGACAAGTTTGTACAAAAAAGCAGGCTCCATGGGAAGACCACCTTGCTGTGATA |
| MYB96 | GGGGACCACTTTGTACAAGAAAGCTGGGTGTCMGAACAAATCAGTAGTTTCCCCT |
| CRO_T137796 | GGGGACAAGTTTGTACAAAAAAGCAGGCTTAATGGGGAGACCTCCTTGCT |
| MYB96b | GGGGACCACTTTGTACAAGAAAGCTGGGTACTAAAACAAATCTTCATCATGATCAAA |
| CRO_T110365 | GGGGACAAGTTTGTACAAAAAAGCAGGCTccATGTATCAATCAAATGCCCATAATTCCG |
| ORCA2 | GGGGACCACTTTGTACAAGAAAGCTGGGTgTCMTTGAGGACGAAGATGACACGATGAA |

**Table S3** Oligonucleotide primers used for cloning of promoter fragments.

| CRO_ID/  Gene or TF family | Sequence fw/  rev | Length  -ATG (0) |
| --- | --- | --- |

| **pT3O** | GGGGACAAGTTTGTACAAAAAAGCAGGCTTAGAAACGATAGATTTCTTTTGATGCCT | -1982 |
| --- | --- | --- |
| **CRO_T113994** | GGGGACCACTTTGTACAAGAAAGCTGGGTAAATCAAAGCTTACTAATTATGAATCTC | 0 |
| **pT3R** | GGGGACAAGTTTGTACAAAAAAGCAGGCTTAAGGCTTCCTTTCTTTTGTTGTTTC | -1992 |
| **CRO_T124298** | GGGGACCACTTTGTACAAGAAAGCTGGGTATAAGGAAAAGACAAGCAAAGCCA | 0 |
| **pD4H** | GGGGACAAGTTTGTACAAAAAAGCAGGCTTAGATCTTATCTCCTAAACCCTAAACC | -1241 |
| **CRO_T127167** | GGGGACCACTTTGTACAAGAAAGCTGGGTATAGGATTACAGACAGCGGAC | -79 |
| **pDAT** | GGGGACAAGTTTGTACAAAAAAGCAGGCTTAGCATCCAACCATAAAACATTACCCT | -934 |
| **CRO_T120021** | GGGGACCACTTTGTACAAGAAAGCTGGGTAATTCAGACCCCAACACACAAC | -17 |
| **pGO** | GGGGACAAGTTTGTACAAAAAAGCAGGCTTATAATTTTTTATATGCTCGAGTACACT | -2000 |
| **CRO_T127440** | GGGGACCACTTTGTACAAGAAAGCTGGGTATGTGTTTTGTTCAAGAGAGGAGAC | 0 |
| **pTAT** | GGGGACAAGTTTGTACAAAAAAGCAGGCTTATGATAAATAATTCTTTCGCTTTTGCA | -2000 |
| **CRO_T120026** | GGGGACCACTTTGTACAAGAAAGCTGGGTATTTGCTCATGCCATTATTATGAACTCA | 0 |
| **pMAT** | GGGGACAAGTTTGTACAAAAAAGCAGGCTTAAAAGTAATTGTTATCAATTTATTGA | -2000 |
| **CRO_T120028** | GGGGACCACTTTGTACAAGAAAGCTGGGTATTTGCTCAATATGCTGCTTTCCA | 0 |

**Table S4** List of Genebank IDs of all cloned TFs and promoter fragments.

| **Genebank accession** | **CRO_ID** | **Name** |
| --- | --- | --- |
| MT414967 | CRO_T127167 | pD4H |
| MT414968 | CRO_T120021 | pDAT |
| MT414969 | CRO_T127440 | pGO |
| MT414970 | CRO_T120028 | pMAT |
| MT414971 | CRO_T113994 | pT3O |
| MT414972 | CRO_T124298 | pT3R |
| MT414973 | CRO_T120026 | pTAT |
| MT414974 | CRO_T100680 | MYB type TF |
| MT414975 | CRO_T101037 | TCP type TF |
| MT414976 | CRO_T103361 | TPR type TF |
| MT414977 | CRO_T105905 | bHLH type TF |
| MT414978 | CRO_T107533 | BIS3 |
| MT414979 | CRO_T109258 | bHLH type TF |
| MT414980 | CRO_T110248 | bHLH type TF |
| MT414981 | CRO_T110359 | ORCA5 |
| MT414982 | CRO_T110361 | ORCA4 |
| MT414983 | CRO_T110364 | ORCA6 |
| MT414984 | CRO_T110868 | AP2/B3 type TF |
| MT414985 | CRO_T112238 | GATA type TF |
| MT414986 | CRO_T112897 | CCT type TF |
| MT414987 | CRO_T115526 | ZF type TF |
| MT414988 | CRO_T116348 | HF type TF |
| MT414989 | CRO_T116828 | WRKY type TF |
| MT414990 | CRO_T117719 | AP2/ERF type TF |
| MT414991 | CRO_T118079 | bHLH type TF |
| MT414992 | CRO_T118542 | ARR type TF |
| MT414993 | CRO_T122239 | NYF-YA type TF |
| MT414994 | CRO_T123350 | bHLH type TF |
| MT414995 | CRO_T124980 | bHLH type TF |
| MT414996 | CRO_T126752 | ANAC type TF |
| MT414997 | CRO_T127367 | AP2/ERF type TF |
| MT414998 | CRO_T127378 | ANAC type TF |
| MT414999 | CRO_T128811 | C2H2 ZF type TF |
| MT415000 | CRO_T129607 | OFP type TF |
| MT415001 | CRO_T130446 | BBX type TF |
| MT415002 | CRO_T131234 | MYB96 |
| MT415003 | CRO_T131857 | AP2/ERF type TF |
| MT415004 | CRO_T132027 | bHLH type TF |
| MT415005 | CRO_T132285 | BBX type TF |
| MT415006 | CRO_T133384 | TLP type TF |
| MT415007 | CRO_T134504 | MYB type TF |
| MT415008 | CRO_T134559 | Homeodomain type TF |
| MT415009 | CRO_T137796 | MYB96b |
| AJ238740 | CRO_T110365 | ORCA2 |
| AJ251249 | CRO_T110360 | ORCA3 |

**Table S5** Oligonucleotide primers used for qPCR

| **Gene name or TF family name** | **Gene ID** | **fw** | **rev** |
| --- | --- | --- | --- |
| **GES** | CRO_T119458 | GGGCAAGGTGTCACTGAAGA | CCCAAATCATCCCAAAGGCG |
| **G8O** | CRO_T133061 | TGCTTTGGCATTCAAACCCG | GCAAATTCTTCGGCCAGCAC |
| **8HGO** | CRO_T107879 | GCATTCCTGGACACGAAGGA | GCATTCCCCACATTCTCCCA |
| **IS** | CRO_T130026 | AGTAGTAGGAGTCACCGGCA | CTTGCCACGCCGTATACCTT |
| **IO** | CRO_T138994 | TCAAGTCCAAATACGGGCCG | CGGCGGAAACTGCATTTTGA |
| **7DLGT** | CRO_T106494 | GCAGAGGGAGTTCTCAAGGC | TGGCCAATGCACATTCTTGG |
| **7DLH** | CRO_T131714 | GGTCTTGAATCACCCTGCCA | TGGGACACCAGCCACAATAC |
| **LAMT** | CRO_T103723 | TGGAAGCCCACCCAATGAAA | ACTGCCTTGGCTGCATCAAT |
| **SLS1** | CRO_T113655 | CCCTGCAGGAACACAAGTGA | GCATTGGCAACTCCATCAGC |
| **SLS2** | CRO_T109472 | CCACTGGAGTTTTGCTCACA | TTATTCCTGCCAAAGGCTTC |
| **TSB** | CRO_T127328 | TGTGTCGGTGGTGGTTCAAA | CCAAAACCAGCAGCTTCCAC |
| **TDC** | CRO_T125328 | GGTCGCTGAAACTTTGGCTC | ATTTTGCCCATTGCGACGTC |
| **STR1** | CRO_T125329 | CGTCCAAGATGGCCGAGTTA | CTGGATCGGTGCTGTTCTCA |
| **SGD** | CRO_T111319* | GGATGGAGCCTCTCAATGAA | CACCCGTTGTTAATGGCTCT |
| **GS1** | CRO_T113154 | TCAAGGCGTAAGGAGAAGGA | AGTACCTGCCAGAGCCTTCA |
| **GS2** | CRO_T113153 | GGTCTTGGTTCCATTGCTGT | TGCTCTTCAATGGCTTCCTT |
| **GO** | CRO_T127440 | TCCTTTGATGCACGTGAAAA | TTACTGACCGATCTGCAACG |
| **Redox1** | CRO_T129272 | GAAGTGACGGAAGTGGGGAACAAA | TCGCATTCGCCACATGAGTCAA |
| **Redox2** | CRO_T132421 | TCGCTTGGGGAAGTAATGCTGT | TGAGACTTGCTCCTTGCTCGTA |
| **SAT** | CRO_T109653 | GGATGGGGAAAGCCTGTTTCTGTT | CTTCAGCCATGCTGATCCATGCTT |
| **PAS** | CRO_T113148 | CTTCACTCCCATGTCCAATCT | CGATAGGATAAGCCCTCGTAATC |
| **DPAS** | CRO_T129267 | GAAATAGCGGCATCGACAAAC | GCTGGGAGTGGTGCTAATAA |
| **TS** | CRO_T110304 | TGCTCCTGGTGGAAATGATAACCC | AATCAGCAACCTCGAGCAACCA |
| **T16H1** | CRO_T110599 | AGGCTTCATCCACCAGTTCC | CCTTCCGATAGCCCATGCAT |
| **T16H2** | CRO_T110598 | GATCAACTCACAGTGGCAGTC | GACTTGAGGACTTGTGATTGGC |
| **16OMT** | CRO_T110596 | CTCTTGTGCCCCCAGTTCAC | CGACTTGATGGGGTAAGGGA |
| **T3O** | CRO_T113994 | CCCATGTGATGAGCAAAGCG | AACAATGGAGCAGGAGGGTG |
| **T3R** | CRO_T124298 | GAAGGGCTACAGGGGAACAC | CACCCACAATTTCATGCCCG |
| **T19H** | CRO_T119486 | TGAGTTGCCAAATGGAGTCA | CAAACGAGAGAGGGTTTTGG |
| **TAT** | CRO_T120026 | CCACAGACTTGGTCCTTCCC | ATGACTGAGCTGCACGATCA |
| **MAT** | CRO_T120028 | ATCGAAGGCCATTGAGTTTG | GCTGCTGATTTCCCTGCTAC |
| **TEX1** | CRO_T122015 | AGGAAGGCGTATTTGTCCGG | TCTTCTGGCTTCATTCCGCC |
| **TEX2** | CRO_T120417 | CCGTTGGCGCAACTTTTGTA | ACCCAAAATCTCCGCCATGT |
| **THAS1** | CRO_T113666 | TGTGGCAAATGTGAAATGTGT | ATTTGAACATGCCCCGTAAT |
| **AS** | CRO_T116107 | GAATCGGCCAACTTTCAGCC | AGTAGTTTGTGGCCTGTCCG |
| **HYS** | CRO_T140758 | AAGTTGGTGTAGGGGGCTTC | CAAAATGGCCATCTGTTGATT |
| **VAS** | CRO_T112618 | GTGTTTGTCCAGGGTTTGCC | GCAGCAAAGCGAGTTTCACA |
| **NMT** | CRO_T111273 | TTCGTGAGATGGTTCGGGTG | CGGCGCCGTCACATATTTTT |
| **CS** | CRO_T139139 | TGGGGCTGGCTTTTGTCTAGAATC | TAAGCTGCGGGTAAAAGGTGCTCT |
| **TPT2** | CRO_T137586 | CCAATGTCACCGGTGCATTC | TCCACCTGTTTTCCGTCCAG |
| **D4H** | CRO_T127167 | ACAGCTGATCACGAACGACA | TCTTGGCGAAACCCCTTCTT |
| **DAT** | CRO_T120021 | GGTTTCAATTTATTTCTCACGTAC | AACTATCAGAAAGGTAAGCATCGA |
| **PRX1** | CRO_T141131 | TAACGGGGAATCAAGGTGAA | AATTTCAGCAGCCTCTTCCA |
| **V19H** | CRO_T135744 | GCTCAGAAACATGGGCCTCT | TTGGCCATTTCGGGTGATGA |
| **BIS1** | CRO_T107535 | ATGGAATCAGTGGTGCTAGTGA | TTCAATTTCAGGGAGCTGTGAC |
| **BIS2** | CRO_T107539 | TGTGCAGTTCTTGGTCAAGACT | AGTGGAATTTGGATCCAAGTTTG |
| **MYC2a** | CRO_T124533 | AGTGGTGAAGGAGGCAGAGA | ATGGCTCTTCCCTTCCATTT |
| **N2227** | CRO_T133527/8* | GGTTGCTCTTCATTACGGATTT | TGCAGCATAGTAATGGTTTTGC |
| **SAND** | CRO_T130722 | CAGTTCCACAATGCTTTCTGAC | GGGACTGATCAATCGAAGTAGC |
| **NPF2.4** | CRO_T131105 | ACAAATATTGGATGAGCCCAAG | CCCTGTTTTTATTCTTGGAAGC |
| **NPF2.5** | CRO_T131100 | AAAGAATGGGAATTGGAATGG | TTCGTGGCTCAAAGCCTAAT |
| **NPF2.6** | CRO_T131101 | CGATTATCGATCAATTGCAAGG | CTTCTGAGGTACCCAAGATTACG |
| **NPF2.9** | CRO_T105710 | CCTCCTCCATTTTCATTTTCTG | GATGCGGTTGGATTTGTGAGTC |
| **ORCA2** | CRO_T110365 | GTTGCGGGAGAACAAGAAGA | AACGCCACGGTACCTAATCC |
| **ORCA3** | CRO_T110360 | CGGAAAGCTGTCAGGAGGAT | CGTCTAACGCCCTTATACCGG |
| **ORCA3 alt** | CRO_T110360 | CGGAAAGCTGTCAGGAGGAT | TCGTATGTACCCAACCAAATCC |
| **ORCA4** | CRO_T110361 | TCAGCCTCCGATCTAGCTCT | GATTCGAGCTGCTGTCGTCA |
| **ORCA5** | CRO_T110359 | CCACGGAGGCTAATAGGGGA | TTCTTCTTCCCGGCCATGTC |
| **ORCA6** | CRO_T110364 | TTATTATTAATTCCCCCAAATGCAGGT | TGCTAATGCAGTCAACTCTTGT |
| **CER1** | CRO_T110442 | GCTCGCTCCTTCATCTCTCC | CCTGCTACTCTGCTTGCACT |
| **BIS3** | CRO_T107533 | TCCTCATGATTAACAATGATGATGAA | TTGACCAACAGTTGCTGCAC |
| **GATA** | CRO_T112238 | GAGGAGGGGGATATGGCTCT | GGGGGTGTAGTTTTGGTGGT |
| **BBX** | CRO_T132285 | CACATTCTGCTAATCGCGGC | ACAACGAGGTTCCTTGCTGT |
| **CCT** | CRO_T112897 | ACTCTCCGACTTCAGAGGCT | TTCCCCCGCTTTCTGAGAAC |
| **AP2/ERF** | CRO_T127367 | GGGCCTCGTTCTGATTCCAA | GGTTGCTGCGACGTTTGATT |
| **BBX** | CRO_T130446 | TATGGGCGCGGTATTCAACA | CATGCACAGTGTCCTGCAAC |
| **bHLH** | CRO_T118079 | GGCTTGAATGTCAGCTGAGC | TGGAGCAAACCACTTGTTCTG |
| **C2H2 ZF** | CRO_T128811 | TGGCAACATCACTTGGCCTC | GCGGTGGACATTCATATGGC |
| **bHLH** | CRO_T132027 | AAGGAAGAAGGCGTCGTCAA | AGAGCAGCCATTTCTTGCCT |
| **bHLH** | CRO_T109258 | ACGGACACAATGGTAAAGCT | GGCTACCTGAGAAAGCAGTGA |
| **ARR** | CRO_T118542 | TGGGTTTGCTGGATGAGGAA | AGCCAGTCATTCCAGGCATG |
| **TPR** | CRO_T103361 | CGGTCAGCAGAAGAAGCAGA | TCAACCCACTTCTCCAGCAC |
| **AP2/ERF** | CRO_T117719 | ATGAGCGAATCGACGAGGAC | CGGACGAGTCGTAGCAACTT |
| **TLP** | CRO_T133384 | CTTACCCCGCTGTTTCTGGT | TCACTGATGCCACTGTCACC |
| **NYF-YA** | CRO_T122239 | TACGAGCATCTTCAGTGGCG | TCCAACATGCGCGTCCATAT |
| **WRKY** | CRO_T116828 | CAAGGGAGAGATGGGTGCTC | TGGATAGGGGGAGCCTTTGA |
| **bHLH** | CRO_T123350 | GTCTACCATGGTTCCTGGCC | TGCGTTCTTGGAGCTGTTCT |
| **bHLH** | CRO_T105905 | TGCAGCAAAAATGGCACCAA | GACTGAGCTGCTCCCTTCTC |
| **ZF** | CRO_T115526 | CCAAAACCACCAACACCACC | CGGAATTCTCCACAACCCCA |
| **OFP** | CRO_T129607 | TGGAAGCCCCAAAACCCTTT | TCCGGAAGAGGGTCAATCGA |
| **AP2/ERF** | CRO_T131857 | GAGATCCAGCGAAGAACGGA | CCGATAAGCTGCTCGATCGT |
| **Homeodomain** | CRO_T134559 | AGGATGGTGGAATTGGCTGG | GCCACTTCCACCAATGCATC |
| **HF** | CRO_T116348 | TCCACGAGAACAAACGCCTC | CAGAACTAGAGTGGGCAGCC |
| **MYB** | CRO_T100680 | CAACCTGACCGAAGCAGTCT | TGCAGCATTCCAGAAGCTCA |
| **TCP** | CRO_T101037 | CCGCATTGTTCGATCCACAG | CTCACCCGTCGATCTCTTGG |
| **AP2/B3** | CRO_T110868 | CGCCCCTTTTTGTATCGACG | CCTTCCAGCAATCGCCTTCT |
| **ANAC** | CRO_T126752 | CGTGACCCTTTCTTTTGGCG | TGTTGGGGAGACTGCCTTTC |
| **MYB** | CRO_T134504 | CACAAGAACGCCAACCCAAG | TGGAACGCCTCTTGAGCTTT |
| **ANAC** | CRO_T127378 | GGAAATGTGCGTCTCAACCG | CGCCATACCAGGAAGATCCC |
| **bHLH** | CRO_T110248 | CGGGTTTGGAGAGGGTTCAG | CCTCAGTGCAGCCCTCTTTT |
| **bHLH** | CRO_T124980 | TTCAGCGGATGACATAGCGG | TAGCCCTCCTCCTCCTCCTA |
| **MYB96** | CRO_T131234 | AATGATCAGCAAGGAGGCGG | AATTTACAGCTGCTTCATCCACA |
| **MYB96b** | CRO_T137796 | TGCATCTAAAGGCCAGTGGG | CCAGTGTCAAGGCATCCTGT |
| **GUN4** | CRO_T139058 | GCCATCCTGCTTTTGAAGGC | GCAGCAGATAAACCCCCACT |
| **CHLI1** | CRO_T103461 | ATTTCACATCCTGCCCGGTT | TGTGCATGCATTCCAAACCG |
| **GLU1** | CRO_T100999 | ATTTTGCCCTGAGGATGCCA | CACAGCAAAACGTTCCCCAG |
| **PLT6** | CRO_T121364 | GCACAAGGGGCTTCAATTGG | TAGCTCCCGCCAATGACAAG |

*ID not correct in Reference Genome, manually corrected based on other transcriptomes.

**Tables S6-S10** MIA levels of different flower petal infiltration experiments as indicated. Values labelled with different letters indicate statistically significant differences (P < 0.05 calculated by ANOVA with Tukey’s correction for multiple comparisons.

**Table S6** MIA levels upon overexpression of *ORCAs*.

|  | ***p35S::GUS* ± SEM** |  | ***p35S::ORCA2* ± SEM** |  | **p35S::ORCA3 ± SEM** |  | ***p35S::ORCA4* ± SEM** |  | ***p35S::ORCA5* ± SEM** |  | ***p35S::ORCA6* ± SEM** |  |
| --- | --- | --- | --- | --- | --- | --- | --- | --- | --- | --- | --- | --- |
| Loganin | 17625.38 ± 2167.60 | a | 16810.58 ± 1795.08 | a | 17077.22 ± 2058.43 | a | 15206.09 ± 1843.48 | a | 15801.42 ± 518.64 | a | 18963.89 ± 922.95 | a |
| Secologanin | 171180.14 ± 14499.32 | a | 168473.28 ± 16717.87 | a | 179176.07 ± 8601.95 | a | 142409.56 ± 9877.43 | a | 171930.23 ± 10586.70 | a | 184295.65 ± 5698.73 | a |
| Strictosidine | 3258939.47 ± 269524.31 | a | 3893459.39 ± 390430.99 | ab | 4310460.85 ± 243889.15 | ab | 5367515.97 ± 589468.00 | b | 3268640.37 ± 262797.49 | a | 2851583.99 ± 167540.26 | a |
| Strictosidine secologanoside | n.d. | a | 143.26 ± 100.39 | a (n.d in 50%) | n.d. | a | 1180.63 ± 504.45 | b | n.d. | a | 27.83 ± 27.83 | a (n.d. in 75%) |
| Unknown 2 (strictosidine aglycone isomer) | 61474.47 ± 2087.38 | a | 87104.91 ± 8849.05 | ab | 115825.82 ± 10617.37 | b | 68817.73 ± 10988.09 | a | 73601.77 ± 6369.61 | a | 84033.37 ± 5165.24 | ab |
| Strictosidinic Acid | 281585.82 ± 21757.76 | a | 496624.55 ± 69389.41 | a | 1276110.50 ± 133520.65 | b | 1151321.40 ± 242493.25 | b | 422227.44 ± 35179.04 | a | 318509.81 ± 17158.80 | a |
| Serpentine | 2324902.62 ± 197161.98 | a | 1982141.11 ± 208396.21 | a | 2475673.54 ± 195546.70 | a | 2077741.85 ± 534684.92 | a | 2339104.21 ± 545604.89 | a | 2298310.64 ± 117079.01 | a |
| Unknown MIA (serpentine isomer) | 647187.00 ± 17487.69 | a | 661450.03 ± 64300.37 | a | 738198.07 ± 50320.83 | a | 735765.70 ± 177241.83 | a | 571635.81 ± 130560.67 | a | 740626.38 ± 57117.53 | a |
| Geissoschizine | 484273.85 ± 56276.89 | a | 1255979.65 ± 126167.58 | b | 2543671.29 ± 208974.63 | c | 852073.67 ± 142919.64 | ab | 965583.71 ± 93937.07 | ab | 868932.16 ± 44977.13 | ab |
| Unknown MIA (geissoschizine isomer 1) | 1004629.00 ± 107253.68 | a | 3021401.96 ± 297899.88 | b | 6758134.73 ± 574717.66 | c | 1881703.04 ± 360679.97 | ab | 2393413.70 ± 242716.16 | ab | 2077270.86 ± 116204.00 | ab |
| Unknown MIA (geissoschizine isomer 2) | 59467.49 ± 6427.65 | a | 120020.73 ± 15061.36 | bc | 155534.40 ± 11283.00 | c | 83274.46 ± 19253.96 | ab | 52809.09 ± 3870.09 | a | 65797.87 ± 7522.57 | a |
| Unknown 1 (isositsirikine isomer) | 313491.34 ± 13274.63 | a | 425925.05 ± 43974.76 | a | 680951.70 ± 64148.28 | b | 438919.85 ± 68368.05 | a | 401676.62 ± 36671.16 | a | 426616.49 ± 16766.70 | a |
| Isositsirikine | 278013.65 ± 16846.72 | a | 373626.86 ± 36350.64 | a | 422133.20 ± 30465.74 | a | 423987.85 ± 67857.06 | a | 280666.38 ± 25145.25 | a | 371290.32 ± 17316.57 | a |
| Perivine | 106630.91 ± 4362.20 | a | 191979.35 ± 19670.21 | ab | 168940.09 ± 17215.88 | ab | 231511.27 ± 46723.09 | b | 125292.97 ± 11991.40 | a | 147817.61 ± 10341.48 | ab |
| Akuammicine | 217149.15 ± 14251.41 | a | 431327.39 ± 44961.30 | b | 301704.50 ± 12925.86 | ab | 494667.49 ± 96705.10 | b | 212423.58 ± 21861.19 | a | 267564.80 ± 16382.93 | a |
| O-acetylstemmadenine | 285384.26 ± 33981.10 | a | 396035.90 ± 32906.25 | ab | 354546.47 ± 12288.51 | ab | 408649.32 ± 33199.65 | b | 291303.13 ± 16727.28 | a | 331319.86 ± 17340.03 | ab |
| 16-hydroxytabersonine | 236505.11 ± 29528.00 | a | 980025.80 ± 100088.65 | b | 709016.29 ± 49945.59 | ab | 1617549.55 ± 256250.18 | c | 351625.97 ± 35739.56 | a | 378565.15 ± 35153.62 | a |
| 16-hydroxytabersonine glucoside | 21106.33 ± 3070.29 | a | 59935.16 ± 5721.10 | b | 56817.49 ± 7688.34 | ab | 114752.23 ± 18055.98 | c | 24312.50 ± 2235.73 | a | 22916.85 ± 2618.79 | a |
| Catharanthine | 9502483.18 ± 527452.99 | a | 10185635.15 ± 954787.04 | a | 10539400.36 ± 436607.75 | a | 11204859.70 ± 1504637.03 | a | 8055529.92 ± 1120877.53 | a | 11840581.16 ± 854946.78 | a |
| Unknown MIA (catharanthine isomer 1) | 5817169.65 ± 261033.71 | a | 6061133.04 ± 614549.65 | a | 7119639.59 ± 500279.02 | a | 6458861.81 ± 954773.77 | a | 5475778.74 ± 656023.90 | a | 7510768.69 ± 615668.52 | a |
| Unknown MIA (catharanthine isomer 2) | 3651172.35  ± 205625.52 | a | 3981720.74 ± 405962.48 | a | 4591785.48 ± 318612.76 | a | 4171041.45 ± 662515.76 | a | 3416365.32 ± 389145.48 | a | 4618904.46 ± 376395.55 | a |
| Unknown MIA (catharanthine isomer 3) | 597799.07  ± 11818.06 | a | 655548.05 ± 68935.66 | a | 794130.57 ± 68009.74 | a | 746452.70 ± 111931.08 | a | 552713.43 ± 62861.96 | a | 800013.94 ± 52882.85 | a |
| Desacetoxyvindoline | 617731.34  ± 87917.14 | ab | 880009.31 ± 70912.99 | b | 647521.57 ± 74506.27 | ab | 606161.19 ± 48638.01 | ab | 597575.29 ± 16299.77 | a | 871300.66 ± 43245.92 | ab |
| Deacetylvindoline | 299605.96  ± 10440.08 | a | 320925.21 ± 27968.21 | a | 331481.32 ± 17608.41 | a | 278500.01 ± 19193.34 | a | 285962.34 ± 22800.68 | a | 367904.07 ± 17380.97 | a |
| Vindoline | 9351770.86  ± 777129.99 | a | 10071645.17 ± 746508.98 | a | 11196624.74 ± 1053443.15 | a | 9325684.41 ± 538140.76 | a | 9458499.26 ± 869042.83 | a | 11205775.39 ± 641733.96 | a |
| Anhydrovinblastine | 315768.67  ± 17360.80 | a | 382207.33 ± 79352.70 | a | 391216.31 ± 31360.38 | a | 313096.30 ± 68086.04 | a | 243123.80 ± 41568.43 | a | 381113.21 ± 33507.31 | a |
| Vinblastine | 66451.18  ± 9908.89 | a | 92626.23 ± 18912.39 | a | 86120.06 ± 8138.94 | a | 70784.57 ± 22432.29 | a | 54781.73 ± 11424.54 | a | 78097.39 ± 20789.89 | a |
| Vincristine | 137836.62  ± 11676.38 | a | 150443.44 ± 15443.39 | a | 159327.51 ± 6368.79 | a | 120598.52 ± 26465.96 | a | 106538.64 ± 14690.10 | a | 143140.55 ± 15745.80 | a |
| Desacetoxyvindorosine | 6304.61  ± 1171.30 | a | 10647.66 ± 892.74 | ab | 6832.81 ± 1165.50 | a | 8631.06 ± 1343.06 | ab | 7049.38 ± 693.49 | a | 12712.49 ± 1340.31 | b |
| Desacetylvindorosine | 17748.33  ± 1093.51 | a | 19353.01 ± 1775.73 | a | 19273.81 ± 1722.43 | a | 18358.39 ± 2715.84 | a | 20048.62 ± 2283.52 | a | 25428.98 ± 1315.57 | a |
| Demethoxyvindoline = vindorosine | 433708.03  ± 18099.78 | a | 504526.26 ± 50526.80 | a | 534409.28 ± 39973.28 | a | 521432.17 ± 94984.45 | a | 416491.63 ± 65335.37 | a | 619774.63 ± 48058.47 | a |
| Vincadifformine | 3510.84  ± 197.70 | a | 5777.01 ± 899.43 | ab | 4524.39 ± 531.70 | ab | 10041.78 ± 2745.91 | b | 2710.44 ± 771.11 | a | 5298.57 ± 1163.96 | ab |
| 16‑Hydroxyvincadifformine | 2628.81  ± 276.42 | a | 7880.72 ± 1207.24 | a | 17880.15 ± 2712.05 | b | 4279.05 ± 1487.13 | a | 5257.53 ± 876.93 | a | 5117.18 ± 697.47 | a |
| Minovincinine | 1537.25  ± 245.75 | a | 2710.81 ± 292.72 | a | 5332.97 ± 868.25 | b | 3413.38 ± 802.19 | ab | 1274.11 ± 176.90 | a | 2060.31 ± 180.76 | a |
| 19-hydroxytabersonine | 40201.69  ± 3409.83 | a | 57404.81 ± 7499.03 | a | 56188.74 ± 6908.92 | a | 49478.63 ± 6201.54 | a | 41563.00 ± 2818.75 | a | 56217.78 ± 3284.85 | a |
| 19-O-acetyltabersonine | 83672.73  ± 7433.12 | a | 215233.42 ± 16510.63 | b | 167834.01 ± 12321.07 | c | 300473.30 ± 35168.08 | d | 116404.38 ± 9199.83 | ac | 129030.59 ± 6893.00 | ac |
| 16-hydroxy-19-O-acetyltabersonine | 41.43 ± 16.00 | a | 152.11 ± 39.56 | a | 19.37 ± 12.29 | a | 55.73 ± 36.25 | a | 91.94 ± 38.29 | a | 146.98 ± 53.37 | a |
| 16-hydroxylochnericine | 181233.31  ± 7502.30 | a | 530591.48 ± 56142.18 | a | 384911.78 ± 29794.10 | a | 1208983.16 ± 212690.52 | b | 237525.87 ± 26963.13 | a | 243321.33 ± 11289.52 | a |
| 16-hydroxylochnericine glucoside | 16762.93  ± 3093.86 | a | 58792.03 ± 5408.12 | a | 35975.42 ± 4500.69 | a | 225106.48 ± 35746.05 | b | 20360.30 ± 2786.09 | a | 8507.11 ± 1746.77 | a |
| Hörhammericine | 30101.62  ± 1432.96 | a | 44844.42 ± 4560.43 | a | 52316.87 ± 5020.81 | a | 54981.20 ± 11522.39 | a | 32818.98 ± 2988.17 | a | 38820.64 ± 1952.01 | a |
| 16-hydroxyhörhammericine | 71.52 ± 22.88 | a | 363.28 ± 123.21 | ab | 870.56 ± 296.20 | ab | 998.66 ± 364.73 | b | 115.80 ± 14.86 | ab | 99.56 ± 27.42 | a |
| 16-methoxyhörhammericine | 8104.56  ± 582.99 | a | 29086.91 ± 2857.31 | a | 117435.00 ± 20099.66 | b | 16640.37 ± 2720.54 | a | 42251.69 ± 2803.09 | a | 12349.81 ± 533.10 | a |
| Vandrikidine | 139663.86 ± 4134.40 | a | 155217.11 ± 16344.87 | a | 161911.98 ± 10945.14 | a | 148383.13 ± 26667.18 | a | 110917.30 ± 14011.74 | a | 178893.60 ± 13500.38 | a |

**Table S7** MIA levels upon overexpression of *ORCA3* mutants.

|  | ***p35S::GUS* ± SEM** |  | ***p35S:ORCA3* ± SEM** |  | ***p35S::ORCA3^K101R^* ± SEM** |  | **p35S::ORCA3^P107L^ ± SEM** |  | **p35S::ORCA3^K101RP107L^ ± SEM** |  | **p35S::ORCA3^K101RK110R^ ± SEM** |  | **p35S::ORCA3^K101RK110T^ ± SEM** |  |
| --- | --- | --- | --- | --- | --- | --- | --- | --- | --- | --- | --- | --- | --- | --- |
| Loganin | 15870.77 ± 1458.96 | ab | 12864.96 ± 1192.62 | a | 14891.98 ± 883.49 | ab | 19717.96 ± 1211.98 | ab | 16149.62 ± 843.79 | b | 16490.56 ± 1042.61 | ab | 15609.33 ± 1390.84 | ab |
| Secologanin | 69445.44 ± 3537.49 | a | 68481.76 ± 3403.10 | a | 71583.40 ± 4175.36 | a | 68643.41 ± 3326.18 | a | 69928.17 ± 796.63 | a | 74543.71 ± 3737.52 | a | 69267.73 ± 2913.20 | a |
| Strictosidine | 2491704.50 ± 220688.23 | a | 2846468.72 ± 148459.61 | ab | 2905514.79 ± 296430.80 | ab | 3757294.17 ± 431858.18 | b | 3182636.10 ± 107286.62 | ab | 3261658.62 ± 143333.09 | ab | 3068013.90 ± 170395.59 | ab |
| Strictosidine secologanoside | 9.69 ± 9.69 | a | 39.89 ± 28.45 | a | 70.19 ± 46.78 | a | 85.82 ± 41.42 | a | 65.64 ± 23.14 | a | 410.58 ± 167.91 | b | 47.60 ± 47.60 | a |
| Unknown 2 (strictosidine aglycone isomer) | 102405.25 ± 13429.58 | a | 118696.97 ± 8931.98 | a | 170719.70 ± 11907.70 | b | 139602.81 ± 14258.81 | ab | 127566.68 ± 4744.53 | ab | 172485.40 ± 11443.52 | b | 129926.43 ± 10934.42 | ab |
| Strictosidinic Acid | 38752.78 ± 5734.89 | a | 346478.86 ± 25775.03 | b | 693798.12 ± 92274.52 | c | 451479.73 ± 33725.08 | bd | 603865.82 ± 41936.16 | cd | 641522.64 ± 55062.92 | cd | 257901.12 ± 13609.91 | b |
| Serpentine | 2212345.63 ± 374693.27 | ab | 1675881.74 ± 219310.15 | a | 3556228.66 ± 780828.03 | b | 2753236.80 ± 338911.27 | ab | 2869463.86 ± 189678.55 | ab | 2597008.84 ± 156549.08 | ab | 2314128.20 ± 293452.54 | ab |
| Unknown MIA (serpentine isomer) | 747125.34 ± 110960.56 | ab | 597302.38 ± 75465.39 | a | 1190529.90 ± 198473.19 | b | 1063275.12 ± 127753.32 | ab | 917204.08 ± 53889.42 | ab | 1009392.84 ± 68169.01 | ab | 870255.96 ± 98521.43 | ab |
| Geissoschizine | 1340801.46 ± 201855.89 | a | 6714390.87 ± 269268.96 | be | 8590381.24 ± 481830.05 | ce | 4961214.37 ± 440360.75 | bd | 4333031.51 ± 140386.18 | d | 10251251.59 ± 675868.73 | c | 6172585.80 ± 569228.45 | bd |
| Unknown MIA (geissoschizine isomer 1) | 855433.79 ± 131489.70 | a | 4446439.46 ± 196721.59 | bc | 5777104.99 ± 342575.90 | ce | 3398546.65 ± 311671.66 | bd | 2752421.45 ± 88262.99 | d | 6892975.47 ± 445481.03 | e | 4200659.74 ± 411108.28 | b |
| Unknown MIA (geissoschizine isomer 2) | 60771.73 ± 13974.72 | a | 126852.38 ± 9214.37 | b | 259793.66 ± 14916.38 | c | 154433.19 ± 19183.68 | b | 159054.48 ± 6013.53 | b | 235498.41 ± 8204.45 | c | 134505.10 ± 17189.43 | b |
| Isositsirikine | 225522.02 ± 29611.46 | a | 259818.09 ± 17884.46 | ac | 421237.32 ± 34205.50 | b | 380875.60 ± 44377.34 | bc | 405217.16 ± 15165.71 | b | 437540.27 ± 28321.47 | b | 304004.78 ± 20724.26 | a |
| Unknown 1 (isositsirikine isomer) | 372859.26 ± 44916.38 | a | 502353.99 ± 34064.79 | b | 811497.66 ± 77791.17 | cd | 652790.17 ± 73439.55 | bcd | 586669.12 ± 23184.49 | ab | 850643.18 ± 55711.20 | d | 572485.94 ± 46757.00 | abc |
| Perivine | 107276.55 ± 14116.95 | ac | 118061.78 ± 10883.12 | ac | 226905.69 ± 13662.81 | b | 176003.82 ± 13725.48 | bc | 197001.11 ± 8529.24 | bc | 202893.73 ± 14417.91 | b | 144921.25 ± 11817.69 | c |
| Akuammicine | 182818.89 ± 31162.33 | a | 226540.97 ± 21275.02 | ab | 495310.09 ± 29322.20 | c | 338896.06 ± 32715.98 | bd | 477632.31 ± 12708.58 | c | 421485.92 ± 30392.35 | cd | 253736.88 ± 27036.65 | ab |
| O-acetylstemmadenine | 155778.26 ± 16492.71 | a | 197183.36 ± 7772.73 | ab | 266716.01 ± 17228.75 | b | 225984.02 ± 14768.30 | ab | 255672.33 ± 7109.68 | b | 272375.70 ± 23637.06 | b | 242422.61 ± 20731.94 | b |
| 16-hydroxytabersonine | 139850.08 ± 26678.16 | a | 473944.09 ± 39684.32 | b | 820746.25 ± 19538.23 | c | 1121219.85 ± 80115.30 | d | 1765345.30 ± 93684.63 | e | 926990.57 ± 69380.26 | cd | 482094.73 ± 48926.74 | b |
| 16-hydroxytabersonine glucoside | 12181.32 ± 2315.99 | a | 38736.11 ± 4627.69 | b | 71702.11 ± 7653.54 | c | 79750.27 ± 4025.20 | c | 188769.17 ± 9185.22 | d | 84504.62 ± 5867.19 | c | 34703.94 ± 2010.77 | ab |
| Catharanthine | 7827404.24 ± 1157412.25 | ab | 6461724.23 ± 603424.79 | a | 11445467.34 ± 1314038.95 | b | 10871245.25 ± 1429655.87 | ab | 9269471.35 ± 404395.32 | ab | 10169822.82 ± 740937.60 | ab | 9544521.27 ± 868385.03 | ab |
| Unknown MIA (catharanthine isomer 1) | 4675344.47 ± 508500.49 | ab | 3924921.19 ± 289243.66 | b | 5972668.89 ± 638619.11 | a | 6050521.54 ± 495682.65 | a | 4915187.19 ± 204238.95 | ab | 5554826.41 ± 392778.00 | ab | 5159076.44 ± 283324.84 | ab |
| Unknown MIA (catharanthine isomer 2) | 3434250.89 ± 418890.54 | a | 2923654.64 ± 247024.68 | a | 4195981.83 ± 406126.08 | a | 4296896.47 ± 381370.45 | a | 3682735.71 ± 141818.72 | a | 4061934.51 ± 241342.34 | a | 3613899.63 ± 222877.46 | a |
| Unknown MIA (catharanthine isomer 3) | 779360.78 ± 88000.19 | ab | 648889.45 ± 56062.97 | a | 951707.83 ± 90936.85 | ab | 1032899.35 ± 71090.50 | b | 812171.66 ± 34075.56 | ab | 909199.58 ± 56347.59 | ab | 836293.76 ± 55638.03 | ab |
| Desacetoxyvindoline | 873307.73 ± 99443.72 | ab | 772870.24 ± 49877.56 | a | 1031084.28 ± 66680.13 | ab | 1232873.85 ± 142466.49 | b | 1006326.28 ± 22033.45 | ab | 1060440.89 ± 46672.71 | ab | 1021962.40 ± 77824.72 | ab |
| Deacetylvindoline | 282304.45 ± 36056.18 | a | 199391.47 ± 19902.41 | a | 290925.18 ± 31137.29 | a | 313968.47 ± 38603.71 | a | 283389.48 ± 6843.87 | a | 275346.99 ± 10809.87 | a | 251926.67 ± 14220.10 | a |
| Vindoline | 11412219.14 ± 1402156.05 | a | 9818761.90 ± 690921.49 | a | 14008785.58 ± 1136842.77 | a | 14299655.06 ± 1534231.47 | a | 12189220.94 ± 267256.00 | a | 13787601.18 ± 529149.77 | a | 12030213.36 ± 801941.91 | a |
| Anhydrovinblastine | 305391.19 ± 33897.09 | ab | 162595.95 ± 25510.34 | a | 599733.78 ± 119951.26 | b | 446950.70 ± 99067.32 | ab | 407433.11 ± 35961.87 | ab | 595111.47 ± 66088.50 | b | 460606.97 ± 72688.60 | ab |
| Vinblastine | 245242.70 ± 27857.46 | ab | 123628.04 ± 12910.92 | a | 275905.49 ± 45990.33 | ab | 289032.11 ± 42902.01 | ab | 298203.22 ± 22738.05 | b | 294603.35 ± 27493.79 | b | 270721.22 ± 54741.81 | ab |
| Vincristine | 129534.57 ± 14596.21 | ab | 102900.24 ± 10925.07 | a | 159258.96 ± 14970.01 | ab | 144193.73 ± 19649.85 | ab | 147159.05 ± 7818.60 | ab | 170893.80 ± 10760.00 | b | 136680.70 ± 12330.71 | ab |
| Desacetoxyvindorosine | 11547.06 ± 1402.21 | a | 13557.14 ± 658.16 | a | 14863.69 ± 1833.83 | ab | 21372.32 ± 2339.35 | b | 15057.58 ± 928.01 | ab | 14106.74 ± 864.10 | a | 14629.20 ± 1225.25 | a |
| Desacetylvindorosine | 17564.11 ± 2199.54 | a | 11931.95 ± 1376.64 | a | 15365.79 ± 1212.46 | a | 16420.60 ± 1459.65 | a | 16468.11 ± 302.60 | a | 14751.65 ± 587.90 | a | 13622.95 ± 584.91 | a |
| Demethoxyvindoline = vindorosine | 591352.75 ± 92196.44 | ab | 459187.72 ± 46216.12 | a | 753659.92 ± 73507.83 | ab | 838896.04 ± 87617.59 | b | 630500.96 ± 22195.72 | ab | 704585.27 ± 35178.09 | ab | 639932.25 ± 72269.23 | ab |
| Vincadifformine | 909.06 ± 218.94 | a | 782.79 ± 235.22 | ac | 3824.19 ± 227.83 | b | 2455.34 ± 453.71 | bc | 3630.81 ± 558.72 | bd | 3294.33 ± 468.95 | bd | 1719.91 ± 458.28 | ac |
| 16‑Hydroxyvincadifformine | 3740.93 ± 557.65 | a | 19104.21 ± 1571.42 | bc | 26752.85 ± 1721.56 | cd | 14668.58 ± 1947.46 | b | 12185.37 ± 916.35 | b | 32663.51 ± 2301.73 | d | 19068.63 ± 2712.30 | bc |
| Minovincinine | 549.12 ± 131.16 | a | 464.20 ± 49.99 | a | 1432.51 ± 142.17 | bc | 1541.26 ± 79.89 | b | 1242.01 ± 100.75 | bc | 1296.04 ± 84.49 | bc | 1012.06 ± 138.30 | c |
| 19-hydroxytabersonine | 5754.82 ± 1216.82 | ab | 4104.39 ± 556.15 | a | 9421.75 ± 1183.57 | b | 9258.87 ± 770.72 | b | 7286.51 ± 721.66 | ab | 8179.48 ± 688.02 | ab | 8717.72 ± 1125.51 | b |
| 19-O-acetyltabersonine | 64850.36 ± 7959.88 | a | 99771.33 ± 5864.43 | ace | 175342.85 ± 5437.02 | bcd | 138808.79 ± 10829.95 | ce | 209987.89 ± 6073.83 | d | 180903.49 ± 12961.44 | bd | 114192.10 ± 7599.20 | e |
| 16-hydroxy-19-O-acetyltabersonine | 67.81 ± 32.99 | a | 14.01 ± 7.35 | a | 35.72 ± 21.12 | a | 211.52 ± 34.48 | b | 59.06 ± 26.21 | a | 43.88 ± 25.56 | a | 108.49 ± 45.56 | ab |
| 16-hydroxylochnericine | 80527.82 ± 9328.14 | a | 156266.70 ± 12522.67 | ab | 787393.21 ± 14979.19 | c | 248122.97 ± 5842.84 | ab | 915934.04 ± 57154.44 | c | 771615.35 ± 73858.75 | c | 260917.60 ± 31555.44 | b |
| 16-hydroxylochnericine glucoside | 12595.60 ± 2218.97 | a | 40952.93 ± 5451.75 | a | 339628.02 ± 19586.33 | b | 61257.70 ± 8489.74 | a | 349901.85 ± 22402.43 | b | 329299.04 ± 18942.78 | b | 69562.16 ± 5871.19 | a |
| Hörhammericine | 6585.21 ± 1179.33 | a | 8275.07 ± 1190.44 | a | 25026.61 ± 2402.81 | b | 9944.88 ± 742.43 | a | 9224.02 ± 712.22 | a | 31158.05 ± 3187.54 | b | 27232.76 ± 3771.75 | b |
| 16-hydroxyhörhammericine | 143.34 ± 17.26 | a | 418.33 ± 44.04 | a | 3662.33 ± 447.86 | b | 463.16 ± 238.07 | a | 602.59 ± 66.16 | a | 6963.80 ± 789.78 | c | 6348.52 ± 927.50 | c |
| 16-methoxyhörhammericine | 7915.35 ± 1058.97 | a | 97351.04 ± 7232.23 | b | 132958.96 ± 13980.19 | bd | 16566.13 ± 1273.76 | a | 26288.89 ± 1448.74 | a | 177294.42 ± 16041.10 | cd | 191384.58 ± 19700.40 | c |
| Vandrikidine | 102280.39 ± 18475.32 | ab | 79259.74 ± 10682.92 | a | 144809.09 ± 20418.22 | ab | 156512.31 ± 16583.05 | b | 113597.73 ± 8984.76 | ab | 120366.81 ± 9606.25 | ab | 114911.85 ± 12721.81 | ab |

**Table S8** MIA levels upon combinatorial overexpression of *CrMYC2a^D126N^* and *ORCAs*.

|  | ***p35S::GUS* ± SEM** |  | ***p35S::CrMYC2a^D126N^* ± SEM** |  | ***p35S::CrMYC2a^D126N^/ORCA2* ± SEM** |  | ***p35S::CrMYC2a^D126N^/ORCA3* ± SEM** |  | ***p35S::CrMYC2a^D126N^/ORCA4* ± SEM** |  | ***p35S::CrMYC2a^D126N^/ORCA5* ± SEM** |  | ***p35S::CrMYC2a^D126N^/ORCA6* ± SEM** |  |
| --- | --- | --- | --- | --- | --- | --- | --- | --- | --- | --- | --- | --- | --- | --- |
| Loganin | 8626.83 ± 1039.10 | ac | 7584.28 ± 799.17 | ac | 12854.25 ± 791.31 | b | 11760.92 ± 454.11 | ab | 11271.50 ± 2170.71 | ab | 6117.48 ± 793.40 | c | 10601.46 ± 740.24 | abc |
| Secologanin | 78983.28 ± 4724.84 | a | 77621.66 ± 5088.83 | a | 89818.44 ± 4344.08 | a | 90499.88 ± 2547.67 | a | 85320.19 ± 4098.18 | a | 73663.00 ± 6000.27 | a | 75629.66 ± 2121.52 | a |
| Strictosidine | 2237933.30 ± 227357.62 | a | 4749196.11 ± 385463.70 | bd | 4841076.75 ± 220047.52 | bd | 6653028.42 ± 345263.64 | bc | 7252442.05 ± 1092499.36 | c | 3232112.01 ± 333184.30 | ad | 3566567.54 ± 371696.25 | ad |
| Strictosidine secologanoside | 64.91 ± 38.14 | a | 519.48 ± 219.62 | ab | 754.22 ± 290.80 | ab | 656.86 ± 229.69 | ab | 2739.79 ± 1262.53 | b | 81.39 ± 81.39 | a | 123.62 ± 61.47 | a |
| Unknown 2 (strictosidine aglycone isomer) | 93556.02 ± 10753.51 | a | 157727.33 ± 14473.97 | abc | 191299.49 ± 11714.30 | b | 216536.96 ± 15089.17 | b | 168298.82 ± 21825.60 | bc | 117386.26 ± 15017.37 | c | 138837.70 ± 11799.74 | c |
| Strictosidinic Acid | 98768.62 ± 20453.62 | a | 498913.63 ± 56348.38 | b | 672303.51 ± 59814.35 | b | 980259.26 ± 70420.12 | c | 1002079.78 ± 112055.50 | d | 423561.61 ± 54153.46 | b | 267124.42 ± 34853.44 | ab |
| Serpentine | 1630727.39 ± 393739.56 | a | 2138763.52 ± 441653.75 | a | 2444748.14 ± 458244.53 | a | 2856104.91 ± 557111.68 | a | 2876271.98 ± 666233.06 | a | 1290570.91 ± 480503.73 | a | 1463399.19 ± 126682.43 | a |
| Unknown MIA (serpentine isomer) | 552494.43 ± 106460.31 | ab | 647786.31 ± 117665.42 | ab | 900296.47 ± 131434.68 | a | 995167.63 ± 137181.30 | ab | 958369.30 ± 186734.28 | ab | 391519.79 ± 112458.42 | b | 612073.16 ± 43479.30 | ab |
| Geissoschizine | 1684870.16 ± 260410.12 | a | 4656122.57 ± 431585.56 | bc | 9501071.02 ± 560489.48 | c | 13470663.44 ± 374367.60 | d | 5290417.91 ± 711635.26 | d | 5429048.70 ± 697783.78 | bc | 5580237.01 ± 372427.90 | b |
| Unknown MIA (geissoschizine isomer 1) | 1085340.19 ± 168209.90 | a | 2760815.32 ± 249803.13 | b | 6089576.41 ± 349748.99 | c | 8926383.61 ± 231247.90 | d | 3321401.52 ± 495045.64 | b | 3549753.38 ± 476536.66 | b | 3568594.57 ± 237245.21 | b |
| Unknown MIA (geissoschizine isomer 2) | 42488.73 ± 10638.50 | a | 276082.65 ± 26894.19 | bcd | 360865.35 ± 39508.28 | b | 377381.19 ± 15620.14 | b | 233250.30 ± 34081.94 | cd | 135202.28 ± 15750.90 | ac | 247215.81 ± 8607.02 | d |
| Isositsirikine | 228517.06 ± 28439.04 | a | 403108.56 ± 42075.05 | ab | 579377.20 ± 37935.19 | bc | 575872.36 ± 37374.66 | bc | 651175.38 ± 74752.76 | c | 300657.94 ± 41243.21 | a | 361984.26 ± 25136.82 | a |
| Unknown 1 (isositsirikine isomer) | 374539.78 ± 58952.37 | a | 562563.80 ± 69036.18 | ab | 781996.45 ± 67027.10 | bc | 1005580.86 ± 97544.60 | c | 731807.25 ± 122872.58 | abc | 470878.67 ± 78838.86 | a | 517133.91 ± 20228.66 | a |
| Perivine | 85247.92 ± 13341.86 | a | 480932.83 ± 63318.36 | bc | 480935.77 ± 42850.18 | bc | 372181.94 ± 33514.28 | b | 656746.96 ± 76721.27 | c | 255415.21 ± 40294.36 | ab | 405010.52 ± 52917.02 | b |
| Akuammicine | 122022.54 ± 16925.89 | a | 937559.91 ± 137598.51 | b | 1201065.95 ± 81081.86 | bc | 773372.22 ± 37980.45 | bd | 1428134.23 ± 167892.85 | c | 412781.48 ± 50887.77 | ad | 818158.19 ± 113077.32 | bd |
| O-acetylstemmadenine | 129894.59 ± 16235.56 | a | 444404.01 ± 30554.41 | bc | 501219.93 ± 29178.61 | b | 402740.03 ± 15511.38 | bc | 428924.91 ± 60737.86 | bc | 226261.13 ± 26047.47 | ad | 319438.58 ± 39559.44 | cd |
| 16-hydroxytabersonine | 69239.25 ± 8369.88 | a | 643021.06 ± 88175.95 | b | 1962267.77 ± 59154.15 | c | 1531087.36 ± 43751.08 | c | 2746220.62 ± 109314.44 | d | 614433.51 ± 78918.72 | b | 792531.63 ± 75621.81 | b |
| 16-hydroxytabersonine glucoside | 7291.49 ± 963.53 | a | 81335.81 ± 7017.24 | bd | 148596.15 ± 10954.46 | cd | 109070.65 ± 4671.16 | d | 273654.33 ± 16100.37 | e | 65262.88 ± 9274.26 | b | 62013.47 ± 6610.35 | b |
| Catharanthine | 6264287.65 ± 858352.19 | ab | 6304859.74 ± 895297.28 | ab | 9679809.00 ± 758529.24 | a | 10121644.32 ± 757858.39 | a | 9348679.55 ± 1237803.54 | a | 4383851.24 ± 886201.96 | b | 6988862.90 ± 499854.43 | ab |
| Unknown MIA (catharanthine isomer 1) | 4337315.88 ± 600986.07 | a | 4502610.17 ± 534354.00 | a | 6042148.63 ± 667862.27 | a | 6285607.40 ± 559530.27 | a | 6068825.67 ± 933979.83 | a | 3497648.91 ± 615798.46 | a | 4418897.74 ± 257472.39 | a |
| Unknown MIA (catharanthine isomer 2) | 3071725.14 ± 484810.49 | a | 3632875.97 ± 459108.94 | a | 4584953.73 ± 395268.04 | a | 4738077.44 ± 372650.22 | a | 4815156.61 ± 673990.72 | a | 2625682.63 ± 524244.26 | a | 3478928.06 ± 163659.70 | a |
| Unknown MIA (catharanthine isomer 3) | 572257.89 ± 75777.88 | abc | 565097.84 ± 73502.17 | ac | 892989.29 ± 76012.39 | abc | 915786.34 ± 72507.02 | b | 827401.26 ± 107631.52 | ab | 466230.00 ± 73245.22 | c | 638512.98 ± 36420.57 | abc |
| Desacetoxyvindoline | 689357.99 ± 57824.76 | acd | 634965.06 ± 66318.19 | a | 1061816.11 ± 21547.91 | bd | 1120700.68 ± 21165.97 | b | 812195.56 ± 82269.64 | c | 535624.59 ± 33122.94 | a | 880164.06 ± 31677.99 | cd |
| Deacetylvindoline | 280988.98 ± 33955.18 | a | 278110.15 ± 23312.99 | a | 278318.03 ± 30193.71 | a | 282441.47 ± 14883.32 | a | 300738.03 ± 42002.99 | a | 150090.30 ± 24836.22 | b | 214372.52 ± 13842.51 | ab |
| Vindoline | 9776321.34 ± 993997.98 | ab | 10440781.30 ± 1364195.83 | ab | 13776099.96 ± 968623.88 | a | 14402922.48 ± 750354.76 | a | 12529975.05 ± 1555210.34 | a | 7104481.84 ± 961226.59 | b | 10650093.45 ± 527464.84 | ab |
| Anhydrovinblastine | 141512.73 ± 28527.72 | ab | 212350.25 ± 29297.84 | ab | 369626.58 ± 84081.20 | a | 363866.10 ± 44151.80 | a | 320329.61 ± 77974.70 | ab | 120954.11 ± 31181.84 | b | 273287.35 ± 41493.00 | ab |
| Vinblastine | 57351.68 ± 11382.30 | a | 88021.62 ± 10303.26 | ab | 179765.81 ± 34719.63 | b | 174082.78 ± 15575.99 | b | 140687.98 ± 39458.90 | ab | 54637.09 ± 11729.77 | a | 130833.87 ± 20103.27 | ab |
| Vincristine | 41640.02 ± 5880.28 | ac | 52310.38 ± 6827.84 | abc | 88201.58 ± 10369.81 | b | 64349.26 ± 8266.85 | abc | 72704.41 ± 13275.37 | ab | 33714.36 ± 4243.72 | c | 57528.26 ± 4473.48 | abc |
| Desacetoxyvindorosine | 7585.41 ± 923.01 | ac | 6360.19 ± 777.65 | a | 18971.11 ± 2928.11 | b | 18705.45 ± 1504.89 | b | 11873.17 ± 1388.42 | ac | 7259.40 ± 885.11 | ac | 13714.07 ± 659.78 | bc |
| Desacetylvindorosine | 15199.29 ± 1781.37 | ab | 17362.99 ± 1992.51 | a | 16882.31 ± 1003.73 | a | 18259.69 ± 689.93 | a | 17619.11 ± 1832.13 | a | 10076.15 ± 1556.73 | b | 14086.73 ± 605.21 | ab |
| Demethoxyvindoline = vindorosine | 386198.50 ± 53653.51 | ac | 408797.04 ± 65847.67 | ac | 642038.32 ± 47690.78 | ab | 694586.44 ± 47929.86 | b | 542592.84 ± 72814.95 | ab | 277838.57 ± 45551.37 | c | 481896.69 ± 29774.34 | ac |
| Vincadifformine | 247.03 ± 49.40 | a | 5889.37 ± 1437.46 | a | 212848.32 ± 10063.38 | b | 226325.24 ± 15431.19 | b | 199138.82 ± 27974.21 | b | 45207.07 ± 9905.86 | a | 32964.75 ± 3702.96 | a |
| 16‑Hydroxyvincadifformine | 3500.91 ± 876.26 | a | 22256.18 ± 3298.01 | ae | 340014.47 ± 13910.91 | b | 379815.11 ± 22893.51 | b | 254677.20 ± 27322.21 | c | 108279.18 ± 17988.03 | d | 95435.29 ± 13339.80 | de |
| Minovincinine | 627.33 ± 167.81 | a | 807.99 ± 152.29 | a | 24469.84 ± 5947.05 | b | 34428.72 ± 3973.01 | c | 10253.38 ± 906.42 | a | 2875.48 ± 648.71 | a | 1380.17 ± 107.00 | a |
| 19-hydroxytabersonine | 2099.44 ± 401.36 | a | 2883.45 ± 389.09 | a | 119314.69 ± 13517.13 | b | 377209.41 ± 24740.83 | b | 104313.19 ± 12347.27 | a | 23898.89 ± 8481.32 | a | 4323.11 ± 401.31 | a |
| 19-O-acetyltabersonine | 55398.90 ± 4662.24 | a | 220765.33 ± 19790.28 | b | 452668.22 ± 9803.61 | cd | 348068.34 ± 12519.35 | bc | 554879.55 ± 80713.83 | d | 205184.02 ± 17613.09 | b | 227463.02 ± 29554.85 | b |
| 16-hydroxy-19-O-acetyltabersonine | 28.50 ± 8.90 | a | 19.30 ± 11.68 | a | 7504.69 ± 815.30 | a | 131981.66 ± 18961.02 | b | 7292.73 ± 1029.33 | a | 4189.18 ± 1508.80 | a | 49.28 ± 25.15 | a |
| 16-hydroxylochnericine | 52510.55 ± 6388.69 | a | 183475.69 ± 21187.12 | b | 463941.53 ± 18700.75 | c | 287130.22 ± 11521.42 | bc | 751298.45 ± 106106.81 | d | 141498.88 ± 19829.92 | ab | 141265.42 ± 9050.20 | ab |
| 16-hydroxylochnericine glucoside | 8980.47 ± 1612.25 | a | 106775.57 ± 21589.15 | bc | 95786.88 ± 12130.77 | bc | 58830.14 ± 1666.06 | ac | 181341.65 ± 25487.06 | d | 31050.31 ± 4324.36 | a | 20284.19 ± 1913.02 | a |
| Hörhammericine | 4724.77 ± 996.81 | a | 5992.97 ± 915.68 | a | 278470.00 ± 38042.54 | bc | 374346.28 ± 39153.63 | b | 185240.87 ± 36359.61 | c | 63542.74 ± 28403.72 | a | 6372.98 ± 543.77 | a |
| 16-hydroxyhörhammericine | 86.72 ± 48.60 | a | 193.00 ± 18.89 | a | 10580.30 ± 1703.02 | b | 26707.28 ± 1398.64 | c | 24552.49 ± 3524.69 | c | 3244.90 ± 986.72 | ab | 174.94 ± 36.97 | a |
| 16-methoxyhörhammericine | 6127.80 ± 1025.41 | a | 14570.47 ± 1812.75 | a | 1455135.43 ± 58409.57 | b | 1768828.83 ± 60800.49 | b | 1476769.83 ± 190025.82 | b | 628559.98 ± 120788.73 | c | 87278.84 ± 15027.98 | a |
| Vandrikidine | 58516.82 ± 8298.02 | a | 61809.06 ± 8650.40 | a | 286373.73 ± 13438.47 | b | 627657.05 ± 36265.94 | c | 311488.93 ± 31407.43 | b | 114670.79 ± 20833.02 | a | 72160.38 ± 6822.54 | a |

**Table S9** MIA levels upon overexpression of *MYB96/b*.

|  | ***p35S::GUS* ± SEM** |  | ***p35S::MYB96b* ± SEM** |  | ***p35S::MYB96* ± SEM** |  | ***p35S::MYB96/96b* ± SEM** |  |
| --- | --- | --- | --- | --- | --- | --- | --- | --- |
| Loganin | 18077.20 ± 1311.99 | a | 23202.36 ± 1551.57 | ab | 28192.15 ± 2957.71 | b | 21051.04 ± 1060.61 | ab |
| Secologanin | 166639.21 ± 11085.70 | a | 189229.95 ± 10593.10 | a | 187008.77 ± 15666.49 | a | 165322.53 ± 9280.56 | a |
| Strictosidine | 3249908.79 ± 32900.11 | ac | 2412417.01 ± 62155.70 | b | 2734654.41 ± 250541.41 | a | 3890892.24 ± 217153.69 | c |
| Strictosidine secologanoside | 3873.44 ± 358.60 | a | 110.62 ± 110.62 | b | 1006.49 ± 912.46 | b | 280.40 ± 280.40 | b |
| Unknown 2 (strictosidine aglycone isomer) | 95098.64 ± 9985.24 | a | 110953.76 ± 5712.12 | a | 95959.85 ± 9104.73 | a | 116455.19 ± 6188.85 | a |
| Strictosidinic Acid | 319097.32 ± 30435.43 | ab | 295479.71 ± 16263.51 | a | 272663.15 ± 37572.91 | a | 429114.31 ± 12376.90 | b |
| Serpentine | 2934652.60 ± 454437.97 | a | 2777297.91 ± 167880.75 | a | 3253367.89 ± 513643.97 | a | 3628148.31 ± 209836.09 | a |
| Unknown MIA (serpentine isomer) | 699111.61 ± 54890.41 | a | 667555.96 ± 41418.47 | a | 852078.76 ± 117001.48 | a | 1203419.96 ± 67104.93 | b |
| Geissoschizine | 1958540.56 ± 300324.84 | a | 2108506.58 ± 124252.41 | a | 1774373.95 ± 219955.12 | a | 1384011.68 ± 59339.72 | a |
| Unknown MIA (geissoschizine isomer 1) | 3517665.90 ± 495131.29 | a | 4181376.41 ± 278217.33 | a | 3440154.48 ± 390808.37 | a | 3122352.57 ± 112133.71 | a |
| Unknown MIA (geissoschizine isomer 2) | 94789.54 ± 12983.37 | a | 99653.36 ± 6191.92 | a | 82520.02 ± 12669.74 | a | 224065.05 ± 15165.13 | b |
| Unknown 1 (isositsirikine isomer) | 633621.94 ± 71704.80 | a | 633009.96 ± 27457.56 | a | 623066.31 ± 64597.28 | a | 658132.61 ± 43777.80 | a |
| Isositsirikine | 416370.09 ± 33899.53 | a | 394503.59 ± 19858.68 | a | 407085.93 ± 49096.73 | a | 540091.67 ± 29580.98 | a |
| Perivine | 143216.57 ± 6677.20 | a | 156338.91 ± 8522.00 | ab | 208846.44 ± 16544.12 | b | 301029.88 ± 17796.69 | c |
| Akuammicine | 243714.10 ± 14999.68 | a | 218826.92 ± 13563.25 | a | 275274.17 ± 32682.01 | a | 418342.16 ± 13918.86 | b |
| O-acetylstemmadenine | 256203.17 ± 9275.95 | a | 259941.53 ± 13593.20 | a | 239481.60 ± 12421.14 | a | 310548.87 ± 9331.95 | b |
| 16-hydroxytabersonine | 204610.96 ± 6092.51 | a | 228663.65 ± 16080.62 | a | 213691.18 ± 22233.43 | a | 373824.82 ± 13240.55 | b |
| 16-hydroxytabersonine glucoside | 12224.51 ± 3589.46 | a | 10193.54 ± 1107.57 | a | 7815.83 ± 768.46 | a | 25295.28 ± 2143.12 | b |
| Catharanthine | 9523886.58 ± 869603.54 | a | 9739259.01 ± 532742.66 | a | 10348636.85 ± 1029051.96 | a | 14733651.38 ± 570844.09 | b |
| Unknown MIA (catharanthine isomer 1) | 7153472.93 ± 456075.95 | a | 7302683.07 ± 500435.51 | a | 7760883.18 ± 763604.39 | a | 10508131.55 ± 751941.65 | b |
| Unknown MIA (catharanthine isomer 2) | 4440041.00 ± 323906.98 | a | 4409152.83 ± 302083.35 | a | 4809815.51 ± 470118.48 | a | 6787739.83 ± 427093.95 | b |
| Unknown MIA (catharanthine isomer 3) | 707826.50 ± 62826.83 | a | 722843.61 ± 48621.69 | a | 805858.79 ± 99615.49 | a | 1162660.09 ± 59521.33 | b |
| Desacetoxyvindoline | 761815.28 ± 44265.57 | a | 731145.46 ± 35310.95 | a | 728638.68 ± 43966.21 | a | 827125.88 ± 23511.19 | a |
| Deacetylvindoline | 349264.72 ± 26595.05 | a | 365332.19 ± 15360.14 | a | 363585.21 ± 37027.74 | a | 448500.41 ± 22624.85 | a |
| Vindoline | 10803870.27 ± 502242.35 | a | 10794229.84 ± 786105.05 | a | 9906909.68 ± 530268.23 | a | 10963867.33 ± 654783.22 | a |
| Anhydrovinblastine | 533421.25 ± 43732.49 | a | 336824.47 ± 41900.30 | a | 280294.74 ± 71595.24 | a | 482590.01 ± 78798.09 | a |
| Vinblastine | 119772.46 ± 10921.38 | a | 87389.33 ± 13087.25 | a | 61875.87 ± 16058.29 | a | 100642.06 ± 20718.21 | a |
| Vincristine | 112736.71 ± 8163.45 | a | 89011.88 ± 7574.91 | a | 83684.92 ± 15750.46 | a | 136641.02 ± 17197.14 | a |
| Desacetoxyvindorosine | 9109.75 ± 541.10 | a | 9099.32 ± 761.80 | a | 10478.56 ± 1119.57 | a | 10672.73 ± 149.04 | a |
| Desacetylvindorosine | 22891.62 ± 1651.31 | a | 24970.16 ± 1896.91 | ab | 33263.04 ± 3260.62 | bc | 36335.65 ± 1254.43 | c |
| Demethoxyvindoline = vindorosine | 623608.99 ± 60975.46 | a | 679723.22 ± 51393.01 | ab | 747367.33 ± 93818.53 | ab | 907585.89 ± 27750.51 | b |
| Vincadifformine | 3970.77 ± 906.62 | a | 4587.13 ± 557.40 | a | 4197.33 ± 611.45 | a | 8178.08 ± 466.37 | b |
| 16‑Hydroxyvincadifformine | 9319.24 ± 1475.38 | a | 10636.60 ± 716.42 | a | 10536.19 ± 1930.16 | a | 11137.91 ± 685.78 | a |
| Minovincinine | 2140.65 ± 168.68 | a | 1876.23 ± 244.38 | a | 2591.28 ± 419.49 | a | 4290.63 ± 301.01 | b |
| 19-hydroxytabersonine | 43433.28 ± 2564.97 | a | 53601.84 ± 5291.97 | a | 53920.66 ± 4842.04 | a | 71636.23 ± 1804.03 | b |
| 19-O-acetyltabersonine | 83064.17 ± 4142.76 | a | 91102.16 ± 5708.77 | a | 93903.91 ± 8907.29 | a | 138903.70 ± 7777.72 | b |
| 16-hydroxy-19-O-acetyltabersonine | 287.33 ± 111.64 | a | 258.19 ± 96.52 | a | 380.19 ± 158.87 | a | 154.46 ± 41.51 | a |
| 16-hydroxylochnericine | 239563.89 ± 24918.66 | a | 284786.38 ± 14148.97 | ab | 313035.09 ± 37059.05 | ab | 392134.46 ± 24480.09 | b |
| 16-hydroxylochnericine glucoside | 11464.25 ± 8059.56 | a | 8321.25 ± 1827.04 | a | 7232.79 ± 1781.23 | a | 35186.17 ± 5258.00 | b |
| Hörhammericine | 37968.54 ± 2655.73 | a | 47364.79 ± 2702.62 | ab | 46619.19 ± 5077.90 | a | 60762.98 ± 2182.34 | b |
| 16-hydroxyhörhammericine | 142.24 ± 25.16 | a | 70.10 ± 24.37 | a | 133.20 ± 16.90 | a | 124.92 ± 46.23 | a |
| 16-methoxyhörhammericine | 9054.03 ± 114.64 | a | 8936.00 ± 402.63 | a | 8733.89 ± 1095.19 | a | 19182.45 ± 1781.51 | b |
| Vandrikidine | 154225.34 ± 14757.24 | a | 170333.52 ± 12008.57 | a | 178765.73 ± 21251.04 | ab | 232296.58 ± 5051.69 | b |

**Table S10** MIA levels upon overexpression of *MYB96/b, CrMYC2a^D126N^* and *ORCA4*.

|  | ***p35S::GUS* ± SEM** |  | ***p35S::CrMYC2a^D126N^* ± SEM** |  | ***p35S::ORCA4***  **± SEM** |  | ***p35S::MYB96* ± SEM** |  | ***p35S::CrMYC2a^D126N^/ORCA4* ± SEM** |  | ***p35S::ORCA4/MYB96* ± SEM** |  | ***p35S::CrMYC2a^D126N^/MYB96* ± SEM** |  | ***p35S::CrMYC2a^D126N^/***  ***ORCA4/MYB96* ± SEM** |  |
| --- | --- | --- | --- | --- | --- | --- | --- | --- | --- | --- | --- | --- | --- | --- | --- | --- |
| Loganin | 9332.21 ± 670.24 | a | 9145.02 ± 1093.29 | a | 9749.58 ± 1349.63 | a | 11278.43 ± 1875.02 | a | 12390.85 ± 627.01 | a | 10389.86 ± 1093.90 | a | 8657.14 ± 629.17 | a | 11832.32 ± 496.03 | a |
| Secologanin | 80576.30 ± 1464.34 | a | 76038.85 ± 2902.62 | a | 76294.12 ± 2023.22 | a | 86004.92 ± 6046.99 | a | 82294.31 ± 2112.81 | a | 76911.70 ± 3463.54 | a | 81200.08 ± 5770.51 | a | 88012.94 ± 1482.64 | a |
| Strictosidine | 1878964.65 ± 32854.65 | a | 3681759.40 ± 390608.12 | b | 3589440.75 ± 367966.61 | b | 2118946.68 ± 340878.91 | ac | 5704397.26 ± 272573.10 | d | 3681482.17 ± 216910.28 | b | 3241250.30 ± 230225.64 | bc | 4298522.19 ± 82989.58 | b |
| Strictosidine secologanoside | 21.18 ± 21.18 | a | 478.54 ± 141.26 | ac | 155.79 ± 65.71 | a | 131.64 ± 89.13 | a | 3221.50 ± 467.63 | b | 266.15 ± 114.89 | a | 167.66 ± 166.01 | a | 1304.77 ± 296.92 | c |
| Unknown 2 (strictosidine aglycone isomer) | 80887.16 ± 2351.80 | a | 98933.16 ± 10800.57 | a | 80384.16 ± 6774.79 | a | 85617.32 ± 8941.26 | a | 109306.10 ± 2558.56 | a | 103188.24 ± 3899.31 | a | 92772.02 ± 5563.26 | a | 101132.05 ± 2185.94 | a |
| Strictosidinic Acid | 74892.04 ± 18046.67 | a | 151651.41 ± 18447.75 | ad | 448524.22 ± 53908.12 | b | 59766.04 ± 7006.73 | a | 730982.85 ± 41114.00 | c | 283137.79 ± 6982.35 | d | 178325.46 ± 20434.58 | ad | 480095.82 ± 38582.79 | b |
| Serpentine | 1773919.33 ± 179007.37 | a | 1797351.47 ± 355495.59 | a | 1411218.50 ± 242832.40 | a | 1945287.83 ± 312072.04 | a | 2116516.31 ± 107442.18 | a | 1944529.14 ± 212919.58 | a | 1680917.92 ± 254719.41 | a | 1922687.75 ± 168717.65 | a |
| Unknown MIA (serpentine isomer) | 543850.31 ± 37936.69 | a | 512655.45 ± 96116.74 | a | 479470.62 ± 79208.50 | a | 558725.21 ± 89192.89 | a | 781436.21 ± 29652.46 | a | 772071.87 ± 79957.80 | a | 521913.71 ± 62477.14 | a | 651391.30 ± 40765.34 | a |
| Geissoschizine | 935438.08 ± 50202.60 | a | 2704771.21 ± 400087.77 | be | 1912838.73 ± 261108.63 | bc | 1589684.30 ± 250103.99 | c | 4414049.70 ± 255513.88 | d | 1951477.70 ± 60394.70 | abc | 2001953.29 ± 66141.24 | bc | 3157425.18 ± 113417.51 | e |
| Unknown MIA (geissoschizine isomer 1) | 570123.83 ± 32701.96 | a | 1571415.11 ± 229194.25 | bde | 1175808.07 ± 176902.25 | bd | 958110.20 ± 141171.70 | ad | 2593610.43 ± 132045.21 | c | 1269091.18 ± 36405.24 | d | 1132908.32 ± 39506.84 | ad | 1878428.23 ± 76769.31 | e |
| Unknown MIA (geissoschizine isomer 2) | 61498.79 ± 8375.42 | a | 198383.12 ± 39106.71 | bc | 90652.22 ± 18901.02 | ad | 101180.94 ± 25685.36 | acd | 246594.52 ± 13297.48 | b | 104928.70 ± 19935.43 | acd | 203999.14 ± 20899.43 | bd | 187425.55 ± 20141.93 | bd |
| Isositsirikine | 195276.61 ± 2013.75 | ab | 289545.92 ± 32738.82 | bd | 246762.85 ± 21950.17 | bd | 217616.68 ± 30006.38 | b | 500285.21 ± 24540.06 | c | 306755.45 ± 14312.96 | bd | 263482.34 ± 15237.61 | bd | 337763.56 ± 7397.93 | d |
| Unknown 1 (isositsirikine isomer) | 304514.64 ± 6690.27 | a | 370906.49 ± 49861.56 | a | 311485.59 ± 30595.22 | a | 319953.69 ± 40361.99 | a | 528541.46 ± 12103.56 | b | 407747.41 ± 20396.93 | ab | 338506.06 ± 21948.50 | a | 408633.12 ± 9314.01 | ab |
| Perivine | 77500.61 ± 1958.16 | a | 329998.96 ± 57036.39 | bcdf | 99439.63 ± 9469.55 | ad | 114359.03 ± 21412.43 | ad | 303961.13 ± 20030.12 | cdf | 210373.57 ± 10500.41 | d | 496394.96 ± 48715.35 | e | 398180.07 ± 42894.45 | ef |
| Akuammicine | 137373.37 ± 5597.61 | a | 727458.69 ± 145196.32 | bdef | 253547.93 ± 32132.97 | c | 183977.91 ± 34245.82 | c | 968382.34 ± 42986.83 | df | 448383.84 ± 31297.80 | ce | 709936.97 ± 69036.30 | ef | 983034.25 ± 77078.21 | f |
| O-acetylstemmadenine | 113816.67 ± 9641.36 | a | 502984.15 ± 62387.95 | bc | 193477.32 ± 7164.20 | a | 133395.35 ± 17818.22 | a | 556439.36 ± 30591.19 | b | 233108.65 ± 8624.69 | a | 408625.87 ± 33884.72 | c | 481489.19 ± 11337.95 | bc |
| 16-hydroxytabersonine | 80914.90 ± 5591.51 | a | 485874.77 ± 84793.03 | bc | 703212.60 ± 55317.69 | c | 112853.49 ± 22198.41 | a | 2379295.53 ± 122007.38 | d | 534157.38 ± 22011.97 | bc | 339230.59 ± 16354.39 | ab | 1567952.30 ± 81104.99 | e |
| 16-hydroxytabersonine glucoside | 11654.71 ± 1357.93 | a | 48498.27 ± 8816.57 | ab | 80475.90 ± 8055.34 | be | 15251.98 ± 4438.25 | a | 233168.69 ± 13661.28 | c | 35502.78 ± 4289.39 | a | 45033.90 ± 5703.39 | ae | 151113.92 ± 17760.97 | d |
| Catharanthine | 6586547.17 ± 240876.72 | ab | 6637131.39 ± 1164743.46 | ab | 6432821.83 ± 637496.81 | ab | 7027293.38 ± 981397.24 | ab | 9873252.58 ± 309436.29 | ab | 10011238.24 ± 981359.59 | a | 6389266.73 ± 679522.44 | b | 8123683.85 ± 645254.12 | ab |
| Unknown MIA (catharanthine isomer 1) | 4483285.77 ± 143565.56 | ab | 4175576.27 ± 498917.24 | a | 4019400.89 ± 314794.82 | a | 4282055.57 ± 458618.14 | a | 5415114.71 ± 172585.81 | ab | 5919705.95 ± 431916.12 | b | 4165164.60 ± 318379.37 | a | 4690857.72 ± 249518.96 | ab |
| Unknown MIA (catharanthine isomer 2) | 3146655.64 ± 93617.46 | ab | 3054348.24 ± 429748.00 | ab | 2777286.75 ± 282517.61 | a | 2949961.39 ± 359451.51 | a | 4019998.50 ± 112737.07 | ab | 4234888.43 ± 264612.26 | b | 3370115.46 ± 283195.76 | ab | 3626089.34 ± 121601.08 | ab |
| Unknown MIA (catharanthine isomer 3) | 589747.11 ± 24632.93 | ab | 517141.25 ± 67525.09 | a | 539712.11 ± 56473.30 | ab | 544340.56 ± 68291.73 | ab | 754205.43 ± 24179.39 | bc | 779315.73 ± 46399.93 | c | 526403.13 ± 49781.83 | a | 629164.36 ± 33344.37 | ac |
| Desacetoxyvindoline | 572496.85 ± 36787.88 | a | 676426.49 ± 108263.61 | a | 652723.69 ± 65240.11 | a | 730985.35 ± 118851.25 | a | 881040.14 ± 34219.94 | a | 716408.76 ± 72773.19 | a | 649538.89 ± 83286.48 | a | 881291.02 ± 36764.44 | a |
| Deacetylvindoline | 237696.83 ± 8784.70 | a | 209199.29 ± 24728.42 | a | 203152.92 ± 19678.85 | a | 253281.45 ± 38531.45 | a | 233532.30 ± 10075.73 | a | 269212.29 ± 20301.13 | a | 259884.80 ± 16518.21 | a | 255929.47 ± 11027.13 | a |
| Vindoline | 9062435.33 ± 413759.41 | a | 8875880.17 ± 1103042.00 | a | 8949539.10 ± 856662.39 | a | 9892297.95 ± 1319574.54 | a | 11190508.80 ± 498438.71 | a | 11137011.40 ± 1017832.04 | a | 9415932.01 ± 896999.74 | a | 10612136.84 ± 460549.93 | a |
| Anhydrovinblastine | 294946.45 ± 7367.57 | a | 330214.63 ± 107480.02 | a | 228997.91 ± 61666.26 | a | 291103.18 ± 71641.33 | a | 498324.27 ± 30687.94 | a | 352055.39 ± 84950.86 | a | 277617.52 ± 78475.24 | a | 281310.18 ± 35430.55 | a |
| Vinblastine | 196068.75 ± 14170.45 | a | 235935.28 ± 75339.01 | a | 145213.82 ± 36744.45 | a | 216357.71 ± 47476.90 | a | 339491.03 ± 28535.21 | a | 254708.50 ± 54303.09 | a | 160679.39 ± 28666.33 | a | 211517.27 ± 35913.34 | a |
| Vincristine | 48430.87 ± 1888.95 | a | 62219.03 ± 12082.38 | ab | 57187.14 ± 11583.32 | ab | 77487.66 ± 11752.22 | ab | 90556.22 ± 3607.04 | b | 73501.85 ± 6853.79 | ab | 65165.31 ± 5437.77 | ab | 74491.45 ± 7056.38 | ab |
| Desacetoxyvindorosine | 6826.85 ± 874.77 | a | 8079.31 ± 1429.56 | a | 8734.79 ± 899.74 | a | 10657.55 ± 1794.77 | abc | 15681.26 ± 1180.10 | bc | 10292.56 ± 776.07 | ac | 8070.30 ± 1287.38 | a | 14655.16 ± 681.84 | c |
| Desacetylvindorosine | 14049.80 ± 591.97 | a | 12911.79 ± 1346.70 | a | 11032.18 ± 1359.84 | a | 14699.43 ± 1910.78 | a | 13289.98 ± 167.77 | a | 14396.46 ± 652.41 | a | 15390.35 ± 1103.33 | a | 15997.52 ± 700.07 | a |
| Demethoxyvindoline = vindorosine | 445804.30 ± 30556.19 | a | 410520.96 ± 77696.18 | a | 403909.35 ± 50553.81 | a | 455758.62 ± 59245.16 | a | 586173.39 ± 37190.59 | a | 561708.00 ± 64070.61 | a | 416085.62 ± 63025.83 | a | 526572.38 ± 49473.15 | a |
| Vincadifformine | 426.83 ± 141.27 | a | 3294.07 ± 936.88 | a | 1609.49 ± 379.28 | a | 367.00 ± 118.28 | a | 161717.94 ± 26586.34 | b | 1824.37 ± 481.91 | a | 3350.49 ± 1811.31 | a | 44784.72 ± 3955.88 | a |
| 16‑Hydroxyvincadifformine | 2523.05 ± 306.05 | a | 9292.96 ± 2029.49 | a | 4864.29 ± 806.95 | a | 4310.45 ± 852.99 | a | 171954.20 ± 16131.49 | b | 6652.05 ± 392.58 | a | 7063.49 ± 490.41 | a | 69848.73 ± 6492.19 | c |
| Minovincinine | 494.47 ± 111.91 | ac | 612.41 ± 183.35 | ac | 641.74 ± 167.28 | ac | 391.15 ± 60.25 | ac | 9238.83 ± 845.70 | b | 1098.05 ± 126.28 | ac | 378.30 ± 54.67 | a | 1892.15 ± 102.36 | c |
| 19-hydroxytabersonine | 3202.36 ± 331.52 | a | 2959.23 ± 657.84 | a | 2635.34 ± 324.46 | a | 2573.33 ± 446.13 | a | 102841.04 ± 15335.70 | b | 4951.32 ± 717.16 | a | 2535.97 ± 315.08 | a | 39611.25 ± 5961.26 | c |
| 19-O-acetyltabersonine | 52096.51 ± 432.48 | a | 163178.45 ± 19004.81 | b | 123485.41 ± 9155.91 | ab | 66025.44 ± 9462.93 | a | 516658.30 ± 37899.16 | c | 148414.08 ± 5677.54 | b | 145557.18 ± 9373.27 | b | 397300.75 ± 7568.17 | d |
| 16-hydroxy-19-O-acetyltabersonine | 37.19 ± 14.46 | a | 46.29 ± 26.90 | a | 18.59 ± 18.01 | a | 59.04 ± 14.46 | a | 713.54 ± 103.70 | b | 38.19 ± 16.85 | a | 35.81 ± 25.18 | a | 99.71 ± 34.43 | a |
| 16-hydroxylochnericine | 60875.77 ± 2312.96 | a | 118921.38 ± 17685.54 | a | 347874.15 ± 40717.38 | b | 66726.99 ± 9524.43 | a | 620212.14 ± 16731.42 | c | 403158.09 ± 10763.91 | b | 119378.89 ± 7251.55 | a | 546424.58 ± 37269.60 | c |
| 16-hydroxylochnericine glucoside | 9703.16 ± 1610.01 | a | 20070.68 ± 4179.77 | a | 126704.48 ± 13047.73 | b | 13469.81 ± 4451.24 | a | 145039.15 ± 9232.74 | b | 83588.78 ± 5371.03 | c | 30701.25 ± 5026.19 | a | 114799.31 ± 16200.68 | bc |
| Hörhammericine | 5649.26 ± 488.75 | a | 5168.97 ± 1037.97 | a | 4944.01 ± 793.23 | a | 4522.21 ± 628.76 | a | 160223.61 ± 16721.94 | b | 7298.05 ± 414.32 | a | 5552.77 ± 844.45 | a | 112375.38 ± 18507.65 | c |
| 16-hydroxyhörhammericine | 205.76 ± 16.17 | a | 94.76 ± 16.01 | a | 151.71 ± 41.33 | a | 106.23 ± 35.07 | a | 12716.05 ± 833.07 | b | 248.22 ± 30.72 | a | 128.68 ± 44.84 | a | 9024.61 ± 1475.29 | c |
| 16-methoxyhörhammericine | 5669.46 ± 299.63 | a | 8416.25 ± 1018.94 | a | 8027.10 ± 969.87 | a | 5994.55 ± 1319.74 | a | 1278211.75 ± 148975.94 | b | 14673.34 ± 856.02 | a | 14074.94 ± 2002.04 | a | 600138.97 ± 80691.00 | c |
| Vandrikidine | 66372.97 ± 3644.32 | a | 59197.58 ± 10499.71 | a | 59401.41 ± 5904.84 | a | 64826.75 ± 9454.28 | a | 293123.07 ± 28000.55 | b | 94476.22 ± 4949.51 | a | 58706.73 ± 5826.50 | a | 138841.73 ± 8077.63 | c |
